# Classification of electrophysiological and morphological types in mouse visual cortex

**DOI:** 10.1101/368456

**Authors:** Nathan W. Gouwens, Staci A. Sorensen, Jim Berg, Changkyu Lee, Tim Jarsky, Jonathan Ting, Susan M. Sunkin, David Feng, Costas Anastassiou, Eliza Barkan, Kris Bickley, Nicole Blesie, Thomas Braun, Krissy Brouner, Agata Budzillo, Shiella Caldejon, Tamara Casper, Dan Casteli, Peter Chong, Kirsten Crichton, Christine Cuhaciyan, L. Daigle Tanya, Rachel Dalley, Nick Dee, Tsega Desta, Samuel Dingman, Alyse Doperalski, Nadezhda Dotson, Tom Egdorf, Michael Fisher, Rebecca A. de Frates, Emma Garren, Marissa Garwood, Amanda Gary, Nathalie Gaudreault, Keith Godfrey, Melissa Gorham, Hong Gu, Caroline Habel, Kristen Hadley, James Harrington, Julie Harris, Alex Henry, DiJon Hill, Sam Josephsen, Sara Kebede, Lisa Kim, Matthew Kroll, Brian Lee, Tracy Lemon, Xiaoxiao Liu, Brian Long, Rusty Mann, Medea McGraw, Stefan Mihalas, Alice Mukora, Gabe J. Murphy, Lindsay Ng, Kiet Ngo, Thuc Nghi Nguyen, Philip R. Nicovich, Aaron Oldre, Daniel Park, Sheana Parry, Jed Perkins, Lydia Potekhina, David Reid, Miranda Robertson, David Sandman, Martin Schroedter, Cliff Slaughterbeck, Gilberto Soler-Llavina, Josef Sulc, Aaron Szafer, Bosiljka Tasic, Naz Taskin, Corinne Teeter, Nivretta Thatra, Herman Tung, Wayne Wakeman, Grace Williams, Rob Young, Zhi Zhou, Colin Farrell, Hanchuan Peng, Michael J. Hawrylycz, Ed Lein, Lydia Ng, Anton Arkhipov, Amy Bernard, John W. Phillips, Hongkui Zeng, Christof Koch

**Author notes:** These authors contributed equally to this work (Nathan W. Gouwens, Staci A. Sorensen, Jim Berg). Correspondence should be addressed to H.Z.

## Abstract

Understanding the diversity of cell types in the brain has been an enduring challenge and requires detailed characterization of individual neurons in multiple dimensions. To profile morpho-electric properties of mammalian neurons systematically, we established a single cell characterization pipeline using standardized patch clamp recordings in brain slices and biocytin-based neuronal reconstructions. We built a publicly-accessible online database, the Allen Cell Types Database, to display these data sets. Intrinsic physiological and morphological properties were measured from over 1,800 neurons from the adult laboratory mouse visual cortex. Quantitative features were used to classify neurons into distinct types using unsupervised methods. We establish a taxonomy of morphologically- and electrophysiologically-defined cell types for this region of cortex with 17 e-types and 35 m-types, as well as an initial correspondence with previously-defined transcriptomic cell types using the same transgenic mouse lines.

## INTRODUCTION

Neurons of the mammalian neocortex exhibit diverse physiological and morphological characteristics. Classifying these neurons into cell types, following Plato’s dictum to “carve nature at its joints,” provides a useful abstraction when investigating how these neurons interact in neocortical circuits^1^. The delineation of neocortical cell types has benefited from a wealth of studies detailing the molecular, morphological, and physiological properties of excitatory and inhibitory neurons, leading to the characterization of three major populations (i.e., Pvalb+, Sst+, and Htr3a+) of neocortical interneurons, each with distinct sub-classes, and excitatory neuron populations with features often linked to their laminar locations as well as the locations to which they project their axons^2–4^. Historically, though, direct comparisons of morpho-electric properties across studies have been constrained by differences in experimental protocols, analysis methods, and access to specific transgenic lines, limiting the construction of a comprehensive, systematic classification scheme. Recent studies performed at larger scales^5,6^, including the Blue Brain Project’s thorough classification of morpho-electric diversity in neonatal rat somatosensory cortex, have overcome many of these issues, using large, coherent data sets to identify principles of local connectivity and computation within a cortical column.

So far, morphological and physiological descriptions of cell types (including in these large-scale studies) have relied on expert annotation and categorization^7^. Such descriptions are valuable and can be consistent with statistical analyses of features^8^, but they can be limited by the number of criteria that can be simultaneously used to differentiate types. In addition, features used to distinguish types can be specific to the population being studied and may not be broadly applicable (or distinguishing) across all cortical neurons. Specific cell types have been characterized quantitatively^9–11^, but these studies frequently have used additional labels (like transgenically-driven reporter expression) to pre-define cell types. In addition, these approaches may successfully identify heterogeneity in features between specific populations under investigation, but these differences may not necessarily co-vary with other features or display discontinuities when examined in the context of a broader set of cortical cells— significant criteria for cell-type classification^1^.

Recent studies using single-cell transcriptomic characterization based on RNA-Seq techniques have performed unsupervised classification to generate a taxonomy of transcriptomic types in the mouse neocortex^12–14^. These approaches rely on data from thousands of neurons generated in a standardized manner. Here we have taken a similar approach to classifying morpho-electric properties of adult mouse visual cortical neurons using data collected from over 1,800 cells. We have characterized mouse visual cortex neurons with a uniform experimental protocol and developed an unsupervised classification scheme of cell types based on electrophysiology and morphology. Every aspect of the pipeline, from slice preparation, recording, and stimulation, to staining, imaging, 3D reconstructions, and mapping of cells to a reference atlas, employed highly standardized and quality-controlled methodology. A comprehensive set of transgenic mouse lines^14–17^ were used to ensure broad coverage of excitatory and inhibitory classes across all cortical layers, to enable selective targeting of rare cell populations, and to link our study with other experimental approaches, such as transcriptomic characterization. Unsupervised clustering methods map neurons of the adult mouse visual cortex to 17 distinct electrophysiological or e-types and 35 morphological or m-types. We examine correspondences between the electrophysiological and morphological types and use transgenic labels to establish preliminary links to major transcriptomic sub-classes and to specific transcriptomic types. All the experimental data and analysis tools are made available as a public resource as part of the Allen Cell Types Database and the Allen Software Development Kit.

## RESULTS

### Creating the in vitro single cell characterization platform

We set out to characterize the diversity of intrinsic electrophysiological and morphological properties of mouse visual cortical neurons by establishing a platform for generating a standardized data set (Fig. 1). We made whole-cell patch clamp recordings from neurons labeled by fluorescent proteins with an expression pattern determined by a Cre-based driver. We used a variety of driver lines to sample broadly across the cortical circuit as well as to target specific neuronal populations (Supplementary Figs. 1 and 2). The recording pipette contained biocytin to enable filling, staining, imaging, and three-dimensional reconstruction of neuronal morphologies. Each neuron was localized within the mouse reference atlas using serial block-face images generated during tissue slicing. Most cells (n=1,525) were from primary visual cortex (VISp); the remainder were in nearby higher visual areas. For morphological analysis, we reconstructed a subset of recorded neurons, selected based on fill quality and data coverage. All recorded cells underwent the same workflow (Fig. 1a), and only cells that met pre-defined quality control (QC) standards were included in the final data set, which consisted of 1,851 cells passing electrophysiology QC. Of these, 885 were spiny neurons, which we assume to be excitatory, and 966 were aspiny or sparsely spiny, which we assume to be inhibitory; this determination was made by examining the image of each cell (Methods). We reconstructed the morphologies (Methods) of 372 of those cells (199 spiny and 173 aspiny/sparsely spiny). Electrophysiological, imaging and morphology reconstruction data (when available) for each cell is freely accessible as part of the Allen Cell Types Database (http://celltypes.brain-map.org/) (Fig. 1b), including an interactive website as well as the opportunity to download all raw data.

**Figure 1:**
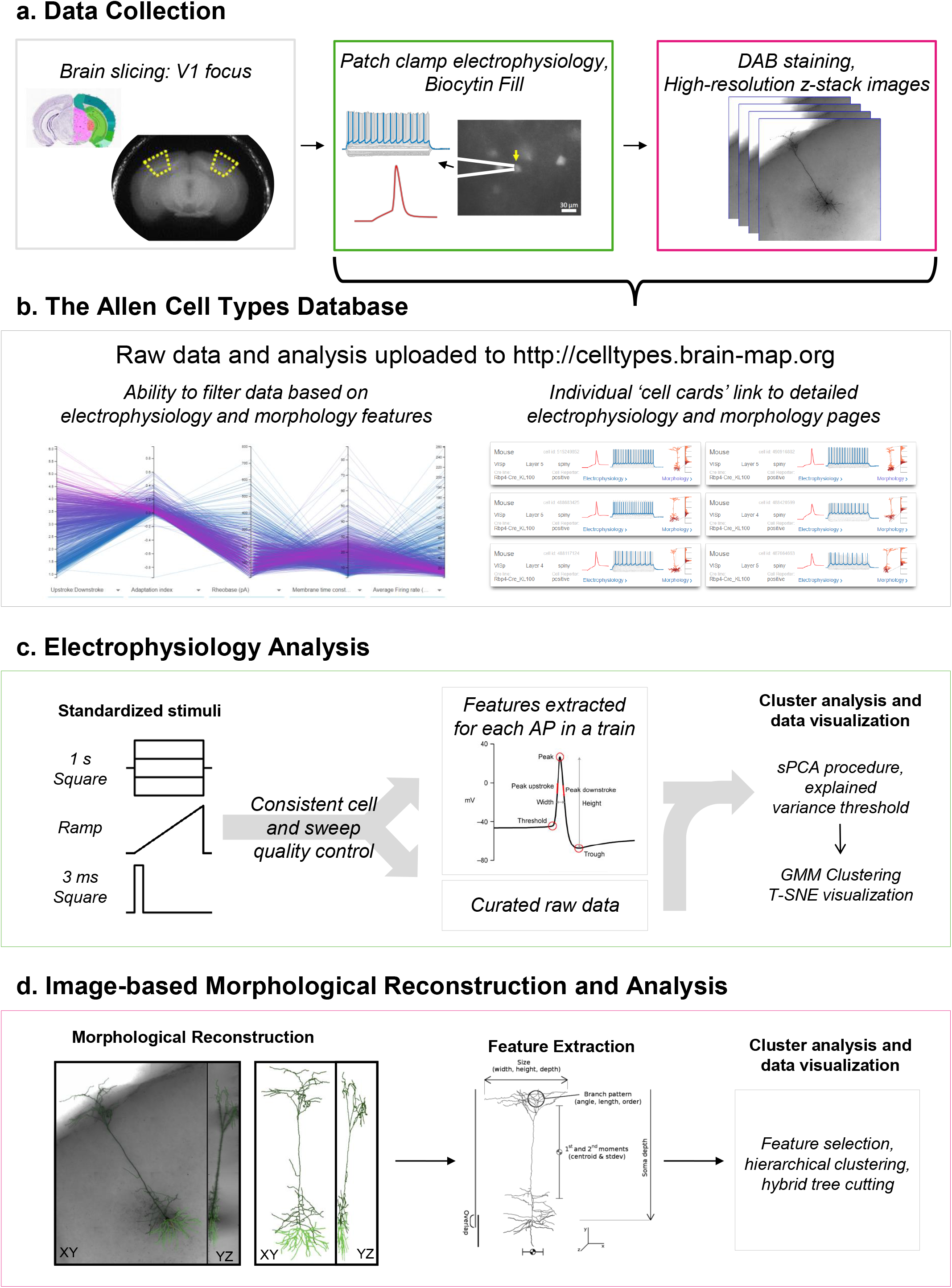
A pipeline to generate and analyze standardized morpho-electric data at scale. (**a**) An in vitro single cell characterization pipeline was established to generate standardized electrophysiological and morphological data from mouse cortical neurons. Mouse brains were imaged during vibratome sectioning to aid in cell localization to a common mouse reference atlas, Allen Mouse Common Coordinate Framework version 3 (CCF v3). Fluorescently labeled neurons from specific transgenic mouse lines were recorded by whole cell patch clamping to characterize each cell’s intrinsic electrical properties. During the electrophysiology recording, cells were filled with biocytin, then tissue slices were fixed, stained and mounted, and imaged in a high-resolution z stack. High quality cells were then manually reconstructed based on the z-stack images. (**b**) Electrophysiology, imaging, and morphology data and metadata for each cell are made freely accessible through the Allen Cell Types Database. An interactive user interface allows users to filter thousands of cells by electrophysiology and morphology features, then each cell has a specific page with detailed electrophysiology and morphology data, when available. (**c**) Each cell was stimulated with a standard electrophysiological stimulation paradigm, allowing for routine feature extraction and alignment of data traces from diverse cell types. Both raw data and series of features extracted from action potential trains underwent sparse principal component analysis followed by Gaussian mixture model fitting and clustering. (**d**) A subset of neurons were morphologically reconstructed followed by feature extraction, including branching and profile statistics. Neurons were clustered morphologically by hierarchical clustering followed by hybrid tree cutting.

### Electrophysiology classification

We characterized the intrinsic electrophysiological or e-properties with a standardized current-clamp protocol that contained a variety of stimuli, including square pulses, ramps, and noisy current injections (Fig. 1c, Supplementary Fig. 3). Our goal was to assess a diverse array of e-features while still enabling comparison across all cells.

We derived features from both membrane potential traces and specific characteristics of APs. For example, waveforms of the first APs evoked from a 3 ms current step, a 1 s current step, and a ramp stimulus were collected from each cell and aligned on the time of their thresholds (Fig. 2a). Responses to a series of hyperpolarizing current steps were also extracted and aligned (Fig. 2b). For each AP evoked by depolarizing current steps, a set of features was calculated (Methods). Since the number of APs evoked varied over two orders of magnitude across stimulus amplitudes and across cells, responses were compared by dividing the stimulus interval into 20 ms bins, calculating the average of the AP features within that bin, and interpolating values for bins without APs (Methods). An example of the threshold voltage feature is shown in Fig. 2c. The shape of the membrane potential trajectory during the interspike interval (ISI) also varied significantly across cells. To capture this, the trajectories were normalized to the same duration and calculated as a difference from the AP threshold voltage (Fig. 2d). Other features were collected in similar ways (Methods) forming 13 different subsets of data (Supplementary Table 1). Together, they represented multiple aspects of the electrophysiological responses that could be robustly extracted and compared across all the cells.

**Figure 2:**
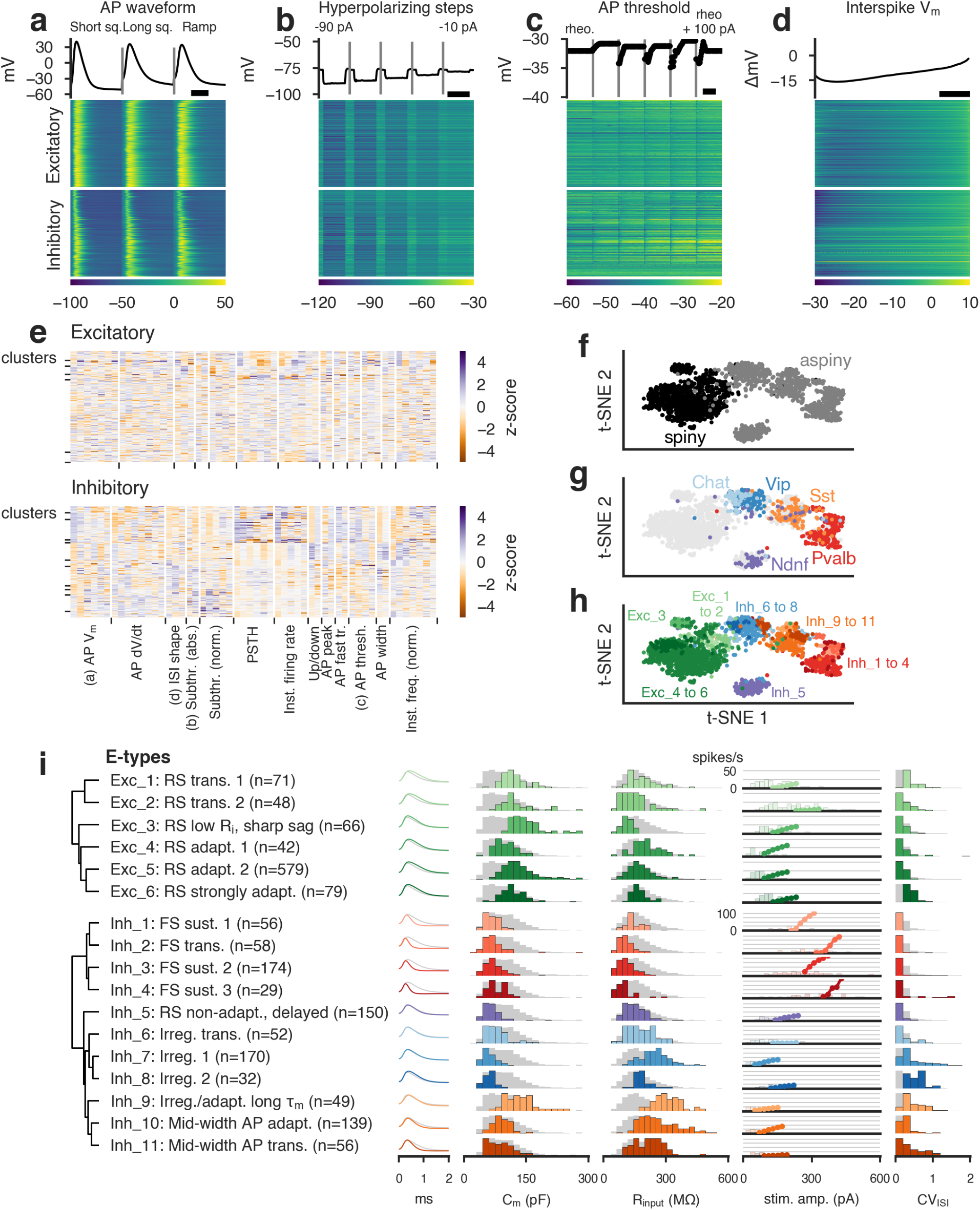
Classification of electrophysiological properties. (**a**) Action potential waveforms of n=1,851 cells evoked by a short (3 ms) current pulse, a long square (one second) current step, and a slow current ramp (25 pA/s). Example trace (top) and heat map of all responses (bottom). The cells in the heatmap are split into excitatory (spiny) cells above and inhibitory (aspiny/sparsely spiny) cells below (as determined from the images of each cell), and ordered within each of those groups by their average upstroke/downstroke ratio during long square current steps. The order of cells is the same in the heat maps of (**a**)–(**d**). In (**a**)–(**c**), vertical lines shown within examples separate data collected from different sweeps. Scale bar: 1 ms. (**b**) Membrane potential responses to hyperpolarizing current steps. Scale bar: 1 s. (**c**) Action potential threshold voltages of spikes evoked by a series of depolarizing current steps. Scale bar: 500 ms. (**d**) Interspike interval membrane potential trajectories. For a given sweep, each interspike interval duration was normalized, resampled to have a consistent number of points, aligned on the action potential threshold (set to 0 mV), and averaged together. Scale bar: 20% of interval. (**e**) Sparse principal component values collected from each data type, indicated by labels at the bottom. Each component’s values were transformed into a z-score. Rows are sorted into clusters indicated by left tick marks. (**f**) t-SNE plot with aspiny/sparsely spiny (collectively referred to as “aspiny”) and spiny neurons identified. (**g**) t-SNE plot with selected inhibitory-dominant transgenic lines identified. Only aspiny neurons from those lines are shown. (**h**) t-SNE plot with electrophysiology clusters identified. (**i**) Electrophysiology clusters (e-types) and specific features. Dendrogram on left created by hierarchical clustering based on distances between each cluster’s centroid. For AP shape, cluster averages are shown as colors and grand average across all cells shown in gray. For histograms, cluster values are shown in colors and full population is shown in gray. All histograms are scaled to their highest value. For the f-I curve, the curve was aligned on the rheobase value and averaged. The average curve was plotted at the median rheobase. All rheobase values for cells in the clusters are shown in the histograms behind the average curve.

These 13 data sets provided the foundation for an unsupervised classification of e-cell types. Given the extensiveness and complexity of the data, we first reduced its dimensionality, then applied an unsupervised clustering algorithm to identify different e-types. We applied a sparse principal components analysis^18,19^ (sPCA) procedure to each collected subset of the data, which typically identified 1–8 components that exceeded an adjusted explained variance threshold per subset (Methods). These components were collected (Fig. 2e; 53 components for excitatory neurons, 54 for inhibitory neurons), and a Gaussian mixture model (GMM) was fit to the data to divide it into clusters^19^, followed by a merging step using an entropy criterion^20^ (Methods, Supplementary Figs. 4 and 5). This procedure yielded 17 total clusters, or e-types: six clusters for the 885 excitatory neurons and eleven for the 966 inhibitory neurons. These clusters were shown to be robust by co-clustering analysis (Supplementary Fig. 5). We clustered these two classes separately since we found that clustering them together resulted in fewer excitatory-dominated clusters and a similar number of inhibitory clusters (Supplementary Fig. 6); it appeared that the diversity of inhibitory neuron electrophysiology drove the clustering in the combined analysis, and the relative similarity of excitatory neurons did not allow as much separation in that context.

We visualized the entire data set by projecting sPCA features (re-calculated using the combined data from both excitatory and inhibitory cells, yielding 52 components) onto two dimensions with the t-distributed stochastic neighbor embedding (t-SNE) method^21^. This procedure performs a nonlinear embedding that attempts to preserve the local similarity structure of high-dimensional data. The projection separated the spiny excitatory neurons from the aspiny/sparsely spiny inhibitory neurons with a reasonably well-defined border (Fig. 2f). Interestingly, populations of cells defined by transgenic drivers were also relatively coherent in this projection (Fig. 2g, Supplementary Fig. 7). For example, Pvalb-labeled cells were adjacent to Sst cells, and Vip cells were found on the other side of Sst cells. These markers label largely separate classes of interneurons^2,12^, and we observed relatively little overlap in the t-SNE projection, as well. On the other hand, we found that Chat cells, a subset of Vip cells^12,22,23^, were located within the broader region populated by the Vip cells near its border with excitatory cells. This cohesiveness suggests that there is similarity in e-features within genetically-defined populations of cortical cells, as well as distinctiveness between non-overlapping populations. The 17 clusters identified by the GMM in the higher-dimensional sPCA space were largely coherent in the t-SNE projection as well (Fig. 2h).

We next examined basic electrophysiological characteristics of the 17 identified clusters, such as the AP shape, the estimated membrane capacitance and input resistance, the average f-I curve, and the coefficient of variation of the ISI durations (CV_ISI_, Fig. 2i). We also examined several other firing pattern characteristics used to classify neurons in previous studies^6,7^, including the delay to the first spike, bursting, pausing, and spike frequency adaptation (Supplementary Figs. 8–11). Note, though, that these specific measures were not directly used in the classification; however, they were used in giving descriptive names to the clusters as shown in Fig. 2i. We also assessed the transgenic-line composition (Supplementary Fig. 12) and laminar distribution (Supplementary Fig. 13) of each electrophysiological cluster.

The excitatory clusters were relatively similar to each other, with wide APs and similar distributions of membrane capacitance and input resistance (Fig. 2i). Across the clusters, firing rates at 100 pA above rheobase rarely exceeded 30 spikes/s. The majority of excitatory neurons, including representatives from each excitatory transgenic line examined and including neurons from L2/3 through L6b, were found in the large cluster Exc_5 (“RS adapt. 2,” Fig. 2i, Supplementary Fig. 12). However, certain excitatory subpopulations had different distributions across the other excitatory clusters. Ntsr1-Cre-labeled neurons (predominantly from L6a) had higher proportions of cells in clusters Exc_1 and Exc_2 (Supplementary Fig. 12). Cells in these two clusters were frequently transient or strongly adapting (Supplementary Fig. 11), and Exc_2 had a higher median rheobase than the other excitatory clusters (Fig. 2i). A subset of L5 cells (including some cells labeled by the Rpb4-Cre, Sim1-Cre, and Glt25d2-Cre lines, Supplementary Fig. 12) were found in cluster Exc_3 (“RS low R_i_, sharp sag”), which had relatively few cells outside of L5 (Supplementary Fig. 13). This cluster displayed the most prominent bursting behavior (Supplementary Fig. 9), though bursts with a range of strengths were observed in all excitatory clusters. Clusters Exc_4 and Exc_6 were in general similar to the large cluster Exc_5; however, Exc_6 (“RS strongly adapt.”) contained relatively more superficial cells (Supplementary Fig. 13), with ˜80% of cells from L2/3 and L4 (primarily from the transgenic lines Cux2-CreERT2, Nr5a1-Cre, Scnn1a-Cre-Tg3, and Rorb-Cre, Supplementary Fig. 12). Layer 6b neurons (primarily labeled by the Ctgf-T2A-dgCre transgenic line) were nearly all in the large cluster Exc_5 (Supplementary Figs. 12 and 13). Note that these reported percentages reflect both our sampling (Supplementary Fig. 2) and the intrinsic distributions of cells; however, sampling biases alone would not produce the relative differences in laminar distributions across clusters that we observe.

The characteristics of inhibitory electrophysiological clusters were more diverse than the excitatory clusters. As expected, membrane capacitance values were nearly all lower than those of the excitatory clusters since interneurons typically have simpler dendritic arbors and lack spines, though Inh_9 was a notable exception (Fig. 2i). Clusters Inh_1 through Inh_4 contained predominantly Pvalb, Vipr2, and Nkx2.1 transgenic line-labeled interneurons (Supplementary Fig. 12). These clusters all exhibited fast-spiking (FS) firing characteristics, with narrow APs, steep f-I curves, and little adaptation (Fig. 2i). Inh_2 (“FS trans.”) often fired a transient set of high-frequency APs, especially at lower stimulus amplitudes (Supplementary Fig. 11), while the others fired in a more sustained manner up to an average of 120–160 spikes/s at 100 pA above rheobase (upper range 200–300 spikes/s). Delays in the onset of firing were more common in Inh_3 (“FS sust. 2”) and Inh_4 (“FS sust. 3”) than other clusters (Supplementary Fig. 8); still, most cells in those clusters did not exhibit a substantial delay. Pauses in firing were observed across all four FS clusters (Supplementary Fig. 10). We did not observe strong laminar biases in any of these clusters, though deeper cells in L5 and L6 may be slightly enriched in Inh_1 and Inh_2 vs Inh_3 and Inh_4 (Supplementary Fig. 13).

Most cells labeled by Ndnf and a subset of those labeled by Htr3a, Nkx2.1, and Nos1 transgenic lines were found in the Inh_5 (“RS non adapt., delayed”) cluster, which frequently exhibited late-spiking behavior (i.e., long delays at amplitudes near rheobase, Supplementary Fig. 7). These cells occasionally exhibited pauses (Supplementary Fig. 10) and were more frequently found in L1 and L2/3 (Supplementary Fig. 13).

Cells labeled by Vip-Cre, Chat-Cre, and a subset of Htr3a-Cre labeled cells were found in clusters Inh_6 through Inh_8 (Supplementary Fig. 11) with most cells in L1 through L4 (Supplementary Fig. 13). These three clusters had relatively wide APs compared to other inhibitory clusters (Fig. 2i). Inh_6 was highly transient (Supplementary Fig. 11), and Inh_7 and Inh_8 had higher CV_ISI_ values, indicating more irregular firing patterns (Fig. 2i).

Sst-Cre labeled cells were most often found in clusters Inh_9 through Inh_11. Inh_9 contained mostly cells from L5 and L6, as did Inh_10 to a lesser extent, while Inh_11 contained cells from more superficial layers (Supplementary Fig. 13). Inh_9 cells had wider spikes than Inh_10 or Inh_11 (Fig. 2i) and exhibited either an adapting pattern of firing similar to Inh_10 or more irregular firing (Supplementary Fig. 11). Inh_11 firing was more frequently transient (Supplementary Fig. 11) or had longer pauses in firing after an initial set of APs (Supplementary Fig. 9). Inh_9 (“Irreg./adapt. long τ_m_”) was also tightly associated with cells labeled by an intersection of Sst and Nos1 drivers (Supplementary Fig. 12). All three clusters had relatively high input resistances and consequently low rheobase values.

Taken together, these findings support treating the clusters identified with our unsupervised analysis as different e-types^6^. Still, it is important to note that each e-type is not entirely homogeneous. Rather, properties may vary considerably within a type as long as gradual variation is present; the existence of these intermediates means that there is no identifiable place to subdivide the cluster. The t-SNE projection, by emphasizing local similarity, can illustrate these patterns within a cluster. An example of this can be seen for cells in Inh_5 e-type (“RS non-adapt., delayed”, Supplementary Fig. 14). There, cells labeled by different transgenic lines (e.g. Htr3a, Ndnf, and Nkx2.1) and in different layers are found near cells with their same label and laminar location while exhibiting gradual transitions in electrophysiological properties across the cluster.

Comparing these results to previous electrophysiological classifications, we found that firing patterns like bursting or pausing, which have often been used to define types, frequently manifested with continuous variation across our data set (Supplementary Figs. 9 and 10). Consequently, while our results are largely in good agreement with cell-type classifications presented in previous studies in the same region^5,9,10,24,25^ and related areas and species^6,8,26^, we do note some particular differences. For example, while Markram et al.^6^ present types explicitly defined by bursting or delayed firing patterns, we find e-types that have higher proportions of neurons that exhibit these patterns (e.g., Exc_3 for bursting, Inh_0 for delayed firing), but the types are not strictly demarcated by a pattern’s presence or absence. Some observed differences could be due to differences in model systems (i.e., P14 rat somatosensory cortex in that study versus P56 mouse visual cortex here). However, these firing patterns could also mark different sub-types, especially if they are found to co-vary with other properties like molecular markers or transcriptomic signatures.

### Morphology classification

To create a comparable standardized, objective m-type classification system, we generated 3D morphological reconstructions for a subset of aspiny and spiny cells described above. Reconstructions were based on a brightfield image stack of single, biocytin-filled neurons (Fig. 1d). We chose neurons for reconstruction to provide representation across broad classes, qualitatively assessed morphological types (see Supplementary Table 2), layers and transgenic lines. The apical dendrite is most often used to distinguish among spiny, excitatory neurons (e.g., star pyramids and thick tufted neurons)^27,28^ and so we reconstructed the apical and basal dendrites and only a small portion of the initial axon of spiny neurons. In contrast, the axon of aspiny neurons has been widely recognized as the most defining feature of inhibitory cortical interneurons (e.g., L1-projecting Martinotti neurons and more layer restricted non-Martinotti neurons)^2,7,15^ and so we reconstructed the axon and dendrites of aspiny neurons to the maximal extent possible (within the confines of the slice).^23,24^

Drawing from the literature and existing analysis tool kits (e.g., L-measure^25^ and the Blue Brain Project^5^), a feature set was created that allowed extraction of multiple, well-known shape features from either dendritic or axonal branches in our reconstructions (e.g., branch number, total branch length). Furthermore, new features, such as spatial overlap between different branch types (e.g., between apical and basal dendrites) and laminar distribution pattern (e.g., branch number in layer 1) were introduced. All 122 features that were accurately represented with respect to the image data were included in the initial round of the clustering analysis (see Supplementary Table 3 for full list of calculated features.)

Due to the fundamentally different nature of the branch structures captured in the spiny and aspiny neuron reconstructions, the data set was divided between spiny and aspiny neurons (see Methods for description). Aspiny neurons were further divided into superficial and deep data sets based on relative soma depth, since they showed well-separated binary partitioning in the PCA domain, which corresponded well to laminar position (Supplementary Fig. 15). We then ran the same unsupervised hierarchical clustering analysis on three populations: aspiny-superficial (-S) (n=109), aspiny-deep (-D) (n=64) and spiny (n=199) morphologies. The number of clusters for each population was determined by maximizing between-cluster variation and minimizing within-cluster variation. In order to have unbiased clusters considering reconstruction sample sizes, an unsupervised feature selection was done by traversing non-terminal nodes of the clustering tree built with the initial feature set and retaining features significantly different between left and right branches. Twenty-five, 30 and 10 features were selected for Spiny, Aspiny-S and Aspiny-D populations, respectively (Fig. 3a and Fig. 4a,b). With retained features, the hierarchical clustering tree was rebuilt (see Methods for more detail). This method allowed for unbiased selection of different feature sets for each population.

**Figure 3:**
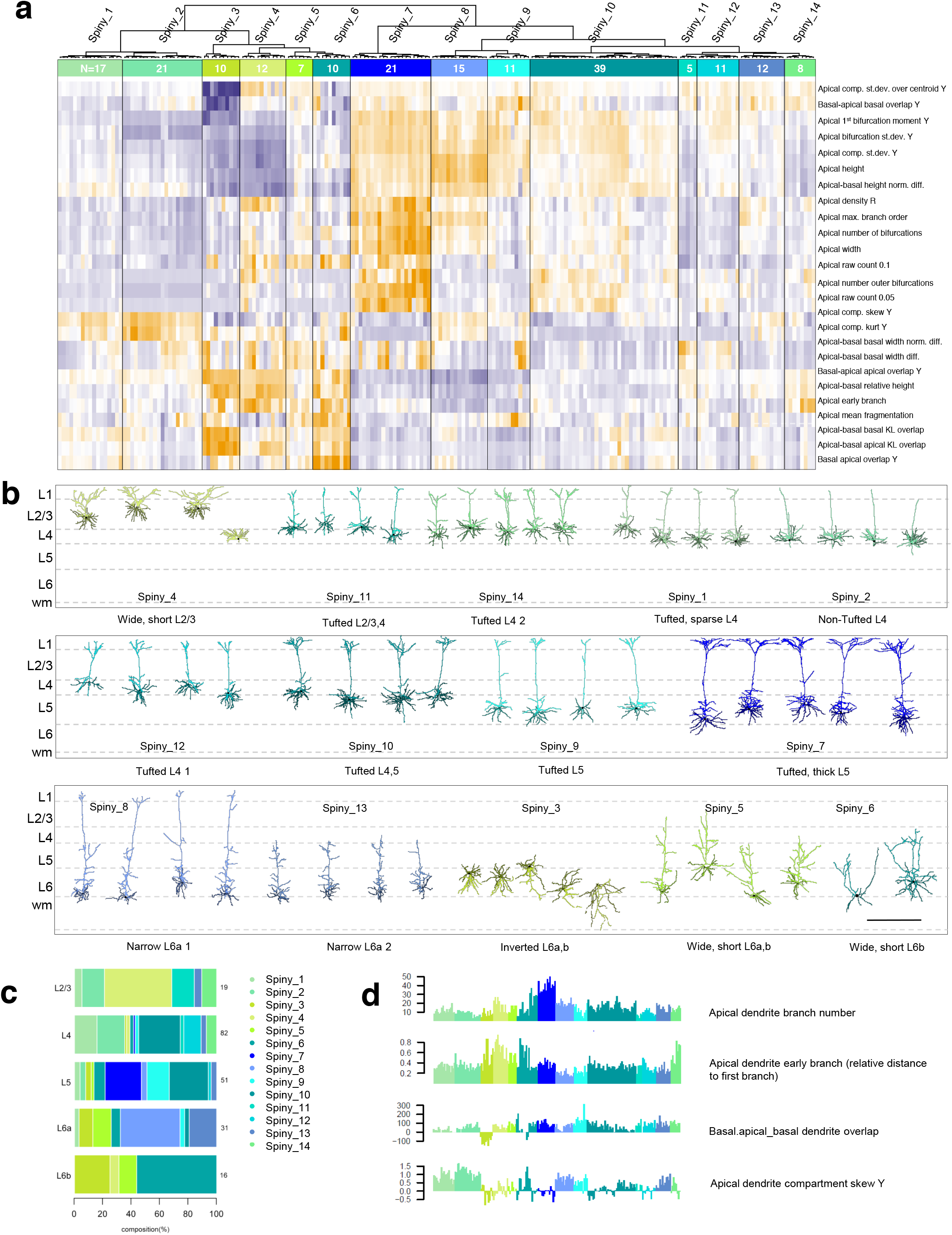
Unsupervised classification of spiny neurons into morphological types. (**a**) Heat map illustrating the distribution of 25 morphological features extracted from reconstructions of the apical and basal dendrites of spiny neurons (n = 199). Using hierarchical clustering and HybridTreeCutting, we identified 14 morphological types of excitatory neurons. The dendrogram shows the relationship among spiny morphological types. Each m-type is assigned a color that is maintained throughout the figure. Each m-type has two names, a numbered name (e.g., Spiny_1) and a descriptive name (e.g., Tufted, sparse L4). (**b**) Representative examples of each m-type are shown (with the exception of the one example of a spiny stellate neuron that clusters with cells in the Spiny_4 m-type), ordered by their representation in layers 2/3–6b rather than by cluster number. Neurons in each m-type are shown in their approximate laminar location. Apical dendrites are presented in the lighter color, basal dendrites in the darker color. Scale bar: 450 µm. (**c**) Bar graph illustrating the relative expression of each m-type across the layers. (**d**) Features of the apical and basal dendrites vary systematically across m-types. See Supplementary Table 2 for a description of how these morphological types relate to previously described types. See Supplementary Fig. 17 for all the morphologies that went into this clustering analysis, Supplementary Fig. 18 for a quantitative view of morphological features across clusters and Supplementary Fig. 16 for m-type representation across transgenic lines. All reconstructions and the corresponding images are available in our cell types database (http://celltypes.brain-map.org/).

In total, 35 m-types were identified, 14 for spiny neurons (Fig. 3), 16 for aspiny-S neurons (Fig. 4a), and 5 for aspiny-D neurons (Fig. 4b). Clusters showed good predictability by cross-validation with two supervised classifiers (support vector machine and random forest, see Methods). Co-clustering analysis^11^ also confirmed the robustness of these clusters (Supplementary Fig. 15). The relationship between m-types and Cre driver lines is shown in Supplementary Fig. 16.

**Figure 4:**
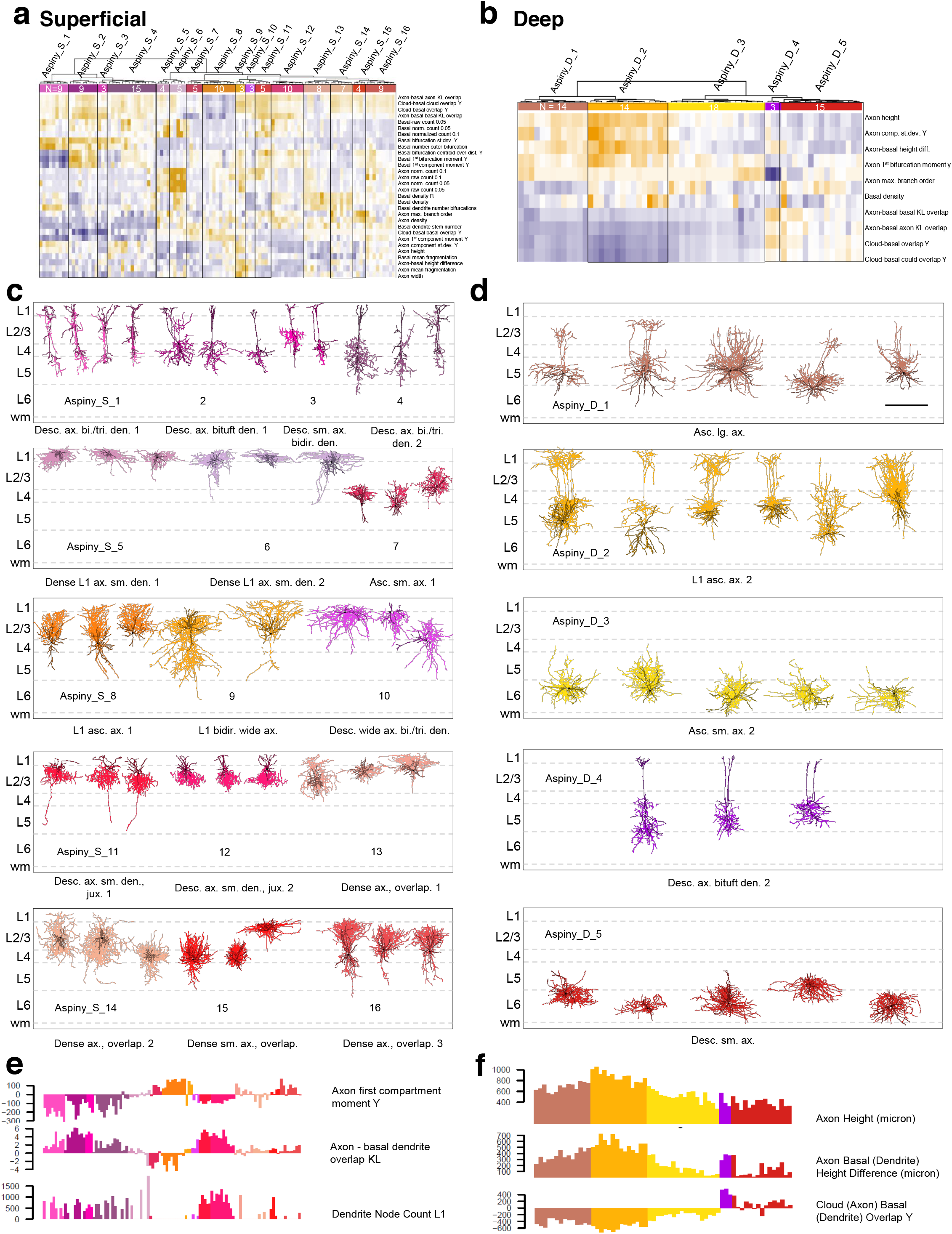
Unsupervised classification of aspiny neurons into morphological types. (**a**) Heatmap illustrating the distribution of 30 features of the axon and/or dendrites extracted from reconstructions of superficial, aspiny neurons (n = 109). Dendrogram shows the relationship among superficial, aspiny m-types. There are 16 different m-types and each m-type is assigned a color that is maintained throughout the figure. (**b**)Heatmap illustrating the distribution of 10 features of the axon and/or dendrites extracted from reconstructions of deep, aspiny neurons (n = 64). Dendrogram shows the relationship among deep, aspiny m-types. 5 m-types were identified. Across the superficial and deep aspiny populations, we identified a total of 21 m-types. Each m-type has two names, a numbered name (e.g., Aspiny_S_1) and a descriptive name (e.g., Desc. Ax. Bi./tri. Den. 1). (**c**) Representative morphologies from each quantitatively defined superficial aspiny type. Neurons are shown in an m-type-specific color in their approximate laminar location. Axons are shown in the lighter color and dendrites in the darker color. (**d**) Representative morphologies from each quantitatively defined deep aspiny m-type. Scale bar: 450 µm (shared with (**c**)). (**e-f**) Features of the basal dendrites and axon vary systematically across superficial (**e**) and deep (**f**) m-types. See Supplementary Table 2 for a description of m-type names and relationship to previously described types. See Supplementary Fig. 19 for all morphologies that went into the analysis, Supplementary Fig. 20 and 21 for a quantitative view of morphological features across deep and spiny m-types, respectively, Supplementary Fig. 22 for the layer distribution of aspiny m-types and Supplementary Fig. 16 for m-type representation across transgenic lines. All reconstructions and the corresponding images are available in our cell types database (http://celltypes.brain-map.org/).

### Spiny (excitatory) neuron m-types

The 14 spiny, excitatory m-types (Fig. 3 and Supplementary Figs. 17 and 18) were distributed across layers 2/3 to layer 6b (All slices were stained for DAPI and soma layer position was determined based on visual inspection of the cell relative to DAPI-defined layers. See methods section for more detail). Beginning in layer 2/3, there was one major morphological (m-) type, Spiny_4 (this is the only L2/3 m-type that had more than 3 neurons). These neurons had a short, densely branched apical dendrite (previously described as Type I and II neurons in rat S1^29^). Neurons with a pronounced apical dendrite with minimal (Spiny_11 and 1), or no tuft in layer 1 (Spiny_2)(often called star pyramids^6,27,30,31^), also had some representation in L2/3, but they were mainly found in layer 4.

We describe five main m-types for layer 4 (Spiny_1,2,10,12,14), which has received relatively little attention in mouse VISp. In addition to minimally and non-tufted, Spiny_1 and 2 neurons, we describe three other m-types in L4, Spiny_10, 12 and 14. These m-types are all distinguished by a larger degree of apical tuftedness. We observed only a single example of a classical spiny stellate cell (part of Spiny_4), which agrees with previous findings^32^. Spiny stellates lack a pronounced apical dendrite and resemble “stellate” inhibitory interneurons with profuse spines. The absence of spiny stellates in adult mouse V1 differs from V1 in the cat and primate^33,34^, and S1 in rat where they are the main L4 excitatory m-type^35^. Based on our sampling strategy, the predominant m-type in this layer appears to be neurons that have an apical dendrite that is relatively unbranched in L2/3 and ends with a tuft of dendritic branches in layer 1 (Tufted, Spiny_10). This m-type is found in both layer 4 and layer 5.

We describe three main m-types for layer 5 (Spiny_7,9,10). Though the morphology of layer 5 excitatory neurons has been thoroughly described for somatosensory cortex in mouse and other species, less work has been done on mouse visual cortex. In addition to the Spiny_10 m-type described above, we identified two additional layer 5 selective tufted m-types, Spiny_9 and 7 (Fig. 3b,d). Spiny_9 (and 10) neurons resemble what has been previously described as layer 5 subgroup 1B neurons in mouse V1^36^ and tall-simple^29^ and slender tufted^6,27^ neurons in mouse and rat S1, respectively. The Spiny_7 m-type, which had a larger number of branches and an increased apical tuft width, is certainly the thick tufted neurons described for multiple cortical regions^6,27,29,36,37^.

In layer 6a, we describe three main types (Spiny_5, 8 and 13). The Spiny_8 and 13 m-types had a narrow dendritic profile that was tall and short, respectively. The dendritic morphology of theses neurons very closely resembles that described for Ntsr1+ neurons that send projections to the thalamus^9,25^. Consistent with this is that these neurons were also frequently labeled by the Ntsr1 Cre line. Neurons in the Spiny _5 m-type was also found predominantly in layer 6, but they were characterized by a relatively short apical dendrite with a large width to height ratio (Supplementary Fig. 18), similar to the Spiny_4 m-type in layer 2/3 (Fig. 3b,d). These neurons resemble the short, wide branching cortico-cortical projecting neurons described by Vélez-Fort et al.^25^.

In layer 6b there are two main m-types (Spiny_3 and 6). Spiny_3 contained neurons with inverted apical dendrites while the Spiny_6 m-type contains neurons previously described as “subplate” neurons with shorter, irregularly oriented apicals^38^.

All m-types mentioned above, except Spiny_6 and 10, which are shared across layers 4 and 5 or 6a and 6b, respectively, were predominantly found in a single layer. However, they were not exclusive to a specific layer (Fig. 3c). For example, all m-types had 1–14 additional neurons in a second layer. This agrees with previous descriptions of excitatory m-type distribution in other brain regions^8,21^.

### Aspiny and sparsely spiny (inhibitory) m-types

Applying the same clustering method to the Aspiny_S and Aspiny_D populations, we identified 21 m-types for the inhibitory interneurons. Aspiny_S neurons displayed the largest diversity with 16 m-types distributed across just three layers (Layers 1, 2/3 and 4; Figure 4 and Supplementary Fig. 19, 20 and 22). Neurons in clusters Aspiny_S_1–4 were predominantly labeled by the Chat and VIP Cre lines (Supplementary Fig. 16), which are part of the Htr3a population of inhibitory neurons (Supplementary Figs. 1 and 2). They most closely resemble neurons previously described as bipolar, bitufted, small basket or double bouquet cells^5,39,40^ due to a small number of bidirectionally-oriented primary dendrites and a sparse, descending axon (Fig. 4c and 4e). Neurons in the Aspiny_S_10 cluster had similar properties, but with horizontally-oriented dendrites and a wide axon. This population of neurons has been described in other cortical areas^40^, but has not appeared in previous descriptions of mouse visual cortex.

Neurons in clusters Aspiny_S_5 and 6 and Aspiny_S_13 and 14 (Fig. 4c,e) are also members of the Htr3a class. Aspiny_S_5 and 6 neurons were found in layer 1 and had small, dense multipolar dendrites and a highly branched axon. Cells with these m-types look like cells that have previously been described as neurogliaform cells (NGC)^5,8,40,41^. Single bouquet cells were not observed in this study, though they have been previously described for mouse VISp^5^. Neurons in the Aspiny_S_13 and 14 clusters had a similar phenotype, but were located in layers 1, 2/3 and/or 4.

Clusters Aspiny_S_7 and Aspiny_S_15 and 16 (Fig. 4c,e) were labeled primarily by the Pvalb Cre line and are part of the parvalbumin (Pvalb) class of aspiny, inhibitory neurons. These neurons, found in layers 2/3 and 4, all had multipolar dendrites that overlapped with a dense axon cluster that frequently extended beyond the dendrites. Similar neurons that form axo-somatic synapses are often described as basket cells^6^, and less frequently, translaminar cells^9^, or shrub cells^5^.

Chandelier cells (ChCs) are another well-known member of the Pvalb class of inhibitory neurons. With their large boutons and unique cartridge-like axon structure, ChCs are some of the most reliably expert-identified inhibitory neurons^42^. The Aspiny_S_11 and 12 clusters (Fig. 4c,e), labeled primarily by the Vipr2 Cre line, can be clearly identified as ChCs. In this analysis, these cells, with minimally branched, L1-restricted dendrites and highly branched, L2/3-restricted axon distinguished them from other m-types. The Aspiny_S_11 cluster contained ChCs with higher density dendrites and a single axon branch that traveled beyond the main axon bundle down to layer 4/5. Though this m-type has been observed before^43,44^, it has not been described for mouse visual cortex.

The somatostatin (Sst) class of inhibitory neurons, labeled by the Sst Cre line, was represented by the Aspiny_S_8 and 9 m-types. Neurons in these two clusters had an ascending axon that frequently innervated layer 1 (Fig. 4c). Neurons with a similar morphology are most commonly described as Martinotti cells (MCs).

Aspiny_D morphologies were separated into five morphological types located in layers 5 and 6 (Fig. 4b; Supplementary Fig. 19, 21 and 22), with representation across the three main molecular classes (Htr3a, Pvalb and Sst). Neurons in Aspiny_D_1 and 2 were characterized by a large ascending axon (Fig. 4d,f). However, only neurons in the Aspiny_D_2 group actually reached layer 1. These neurons were primarily labeled in the Sst and Chrna2 Cre lines.

Comparable neurons have recently been described as “Fanning-out” Martinotti cells (within this same m-type we also see one example of a T-shaped MC)^45^. The Aspiny_D_1 group was labeled by a mixed population of Cre lines, including Pvalb Cre. The Pvalb neurons in this m-type have an axon that spans multiple layers and resemble fast-spiking, translaminar cells described previously in mouse visual cortex^9^. Neurons in the Aspiny_D_3 and 5 clusters had multipolar dendrites and an axon that ascended into an adjacent layer. The Aspiny_D_3 m-type was labeled roughly equally by Nos1 Cre and Pvalb Cre, and can be most readily compared to non-Martinotti cells (NMCs) and BCs, while Aspiny_D_5 also contained NGC-like cells. Finally, neurons in the Aspiny_D_4 cluster were labeled by the Htr3a Cre line, and had bitufted dendrites and a descending axon. These neurons are very similar to the bipolar/bitufted neurons described above (Aspiny_S_1–4). Combined, these analyses revealed 21 aspiny, morphological types distributed across all cortical layers and all major molecularly-defined inhibitory neuron groups.

Using an unsupervised clustering approach, we describe 35 m-types in adult mouse V1. The 14 spiny m-types span all layers, have layer selectivity, and in many cases, good agreement with previously defined morphological types. Additionally, we describe a larger diversity of spiny m-types in layer 4 than has been observed before. The 21 aspiny m-types also span all layers and in many cases have good correspondence with molecular classes (as defined by Cre lines) and electrophysiological types (described in the next section). We also describe additional diversity for subsets of neurons in the Htr3a class (bipolar-like cells) and Pvalb class (Chandelier-like cells).

### Correspondence between morphology and electrophysiology

As described above, we found that the t-SNE projection of electrophysiological properties allowed the relationships among the 17 e-types to be clearly visualized (Figs. 2h and 5a). We quantified the degree of separation in the projected space by calculating the Jensen-Shannon divergence (JSD) among the different e-types (Methods). The JSD is calculated as a symmetric variant of the Kullback-Leibler divergence, ranging from 0 (entirely overlapping) to 1 (entirely non-overlapping). JSD values were generally high between e-types (Fig. 5b, mean JSD = 0.95), though lower values were found between some related types (JSD range 0.51 to 1.00).

**Figure 5:**
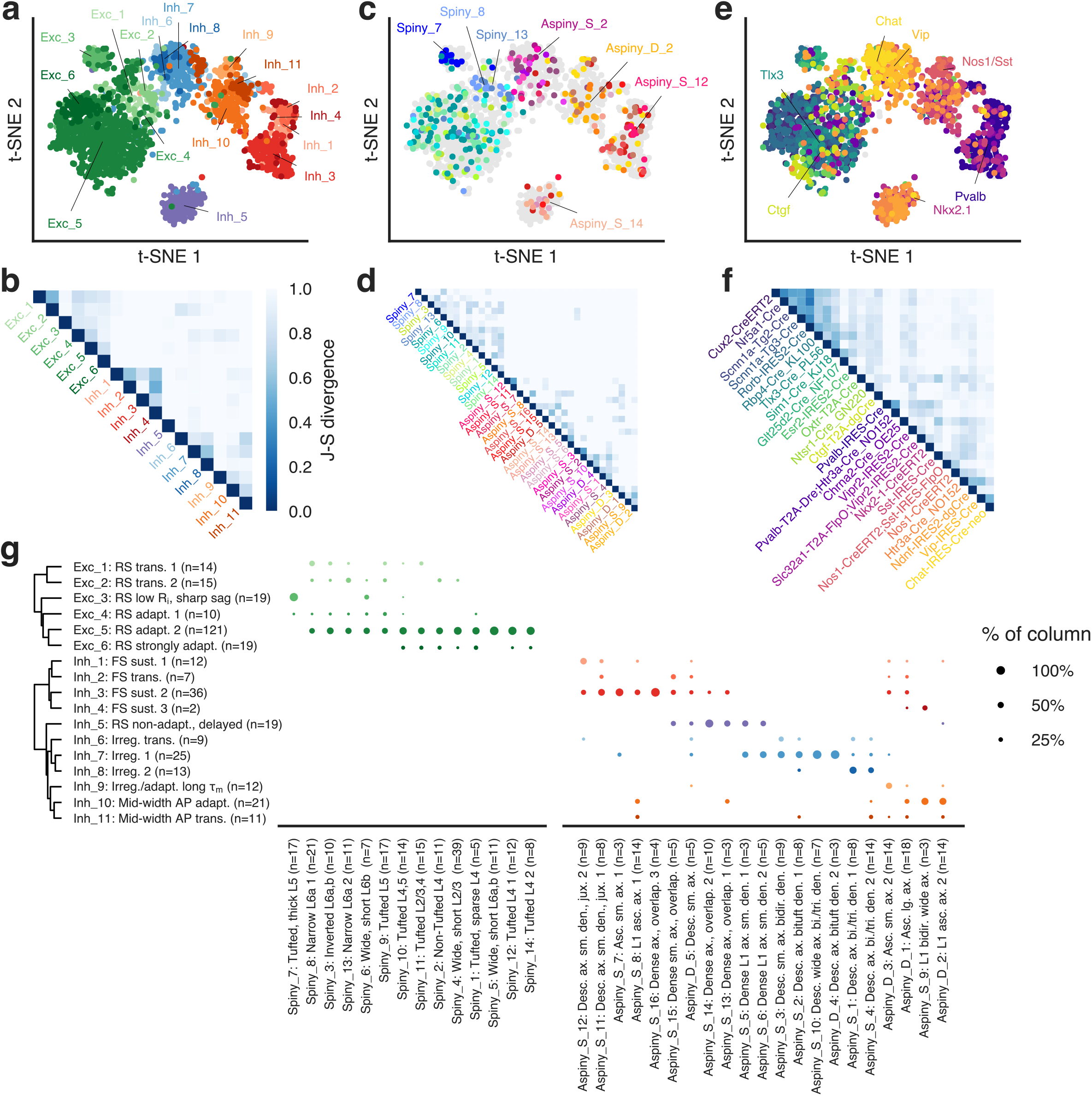
Correspondence between electrophysiology, morphology, and transgenic labels. (**a**) t-SNE plot of electrophysiological features with e-types identified. (**b**) Jensen-Shannon divergence (JSD) between e-types in the electrophysiological projection. (**c**) t-SNE plot of electrophysiological features with m-types identified. (**d**) JSD between m-types in the electrophysiological projection. The colors of the m-types are the same in c and d. (**e**) t-SNE plot of electrophysiological features with transgenic lines identified. (**f**) JSD between transgenic lines in the electrophysiological projection. The colors of the transgenic lines are the same in (**e**) and (**f**). (**g**) Correspondence between m- and e-types. The size of the marker indicates the fraction of cells with a particular e-type within a given m-type (in columns). 44 excitatory and 68 inhibitory combinations were observed out of 84 and 231 possible combinations, respectively. Reported n’s for both m- and e-types only include cells that have a morphological reconstruction.

We next used this projection defined by electrophysiological features of 1,851 cells to visualize the locations of the 372 morphological reconstructions in this space, with colors indicating membership in the 35 distinct m-types. Most m-types appeared in consistent locations in the electrophysiology-based t-SNE projection (Fig. 5c and Supplementary Fig. 23), suggesting that cells with similar morphologies frequently have similar electrophysiological characteristics. Aspiny m-types exhibited relatively little overlap with each other (mean JSD = 0.94), while more overlap was observed among spiny m-types (mean JSD = 0.90, Fig. 5c, d, aspiny vs. spiny *p* = 5.52 × 10^-6^ by two-sided Mann-Whitney U-test). Despite the greater overlap, certain spiny m-types, such as narrow L6a cells (Spiny_8, mean JSD vs. other spiny m-types = 0.95) and thick-tufted L5 cells (Spiny_7, mean JSD vs. other spiny m-types = 0.97), were found in distinct, compact parts of the t-SNE space (Fig. 5c). Both e-types and m-types exhibited a similar degree of separation in the t-SNE projection (mean JSD = 0.95 and 0.96, respectively).

When transgenic lines were overlaid on the t-SNE projection, a similar consistency in location was seen (Fig. 5e). However, as expected, many lines overlapped with each other (Fig. 5f, mean JSD = 0.89) since transgenic lines frequently label heterogeneous sets of cells (for example, some lines label populations that include both excitatory and inhibitory neurons). In addition, certain lines label subsets of populations labeled by other lines.

We next compared the m- and e-types for cells that had a 3D reconstruction (Fig. 5g). We observed 44 excitatory combinations out of a possible 6 × 14 = 84 (with 28 of them having n > 1 observations) and 68 inhibitory combinations out of a possible 11 × 21 = 231 (41 with n > 1 observations). The me-combinations with n > 1 observations exhibited a high degree of divergence in the t-SNE projection (mean JSD = 0.99). Still, we note that additional validation of the combinations with low numbers of cells will be necessary.

For excitatory cells, nearly all m-types had some degree of membership in the large e-type Exc_5 (“RS adapt. 2”). Some m-types, like the narrow tall/short cells in L6a (Spiny_8 and Spiny_13), had a higher proportion of cells with other e-types (Exc_1 and Exc_2). The thick-tufted cells of L5 (Spiny_7) had a nearly one-to-one relationship with e-type Exc_3 (“RS low R_i_, sharp sag”), most distinctive among all the excitatory m- or e-types.

Among the inhibitory cells, the majority of neurons in the NGC-like, dense axon m-types (Aspiny_S_5 and 6, 13 and 14) were in the RS non-adapting/delayed inhibitory cluster (Inh_5), and most descending axon cells with bipolar or bitufted dendrites (Aspiny_S_1–4, 10 and Aspiny_D_4) were in one of the irregularly-firing clusters (Inh_6 through Inh_8) with some bias for different e-types in different bipolar/bitufted m-types (e.g. Inh_8 and Aspiny_S_1). However, we did observe a number of cells that were labeled by the Ndnf-Cre line and solidly clustered with the NGC-like m-types, but mapped to an irregularly-spiking cluster (Inh_7). Neurons with a similar me-profile have been described in juvenile mouse neocortex.^46^

A similar situation was observed for the m-types with non-Martinotti (Aspiny_D_1 and 3) and Martinotti-like (Aspiny_S_8 and 9 and Aspiny_D_2) features. Approximately half of the Aspiny_D_1 and D_2 m-types had the Inh_9 (“Irreg./adapt. long τ_m_”) e-type, and Inh_10 (“Mid-width AP adapt.”) e-type, respectively. While the Martinotti-like, L1 ascending axon m-types were predominantly found in e-type Inh_10, the remaining cells in these m-types were fast-spiking e-types (Inh_1 to Inh_4). For all of these m-types, the fast-spiking cells were morphologically very similar to other cells with Inh_9 and 10 e-types. Surprisingly, the Aspiny_S_8 m-type, which very uniformly looked like Martinotti cells, had multiple cells labeled by Sst-Cre with the fast-spiking Inh_4 e-type. Apart from those cases, the fast-spiking e-types Inh_1 through Inh_4 were primarily associated with the basket cell-like and chandelier cell-like m-types.

### Comparison to transcriptomic characterization via specific transgenic lines

Connecting transcriptional profiles with morphological and electrophysiological properties is a powerful way to understand the functional implications of diverse gene expression^47^. Recent studies have defined mouse neocortical cell types by single-cell RNA-seq transcriptomic profiling of isolated neurons and glia^12,13^. However, it remains an open question as to how a cell’s transcriptomic identity corresponds to its electrophysiological and morphological phenotypes. To relate our findings to this effort, we took advantage of the overlap in transgenic lines used to label the cells in our study and another transcriptomic-focused effort^14^. While transgenic lines are imperfect labels of transcriptomic cell types, we were nevertheless able to identify transgenic line and layer combinations that were each selective for a small number of transcriptomic types (t-types, Fig. 6). For example, the cells in L4 labeled by the Nr5a1-Cre line are predominantly of a single t-type (Fig 6a, 95% in one type), while cells in L6 labeled by the Ntsr1-Cre line are mostly one of three related t-types (Fig 6b, 91% combined across three types).

**Figure 6:**
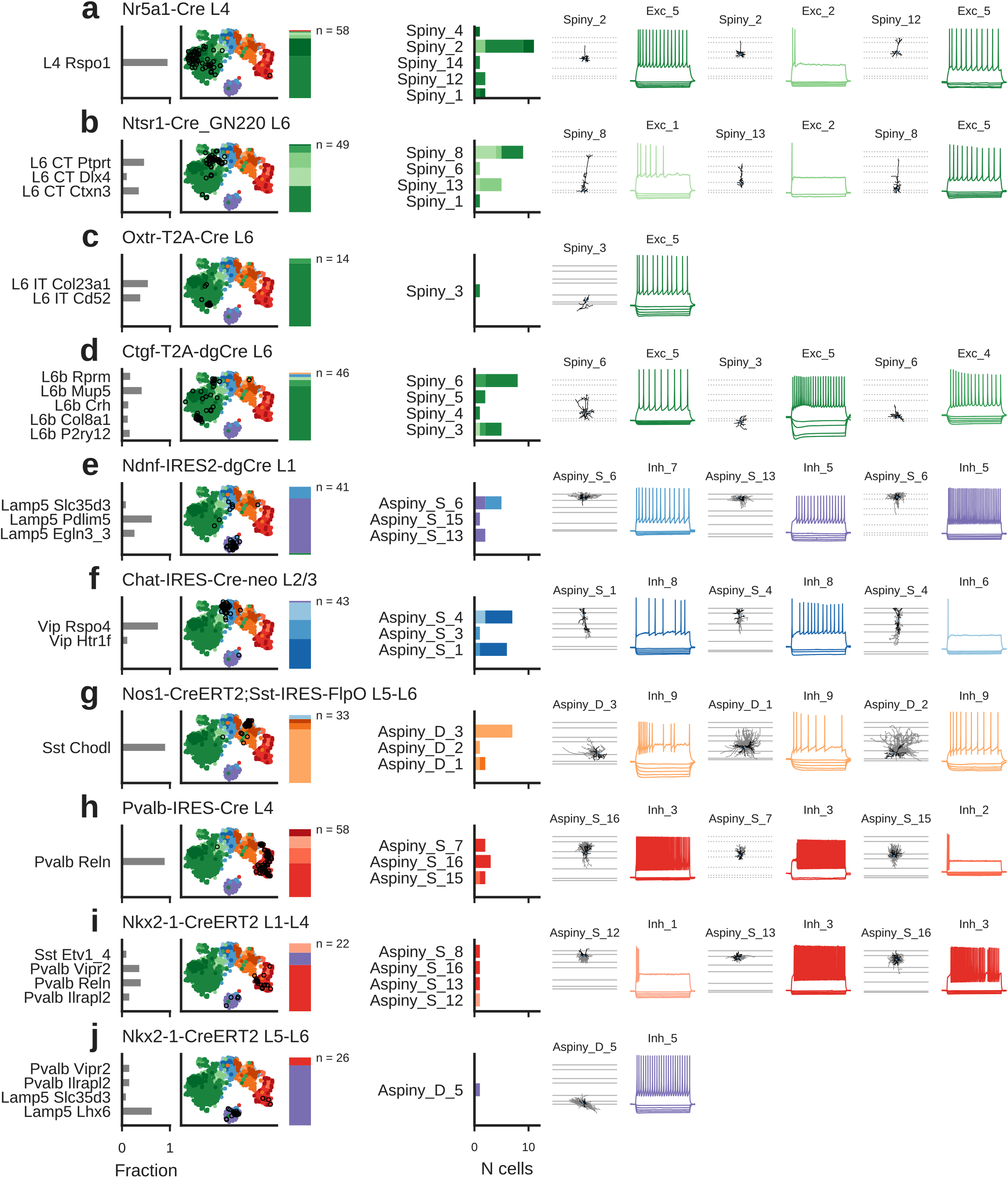
Correspondence of electrophysiological and morphological types with transcriptomic types. (**a-j**) Transgenic line/layer combinations that label relatively specific transcriptomic types (t-types) and their corresponding e- and m-types. Leftmost bar chart shows the fraction of cells in a t-type identified by another study from FACS data collected with the same transgenic line and layer sampling^14^. T-types with fractions of 0.05 or more are shown. tSNE plots show the e-types (colors) and cells in this study collected from the given transgenic line/layer combination (circles). Stacked bars indicate the proportions of e-types observed. Center bar plot shows the different m-types observed in a given transgenic line/layer combination and the distribution of me-types within that m-type. Right panels show example cell morphologies and electrophysiological responses for up to three most populous me-types for that transgenic line/layer combination. Horizontal lines on morphology plots indicate layer boundaries (solid: measured for given cell, dotted: averages when cell-specific boundaries not available). Gray is axon, black is dendrite.

Using these selective transgenic line/layer combinations, we examined the e- and m-type diversity associated with a small set of t-types. We first examined excitatory neuron types identified with this approach. Both tufted (e.g., Spiny_12) and non-tufted (Spiny_2) neurons were observed among L4 Nr5a1-labeled neurons, though these two m-types are not distinguishable by electrophysiology, suggesting some morphological heterogeneity within the single t-type VISp L4 Rspo1 (Fig. 6a). The L6 Ntsr1-labeled neurons (linked to three related VISp L6 CT t-types) had narrow tall and short morphologies (Spiny_8 and Spiny_13), as expected for deep corticothalamic (CT)-projecting neurons (Fig. 6b). In addition, most of the cells were found in a consistent location in the t-SNE projection; even cells classified into different e-types (Exc_1, Exc_2, and Exc_5) were still near each other in the projected space. L6 Oxtr-labeled neurons had very consistent electrophysiological phenotypes, as evidenced by their tight clustering in the t-SNE projection (Fig. 6c). L6 Ctgf-labeled neurons (associated with several L6b subplate t-types) exhibited some heterogeneity both in electrophysiology and morphology (Fig. 6d), with a set of cells nearer the Ntsr1 neurons in the t-SNE projection and another nearer the Oxtr neurons.

Among inhibitory neurons, the majority of neurons labeled by Ndnf in L1, associated with a handful of Lamp5 t-types, had the Inh_5 e-type and were NGCs (Aspiny_S_6 and Aspiny_S_13), consistent with previous studies^1^2,14 (Fig. 6e). The L2/3 Chat-labeled neurons, mostly associated with the Vip Rspo4 transcriptomic type, exhibited several similar m- and e-types (again, as supported by a cohesive location in the t-SNE projection, Fig. 6f). The L5-L6 neurons labeled by a Nos1/Sst intersectional strategy were quite consistent in terms of electrophysiology (Inh_9 e-type) and morphology (non-Martinotti type, Aspiny_D_3); these cells are expected to have the Sst Chodl t-type, linked to deep long-range projecting interneurons^12,14,48,49^ (Fig. 6g). L4 Pvalb-labeled cells are associated with the Pvalb Reln t-type and were found in different fast-spiking e-types and in three basket-cell-like m-types (Fig. 6h). The Nkx2–1 line labeled several relatively specific populations. In L1-L4, the line is associated mostly with the Pvalb Vipr2 type (expected to be chandelier cells^14^) and the Pvalb Reln type (Fig. 6i). Accordingly, we found that these neurons were fast-spiking (mostly in the Inh_3 e-type) and had basket-cell and chandelier-cell morphologies. In L5-L6, the Nkx2–1-labeled cells are mostly associated with the Lamp5 Lhx6 transcriptomic type, and most of these cells in our data set had the Inh_5 e-type (shared with other NGCs) and a deep NGC morphology (Aspiny_D_5, Fig. 6j).

We next used transgenic labels and laminar positions to connect our results with major subclasses of the transcriptomic taxonomy of this region of cortex^14^ (Fig. 7). Excitatory cell transcriptomic subclasses are closely associated with projection targets and laminar position; though we lack the former in our data set, we can use prior studies of associations between dendritic morphologies and long-range projections to infer relationships with excitatory me-types here (Fig. 7a). As discussed above (Fig. 6b), the Spiny_8 and 13 / Exc_1 and 2 me-types likely belong to the L6 corticothalamic (CT) transcriptomic subclass. The Spiny_7/Exc_3 me-type, containing the well-studied L5 thick-tufted cells, are expected to belong to the L5 pyramidal tract (PT) subclass^3,10,28,50^. The intratelencephalic (IT) subclass, spanning L2/3 through L6a^14^, appears from our data to be associated most strongly with the Exc_5 e-type and multiple m-types; Exc_5 is also linked to the L6b subplate transcriptomic sub-class (see Fig. 6d).

**Figure 7:**
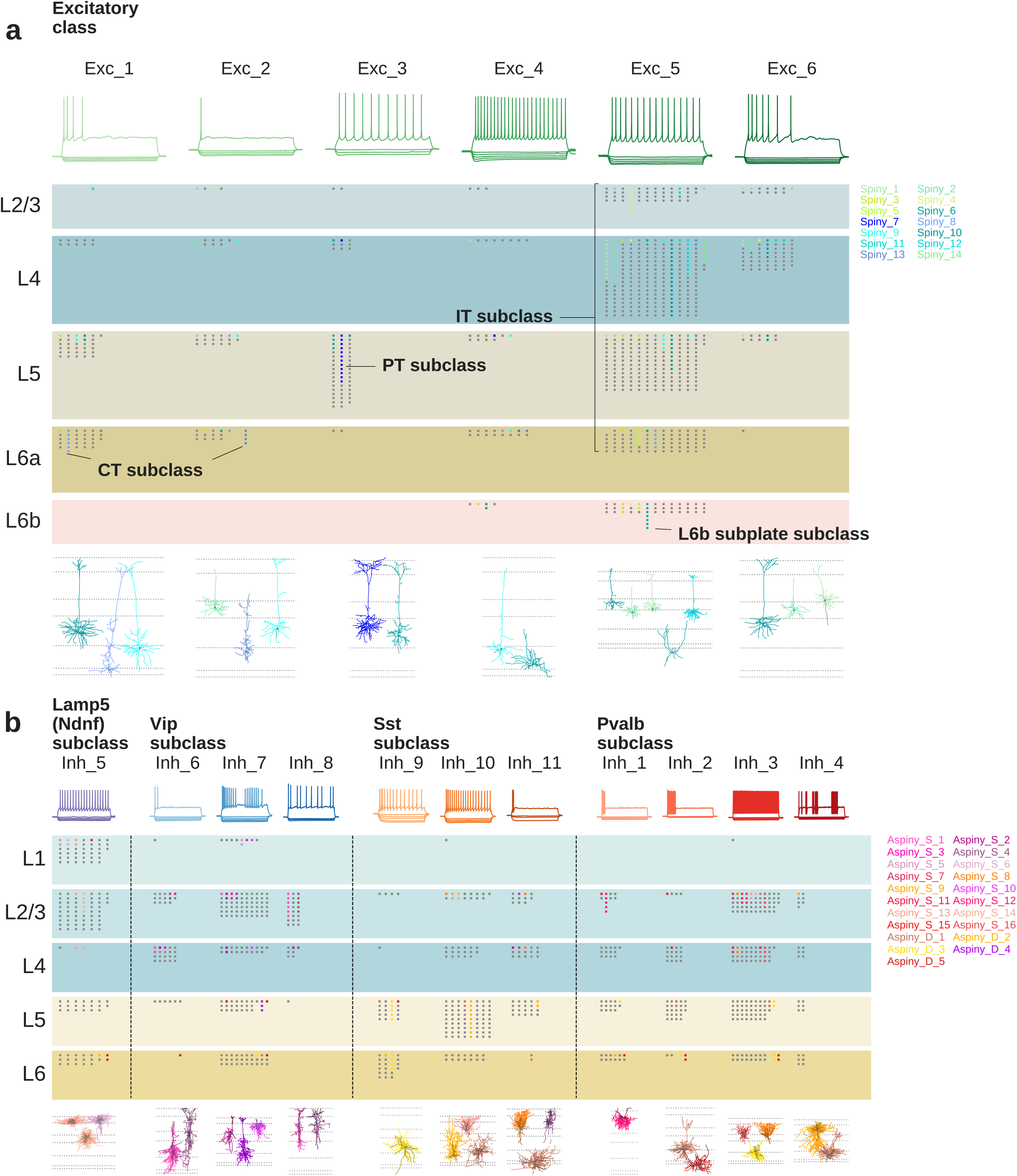
Morphological/electrophysiological types and transcriptomic subclasses. (**a**) Layer distribution of all recorded excitatory cells grouped by e-type and m-type. Top traces show representative responses for each e-type. Each marker represents a cell; colors indicate a morphologically reconstructed cell (key on right) while gray cells do not have a reconstructed morphology. Cells with the same m-type appear in the same column within an e-type. Example morphologies of the most common m-types within a given e-type are shown at bottom. Dotted lines indicate average layer borders. CT: corticothalamic, PT: pyramidal tract, IT: intratelencephalic. (**b**) Same as (**a**) for inhibitory cells. In (**b**), 81% to 100% of cells with transgenic labels within an e-type were consistent with the denoted transcriptomic subclass; we excluded the remaining cells that were inconsistent from the plot so as not to indicate correspondences between me-types and transcriptomic subclasses that are not supported by the transgenic labeling. For morphology plots, darker colors are dendrite, lighter are axon.

Inhibitory e-types were strongly associated with specific inhibitory transcriptomic subclasses (Fig. 7b). Across e-types, 81% to 100% of cells were labeled by a transgenic line consistent with a specific transcriptomic subclass; in addition, we found that related e-types were associated with the same transcriptomic subclass. Therefore, an inhibitory neuron’s electrophysiological phenotype can serve as a reasonable predictor of coarse transcriptomic identity. Our results also strongly relate specific sets of me-types to these major sub-classes.

## DISCUSSION

Obtaining the cell type composition, the “parts list”, of neural circuits is foundational to understanding circuit function. To do so it is essential to take a systematic, unbiased and quantitative approach towards cell type classification using multi-dimensional criteria, in order to resolve debates and enable the field to adopt a common set of standards. Here we describe such an effort in the morpho-electric domain in the adult mouse visual cortex. We acquired data using a standardized pipeline and uniform quality control checks such that each cell was subject to an identical process; this data production enabled the combined analysis of over 1,800 patch-clamp recordings. We developed unsupervised classification methods that were consistently applied across all recorded cells. This leads to the identification of 17 e- and 35 m-types that exhibit a large degree of correspondence with each other and show strong correlation with transcriptomically defined neuronal subclasses and types.

Conventional electrophysiological and morphological classification often relies on predetermined, cell-specific feature selection and/or qualitative assessments of firing pattern or cell shape^5,7^. Our approach, while certainly not free of biases (e.g., choice of stimuli delivered to the cells, morphological features quantified), has the advantage of being quantitative, reproducible, transferable to new data, and robust across the diverse electrophysiological responses of the entire population of adult mouse visual cortical neurons. These methods and data are also publicly accessible as part of the Allen Cell Types Database, allowing other investigators to build upon or independently evaluate this classification scheme (see http://celltypes.brain-map.org/).

When we classify neurons based on morphological features alone, we find types that have very distinct features while others display much more continuous variation across types. Though not perfect, one of the main advantages of our quantitative morphological classification approach is that it can be executed objectively to identify functionally-relevant, established and/or novel morphological types that can be applied to other systems as well. It will be important to test these methods on other datasets, such as morphologies available through the Neocortical Microcircuit Collaboration Portal (http://microcircuits.epfl.ch/#/main).

The data set presented here offers links to other studies via transgenic labels and cortical location of the recorded cells. These standardized, publicly accessible data can be used for investigations beyond what we describe here; for example, models of different levels of complexity have been built using these electrophysiological and morphological characterizations^51,52^, tools have been created to integrate these data into an automated analysis workflow^53^, and genetic – electrophysiological correlations have been inferred using these data in combination with the publicly released single cell transcriptomic data that is also part of the Allen Cell Types database^54^.

Large-scale transcriptomic studies provide informative taxonomies of cortical cell types^12–14^. Relating morphological and physiological properties to those results will refine our understanding of cell type hierarchies and the potential functions of those cell types in cortical circuits. The preliminary correspondences we find here support many of the major transcriptomic subclasses identified in the mouse visual cortex. Related inhibitory e-types exhibit good correspondence to major inhibitory subclasses, and many excitatory me-types can be related to excitatory subclasses as well. We note that we are not yet able to establish clear links to a few transcriptomically-identified subclasses (i.e., the recently described Scng inhibitory subclass and the L5 near-projecting (NP) subclass^14^); we presume that these cells are underrepresented in our data set and that adjustments to the sampling strategy could increase the rate of collection for additional study

Interestingly, we also observe some degree of heterogeneity in e- and m-types even among cells putatively from a single t-type (see, for example, Fig. 6a). However, it is known that individual t-types can exhibit substantial continuous variation within a type^14^; it remains to be seen if these variations correlate with morphological and physiological differences. The recent finding of shared t-types among inhibitory cells but different t-types among excitatory cells from separate cortical areas^14^ suggests that applying the methods of this study to a different cortical area may identify the same inhibitory e- and m-types. How excitatory e- and m-types vary across areas is more of an open question, since the divergent excitatory t-types could correspond to differences in projection targets or other characteristics. It will be of considerable interest for future studies to investigate how these three modalities co-vary on a cell-by-cell basis, to understand the relationship among molecular, physiological and morphological features as they relate to cell type definition or cell state-dependent variations.

## METHODS

Detailed descriptions of all experimental data collection methods in the form of technical white papers can also be found under ‘Documentation’ at http://celltypes.brain-map.org.

### Mouse breeding and husbandry

All procedures were carried out in accordance with Institutional Animal Care and Use Committee at the Allen Institute for Brain Science. Animals (< 5 mice per cage) were provided food and water ad libitum and were maintained on a regular 12-h light/dark cycle. Animals were maintained on the C57BL/6J background, and newly received or generated transgenic lines were backcrossed to C57BL/6J. Experimental animals were heterozygous for the recombinase transgenes and the reporter transgenes. Transgenic lines used in this study are summarized in Supplemental Table 4. Standard tamoxifen treatment for CreER lines included a single dose of tamoxifen (40 μl of 50 mg ml−1) dissolved in corn oil and administered via oral gavage at postnatal day (P)10–14. Tamoxifen treatment for Nkx2.1-CreERT2;Ai14 was performed at embryonic day (E)17 (oral gavage of the dam at 1 mg per 10 g of body weight), pups were delivered by cesarean section at E19 and then fostered. Cux2-CreERT2;Ai14 mice received tamoxifen treatment at P35 ± 5 for five consecutive days. Trimethoprim was administered to animals containing Ctgf-2A-dgCre by oral gavage at P40 ± 5 for three consecutive days (0.015 ml per g of body weight using 20 mg ml−1 trimethoprim solution). Ndnf-IRES2- dgCre animals did not receive trimethoprim induction, since the baseline dgCre activity (without trimethoprim) was sufficient to label the cells with the Ai14 reporter^12^.

### Tissue Processing

Mice (male and female) between the ages of P45-P70 were anesthetized with 5% isoflurane and intracardially perfused with 25 or 50 ml of ice cold slicing artificial cerebral spinal fluid (0.5mM calcium chloride (dehydrate), 25 mM D-glucose, 20 mM HEPES, 10 mM magnesium sulfate, 1.25 mM sodium phosphate monobasic monohydrate, 3mM myoinositol, 12 mM N-acetyl-L-cysteine, 96 mM N-methyl-d-glucamine chloride (NMDG-Cl), 2.5 mM potassium chloride, 25 mM sodium bicarbonate, 5 mM sodium L-ascorbate, 3 mM sodium pyruvate, 0.01 mM taurine, and 2 mM thiourea, pH 7.3, continuously bubbled with 95% O_2_ / 5% CO_2_). Coronal slices (350μm) were generated (Compresstome VF-300 vibrating microtome, Precisionary Instruments), with a block-face image acquired (Mako G125B PoE camera with custom integrated software) before each section to aid in registration to the common mouse reference atlas.

Slices were transferred to an oxygenated and warmed (34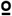C) slicing (group A) or incubation solution (group B, 2 mM calcium chloride (dehydrate), 25 mM D-glucose, 20 mM HEPES, 2 mM magnesium sulfate, 1.25 mM sodium phosphate monobasic monohydrate, 3 mM myo inositol, 12.3 mM N-acetyl-L-cysteine, 2.5 mM potassium chloride, 25 mM sodium bicarbonate, 94 mM sodium chloride, 5 mM sodium L-ascorbate, 3 mM sodium pyruvate, 0.01 mM taurine, and 2 mM thiourea, pH 7.3, continuously bubbled with 95% O_2_ / 5% CO_2_) for 10 minutes then transferred to room temperature incubation solution (group A), or allowed to cool gradually to room temperature (group B).

### Patch clamp recording

Slices were bathed in warm (34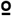C) recording ACSF (2 mM calcium chloride (dehydrate), 12.5 mM D-glucose, 1 mM magnesium sulfate, 1.25 mM sodium phosphate monobasic monohydrate, 2.5 mM potassium chloride, 26 mM sodium bicarbonate, and 126 mM sodium chloride, pH 7.3, continuously bubbled with 95% O_2_ / 5% CO_2_). The bath solution contained blockers of fast glutamatergic and GABAergic synaptic transmission, 1 mM kynurenic acid and 0.1 mM picrotoxin, respectively. Thick walled borosilicate glass (Sutter BF150–86-10) electrodes were manufactured (Sutter P1000 electrode puller) with a resistance of 3 to 7 MΩ (most 3–5 MΩ). Prior to recording, the electrodes were filled with 20 µl of Internal Solution with Biocytin (126 mM potassium gluconate, 10.0 mM HEPES, 0.3 mM ethylene glycol-bis (2-aminoethylether)-N,N,N’,N’-tetraacetic acid, 4 mM potassium chloride, 0.3 mM guanosine 5’-triphosphate sodium salt hydrate, 10 mM phosphocreatine disodium salt hydrate, 4 mM adenosine 5ʹ-triphosphate magnesium salt, and 0.5% biocytin (Sigma B4261), pH 7.3). The pipette was mounted on a Multiclamp 700B amplifier headstage (Molecular Devices) fixed to a micromanipulator (PatchStar, Scientifica).

The composition of bath and internal solution as well as preparation methods were made to a) maximize the tissue quality of slices from adult mice, and b) align with solution compositions typically used in the field (to maximize the chance of comparison to previous studies). Despite these efforts, direct comparisons with previous studies should take into account the fact that specific protocols and solution composition vary within the literature^55^. An advantage of the present study is that the same protocols / conditions were used for each cell type targeted, making it an ideal dataset to bridge data collected in different laboratories, targeting different neurons^55^.

Electrophysiology signals were recorded using an ITC-18 Data Acquisition Interface (HEKA). Commands were generated, signals processed, and amplifier metadata was acquired using a custom acquisition software program, written in Igor Pro (Wavemetrics). Data were filtered (Bessel) at 10 kHz and digitized at 50 or 200 KHz. Data were reported uncorrected for the measured^56^ –14 mV liquid junction potential between the electrode and bath solutions.

After formation of a stable seal and break-in, the resting membrane potential of the neuron was recorded (typically within the first minute and not more than 3 minutes after break-in). A bias current was injected, either manually or automatically using algorithms within the custom data acquisition package, for the remainder of the experiment to maintain that initial resting membrane potential. Bias currents remained stable for a minimum of 1 second prior to each stimulus current injection.

To be included in analysis, a > 1 GΩ seal was recorded prior to break-in and the initial access resistance < 20 MΩ and < 15% of the Rinput. To stay below this access resistance cut-off, cells with a low input resistance were successfully targeted with larger electrodes. For an individual sweep to be included: 1) the bridge balance was < 20 MΩ and < 15% of the Rinput, 2) Bias (leak) current 0 +/- 100 pA, 3) Root mean square (RMS) noise measurements in a short window (1.5 ms, to gauge high frequency noise) and longer window (500 ms, to measure patch instability) < 0.07 mV and 0.5 mV, respectively and 4) The difference in the voltage at the end of the data sweep (measured over 500 ms of rest) and the voltage measured immediately prior to the stimulus onset < 1 mV.

### Biocytin histology

A horseradish peroxidase (HRP) enzyme reaction using diaminobenzidine (DAB) as chromogen was used to visualize the filled cells after electrophysiological recording. Following electrophysiology recording, slices were fixed in 4% PFA +/- 2.5% Glutaraldehyde, then kept in PBS (4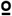C) until staining. Slices were stained with DAPI, then incubated in 1% hydrogen peroxide (H_2_O_2_) for 30 min to block endogenous peroxidases. Following permeabilization (2% or 5% Triton-X 100 detergent in PBS, 60 min, RT) slices were incubated in ABC (Vectastain, Vector Laboratories) with 0.1% Triton at 4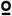C overnight to 2 days.

After a final series of three washes in 1X PBS, tissue slices were mounted on gelatin coated slides and coverslipped with glycerol-based Mowiol mounting media. Slides were dried for approximately 2 days prior to imaging. Mowiol mounting media was made in batches of 100ml and contained: 24g glycerol, 9.6g Mowiol 4–88 (Calbiochem 475904), 24ml MilliQ water, and 48ml 0.2M Tris base (pH 8.5). Slides were dried prior to imaging.

### Imaging

Mounted sections were imaged on an upright bright-field AxioImager Z2 microscope (Zeiss, Germany) equipped with an Axiocam 506 monochrome camera (6 megapixels with a 4.54 µm per pixel size). Two-dimensional (2D) images were captured with a 20X objective lens (Zeiss Plan-NEOFLUAR 20X/0.5) using the Tile & Position Zeiss Efficient navigation (ZEN) 2012 SP2 software module (Zeiss). Image quality evaluation included a qualitative evaluation of signal to noise for the imaged object (with high signal apparent in the cell body and dendrites, as opposed to background stain in the surrounding tissue, which can occur when cell filling leaks), in-focus cell body, and absent or negligible tessellation (tiling and stitching edge artifact). Overall evenness of section illumination and bounding box region for target tissue inclusion was evaluated.

Individual cells were imaged at higher resolution for the purpose of automated and manual reconstruction, quantitation and display. Light was transmitted using an oil-immersion condenser (1.4 NA). Series of 2D images of single neurons were captured with a 63X objective lens (Zeiss Plan APOCHROMAT 63X/1.4 oil), using the Tile & Position and Z-stack ZEN 2012 SP2 software modules (Zeiss). The composite 2D tiled images (X-Y resolution was set to 0.114 × 0.114 micron) were acquired at an interval of 0.28 µm along the Z-axis. Images were exported as 8-bit TIFF. Image series from individual slices or cells were processed and managed via a custom Laboratory Information Management System (LIMS).

Full dynamic range was achieved with a 20 ms exposure time and an optimal Tl VIS-LED lamp voltage control adjustment. Tiles were stitched with a minimum of 5% overlap and a 10% maximum shift. Image quality control included a z-stack plane count, a visual check for proper stitching alignment and even illumination throughout the images. 63X Z-stacks were evaluated based on quality metrics that would impact cell reconstruction, as opposed to aesthetic quality.

### Electrophysiological feature analysis

Electrophysiological features were measured from responses elicited by short (3 ms) current pulses, long (1 s) current steps, and slow (25 pA / s) current ramps. The code for feature analysis is publicly available as part of the Allen SDK. APs were detected by first identifying locations where the smoothed derivative of the membrane potential (dV/dt) exceeded 20 mV/ms. Putative AP peaks were identified as the maximum potential between detected events, and the putative AP threshold was identified by the point before the peak where the dV/dt was 5% of the maximum dV/dt. Putative APs were refined by several criteria: threshold-to-peak voltage difference must exceed 2 mV, threshold-to-peak time difference must be below 2 ms, and putative peak must be above –30 mV. The threshold was then re-calculated by finding the point for each AP where the dV/dt was 5% of the average maximal dV/dt across all APs. For each AP, several features were calculated: threshold, peak, fast trough (defined as where the dV/dt was 1% of the peak downstroke), and the width (defined as the width at half-height, where height was the difference between peak and fast trough)^57^. The ratio of the peak upstroke dV/dt to the peak downstroke dV/dt was also calculated (“upstroke/downstroke ratio”). In addition, the waveforms of the first APs elicited by the lowest-amplitude current pulses, steps, and ramps were analyzed by concatenating the 3 ms-long intervals following the AP threshold for the three conditions. The derivatives of these waveforms were also analyzed in this way.

The voltage trajectory of the ISI was also characterized to allow comparison across cells. For each cell, the sweep with the lowest stimulus amplitude that had at least five APs was identified (if a cell never fired at least five APs, the highest amplitude step was chosen instead). For each ISI, the voltage trajectory between the fast trough of the initial AP and the threshold of the following AP was extracted, and the threshold level of the initial AP was subtracted from it. The durations were normalized, then the traces were subsampled to 100 data points and averaged together. If the highest-amplitude step only elicited a single AP, a 100 ms interval following the fast trough was used in place of an ISI.

To enable comparison of AP features across the responses to long current steps given different numbers of APs across stimulus amplitudes and cells, the 1 s-long response was divided into 20 ms bins, and feature values of all APs falling within a bin were averaged. If no APs fell within a bin, the value was interpolated from neighboring bins that had APs. This was done for stimulus amplitudes starting at a given cell’s rheobase up to values +100 pA above rheobase, with a difference between amplitudes of +20 pA. If a sweep of an expected amplitude was unavailable (for example, if it failed one of the QC criteria), the missing values were interpolated from neighboring QC-passing sweeps. The instantaneous firing frequency (defined as the inverse of the ISI) was also binned and interpolated with 20 ms bins. In addition, a “PSTH” was estimated by counting APs in 50 ms bins, then converting to a firing rate by dividing by the bin duration. These two measures yield similar, but not identical, profiles of the firing pattern during a long current step response. The instantaneous firing frequency was also analyzed by normalizing to the maximum rate observed during the step to emphasize features like the adaptation of the firing frequency during the response. Though not used in the clustering analysis, the adaptation index was measured for each long step response by averaging the differences between consecutive ISIs normalized by their sums. The latency between the start of the current step and the first AP elicited was also measured.

To identify periods of high-frequency firing (“bursts”) and periods where firing temporarily but substantially slowed (“pauses”) robustly across different firing patterns, ISI shapes, and average firing rates, the following procedure was used. The coefficient of variation of the instantaneous frequency was calculated for all sets of five consecutive ISIs observed during long current steps across all cells in the data set. The distribution of these CVs was bimodal, with a large, narrow peak at low CV values (considered to represent firing at a relatively constant rate) and a wide peak at higher CV values. The minimum value of between these peaks was found at CV = 0.18. This value was used as a threshold to define segments during a response where the firing rate was relatively unchanging. For each sweep, the instantaneous firing rate was analyzed using a change-point detection algorithm^58^ to identify locations where the mean firing rate changed. The CV of the instantaneous firing rate for each segment was compared to the threshold, and if all passed, the segmentation was accepted. If not, the change-point detection penalty was lowered and the analysis was repeated until all segments passed. Once this was completed, the segment with the most APs was identified, and the firing rate ratios between all segments and that largest segment were calculated. If more than one segment was tied for the most APs, the ratios were calculated using the median of the tied segments. Segments with high ratios were considered putative bursts, and segments with low ratios were putative pauses.

Subthreshold responses to hyperpolarizing current steps were analyzed using downsampled (to averages in 10 ms bins) membrane potential traces that was concatenated together. Responses from –10 pA to –90 pA steps (at a –20 pA interval) were used, and 200 ms of the time before and after the step were included as well. In addition, the largest amplitude hyperpolarizing step response was analyzed by normalizing to the minimum membrane potential reached and the baseline membrane potential. This emphasized the “sag” in the membrane potential due to the activation of *I*_h_ observed in some cells. Though not used for clustering analysis, the input resistance was calculated by the slope of a linear fit to the minimum membrane potentials during these hyperpolarizing step responses, and the membrane time constant was estimated by exponential fits between 10% of the maximum voltage deflection and that maximum deflection. The membrane capacitance was estimated by dividing the membrane time constant by the input resistance.

### Electrophysiological classification

Data sets were built by accumulating the feature vectors in each category (e.g. AP waveform, each AP feature across long steps, subthreshold response waveforms, etc.; see Supplementary Table 1). Data from putatively excitatory cells and inhibitory cells (determined by the presence and type of dendritic spines) were analyzed separately, though similar results were observed when all cells were analyzed together (see Supplementary Fig. 6). Sparse principal component analysis^18^ was performed separately on each data set. Principal components with an adjusted explained variance exceeding 1% were kept (typically 1 to 8 components from a given data set). Analysis of inhibitory neurons yielded 54 total components, excitatory neurons yielded 53 components, and all neurons combined yielded 52 components. The components were then z-scored to standardize the scale and combined to form a reduced dimension feature matrix. The matrix was then fit with a series of Gaussian mixture models (GMMs) with a diagonal covariance matrix using different numbers of components; the GMM that minimized the Bayes information criterion was chosen as the best representation for the data^19^. Next, components of the selected GMM were iteratively merged^20^ to identify clusters that may have had non-Gaussian structure (and therefore would have been fit by the GMM with multiple components). At each step, the merge that maximized the change in entropy was identified, and the number of cells affected by the merge was recorded. The point where the rate of entropy decrease versus number of cells merged slowed was identified by a two-part linear fit, and the merges up to that point were used to define the final clusters.

Robustness of clustering was evaluated by co-clustering analysis. Random subsamples containing 80% of the data set were generated 100 times, and the subsamples were fit with a GMM using the number of components of the best GMM fit to the full data set. Components were then merged as described above. The fraction of times a pair of cells was found in the same cluster (out of the number of times both cells appeared in the same subsample) was calculated for all pairs. Average co-clustering fractions were calculated between all clusters defined by the analysis of the full data set.

The electrophysiological feature matrix used in the clustering analysis was also visualized with a two-dimensional projecting using the t-distributed stochastic neighbor embedding (t-SNE) technique^21^. Cluster identities and other features of the cells were visualized using this projection throughout this study. Comparisons between groups in the t-SNE projection were made by calculating the Jensen-Shannon divergence^59^ (JSD). The distributions of each group were calculated as two-dimensional histograms with the t-SNE space divided into a set of 20 × 20 bins. The JSD value between groups P and Q was computed as 
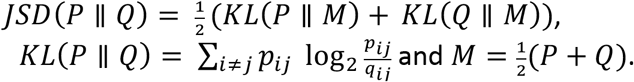

### Anatomical location

To characterize the position of cells analyzed from mouse brain, a 4-step process was used. Briefly, 20x brightfield and/or fluorescent images of DAPI (4’,6-diamidino-2-phenylindole) were analyzed to determine layer position and region, of biocytin-filled cells. Soma position was annotated and used to calculate soma depth relative to pia and white matter. Individual cells were then manually placed in the appropriate cortical region and layer within the Allen Mouse Common Coordinate Framework (CCF) by matching the 20x image of the slice with a “virtual” slice at an appropriate location and orientation within the CCF. Using the DAPI image, laminar borders were also drawn for all reconstructed inhibitory neurons.

### Dendrite type assignment

The dendritic morphology of each neuron (N=1851) was identified as either aspiny, sparsely spiny or spiny^60^ based on a qualitative assessment of the neuron’s dendrites by viewing the slides under the microscope or in the 63X image. These different dendritic types roughly equate to interneurons (aspiny and sparsely spiny) and pyramidal or spiny stellate neurons (spiny). Aspiny dendrites were defined by the absence of spiny protrusions and checked against a lack of a pronounced apical dendrite and/or axon emerging from the soma or dendrite at odd angles, and branched extensively. Sparsely spiny dendrites were defined by these same features, except that spines appeared with infrequent to moderately frequent expression (∼ 1 spine/10 microns). Spiny dendrites were defined by the presence of frequent spiny protrusions (approximately one spine per 1–2 microns), and validated by axon that descended perpendicularly down to the white matter with sparse, proximal branching occurring at right angles to the primary axonal branch and/or a pronounced primary, apical dendrite.

### Morphological reconstruction

Three-dimensional (3D) reconstructions of the dendrites and the initial part of the axon (spiny neurons) and/or the full axon (aspiny/sparsely spiny neurons) were generated for a subset of neurons with good quality electrophysiology and biocytin fill. 3D reconstructions were generated based on a 2D image stack that was run through a Vaa3D-based image processing and reconstruction pipeline^15^. The process included a variable enhancement of the signal to noise ratio in the image^61^. The enhanced image was then used to generate an automated reconstruction of the neuron using Neuron Crawler^62^ or TReMAP^63^. Automated reconstructions were then extensively manually corrected and curated using a range of tools, e.g., virtual finger, polyline, in the Mozak extension (Zoran Popovic, Center for Game Science, University of Washington) of Terafly tools^61,64^ in Vaa3D. Every attempt was made to generate a completely connected neuronal structure while remaining faithful to image data. If axonal processes could not be traced back to the main structure of the neuron, they were left unconnected. Using the most updated version of the Mozak-Terafly-Vaa3d tools, on average, dendrite only reconstructions of spiny neurons took 4.5 hours and full reconstructions of the axon and dendrites of aspiny neurons took 16 hours. Connected and disconnected axon components (axon cloud) were used in the quantitative analysis. As a final step in the manual correction and curation process, an alternate analyst checked for missed branches or inappropriate connections. Once the reconstruction was deemed complete, multiple plugins were used to prepare neurons (saved as SWC file) for qualitative and quantitative morphological analyses.

### Morphology feature design, feature selection, and clustering

Features were designed to describe characteristics of neuron morphology based on reconstruction data. They can be categorized into branching pattern, size, density, soma position, estimated layer-by-layer node counts, y-directional profile^23^, and overlap feature between apical (for spiny neurons) or axon (for aspiny neurons) and basal dendrite. The same set of features listed in Supplementary Table 3 were calculated for axon, apical dendrite, basal dendrite, cloud (connected plus all disconnected axon branches), and neurites.

From the initial set of features, ones with low variance (coefficient of variance < 0.25) were removed and a representative feature was chosen among highly correlated features (correlation > 0.95). These features were scaled to form a feature set on which a standard hierarchical clustering with Ward’s agglomeration method using Euclidean distance was applied. The number of clusters is determined by cutting the hierarchical tree using R function CutreeHybrid^26^. Further feature selection was done by traversing each non-terminal node of the tree and selecting features significantly different between left and right branches with the criteria, adjusted t-test p-value < 0.01 and |log (foldchange)| > log(1.25). With this reduced set of features, the clustering tree was rebuilt. This feature reduction and tree update continued until there was no change in the number of clusters or no further reduction in features.

Classifiers used in checking predictability of clusters were designed by R functions svm() and RandomForest(). Prediction rates for clusters from Spiny, Asiny-S, and Aspiny-D groups were 87.4(85.9)%, 84.4(78.0)% and 92.2(87.5)% by SVM(RF) classifiers. Robustness and homogeneity of clusters were shown by co-clustering analysis^11^, which accumulated over all 100×10 clustering runs with 90% subsampling in 10-fold cross validation manner and summarized them in co-clustering rates (Supplementary Fig. 15). Lower co-clustering rates in the Spiny groups were due to the existence of subgroupings within the clusters. For Aspiny groups, more informative features elucidating axon’s various patterns would help solidify clusters, especially for Aspiny-D neurons.

### Transcriptomic correspondences

The associations between transgenic lines and transcriptomic types (t-types) were investigated using data from a recent study^14^ to establish preliminary correspondences with the results here. Specific transgenic line/layer combinations that labeled a small number of t-types, defined as having five or fewer t-types containing at least 5% of the cells from that line and layer set, were identified. Correspondences between transgenic lines and broader inhibitory sub-classes defined by that study (Lamp5, Vip, Sst, and Pvalb) were also analyzed. Cells labeled by a transgenic line were considered to be consistent with a given inhibitory sub-class if at least 7% of inhibitory cells labeled by that line were found in that sub-class.

### Data availability

The electrophysiological and morphological data supporting the findings of this study are available in the Allen Cell Types Database, celltypes.brain-map.org. Morphological data are also available through the NeuroMorpho.org repository^65^, neuromorpho.org.

### Code availability

The Vaa3D morphological reconstruction software, including the Mozak extension, is freely available at www.vaa3d.org and its code is available at https://github.com/Vaa3D. The code for electrophysiological and morphological feature analysis is available as part of the open-source Allen SDK repository (alleninstitute.github.io/AllenSDK).

## ACKNOWLEDGEMENTS

We thank Zoran Popovich for creating the Mozak custom user interface for the 3D reconstruction software, Terafly-Vaa3D. We thank Barb Berg, Sil Coulter, Chinh Dang, and Allan Jones for leadership and guidance. This work was funded by the Allen Institute for Brain Science, and by National Institutes of Health grant U01MH105982 to H.Z. The authors thank the Allen Institute founder, Paul G. Allen, for his vision, encouragement and support.

## AUTHOR CONTRIBUTIONS

H.Z. and C.K. conceived the study. T.L.D., B.T., T.N.N., and E.G. contributed to the generation and/or characterization of specific transgenic mouse lines. J.H., M.G, M.R., and N.B provided mouse colony management. N.D., S.P, N.T., T.C., M.K., J.S., K.C., H.T., and E.B. prepared tissue slices. A.O., D.H., K.H., S.J., L.N., L.K., and R.M. performed electrophysiology experiments. T.L., M.M., K.B., A.D., C.H., D.P., A.G., T.E., H.G., and K.B. processed slices for biocytin staining. S.C., C.C, M.G., S.D., N.D., K.N., and L.P. imaged biocytin-stained slices and cells. S.A.S., T.D., M.F., A.H., D.S., N.T., R.D., G.W., A.M., R.A.D., and S.K. reconstructed neurons and provided anatomical annotations. N.W.G., X.L., C.L., A.B., J.B., S.A.S., K.G. performed analysis. J.B., A.O., J.T., B.L., P.C., S.A.S., and N.D. contributed to methods development studies. H.P., Z.Z., B.L., C.F., J.P., C.S., M.S., D.R., T.B., D.C., and T.J. designed, wrote, or built tools for pipeline data generation. S.M.S. provided program management support. J.W.P., C.K., H.Z., A.B., J.B., T.L., M.M., N.G., P.N., L.P., S.A.S., N.D., and S.P. organized and managed pipeline data generation. N.W.G., K.G., L.N., W.W., R.Y., D.F., and A.S. organized and managed pipeline data storage and processing. N.W.G., C.A., A.A., S.M., H.P., C.T., M.J.H., J.B., T.J., G.S.L., J.T., B.L., G.J.M., E.L., J.W.P., C.K., H.Z., A.B., S.A.S., J.H., and B.T. provided scientific direction. N.W.G., C.L., J.B., and S.A.S. prepared the figures. N.W.G., J.B., and S.A.S. wrote the manuscript in consultation with all authors. H.Z., C.K., and A.A. provided substantial review and edits to the manuscript.

---

**Supplementary Figure 1:**
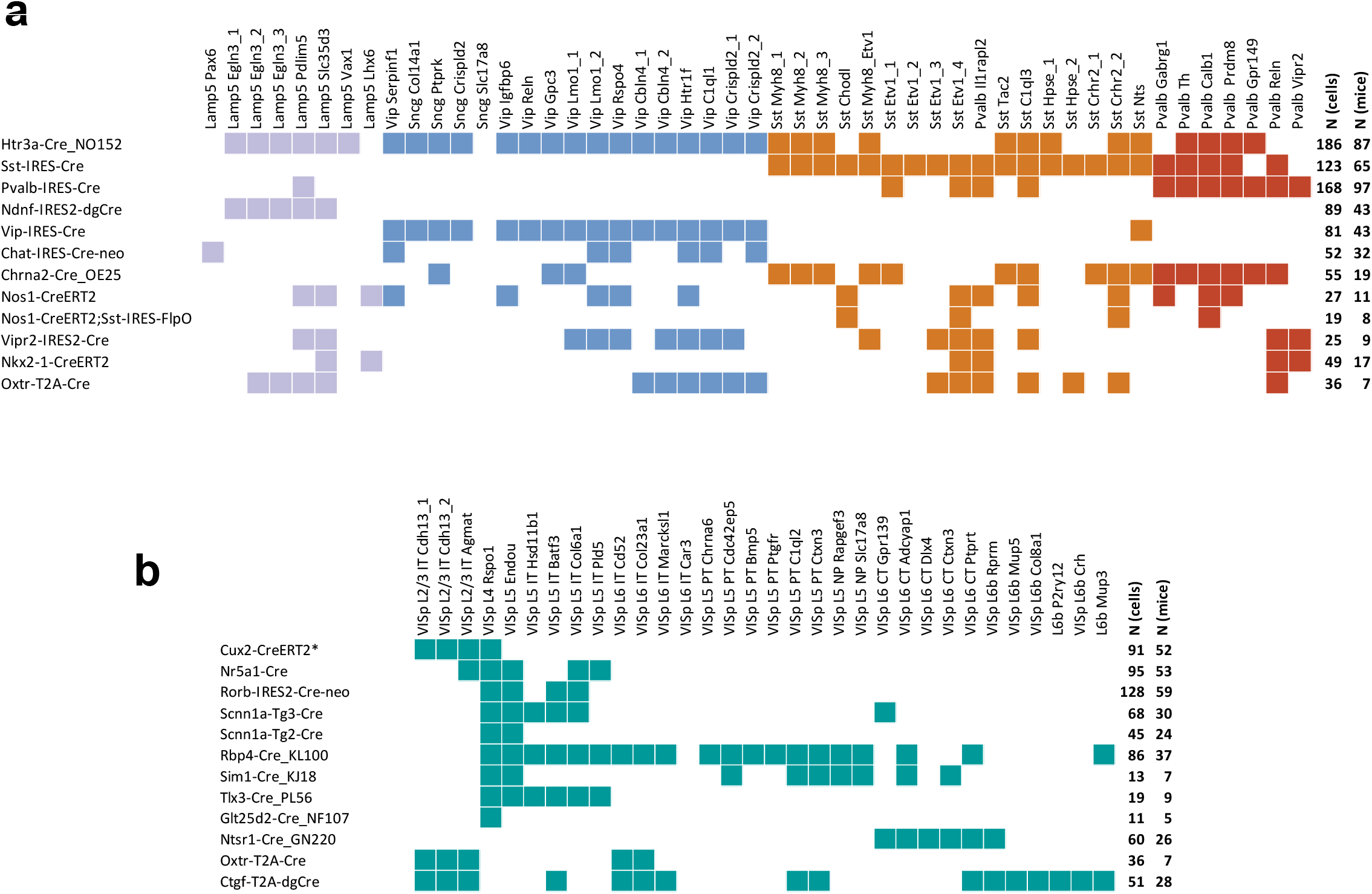
Transgenic line-based sampling strategy. (**a**) Inhibitory-dominant transgenic lines used in study (rows) and inhibitory transcriptomic types (columns) labeled by each line. Broad lines (e.g. Htr3a-Cre_NO152, Sst-IRES-Cre, Pvalb-IRES-Cre) were chosen to cover the majority of inhibitory transcriptomic types in VISp. Additional lines were chose to fill in missing types and to label specific types more selectively. (**b**) Excitatory-dominant transgenic lines used in study (rows) and excitatory transcriptomic types (columns) labeled by each line. Excitatory-dominant lines tended to be more selective and were enriched in specific cortical layers.

**Supplementary Figure 2:**
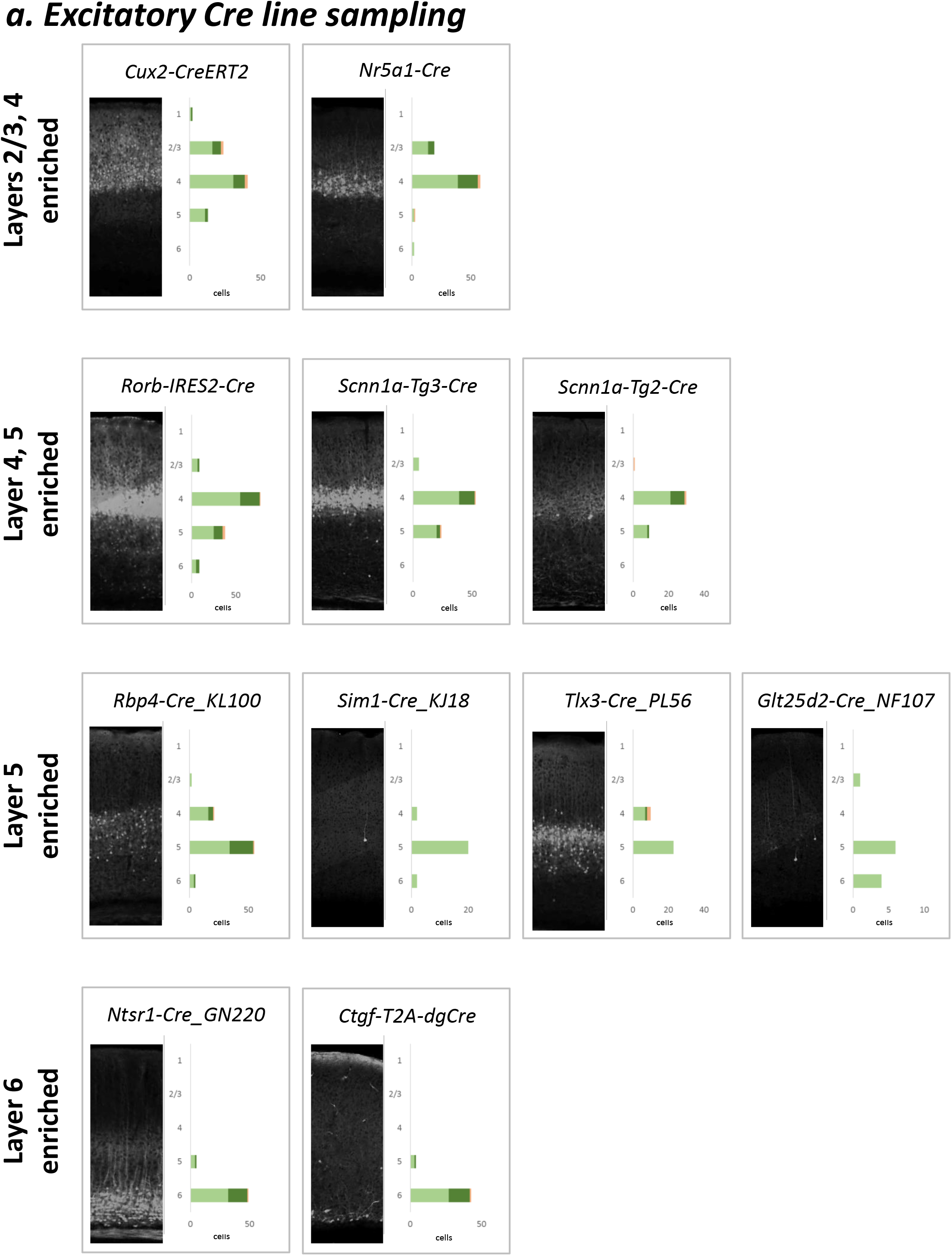
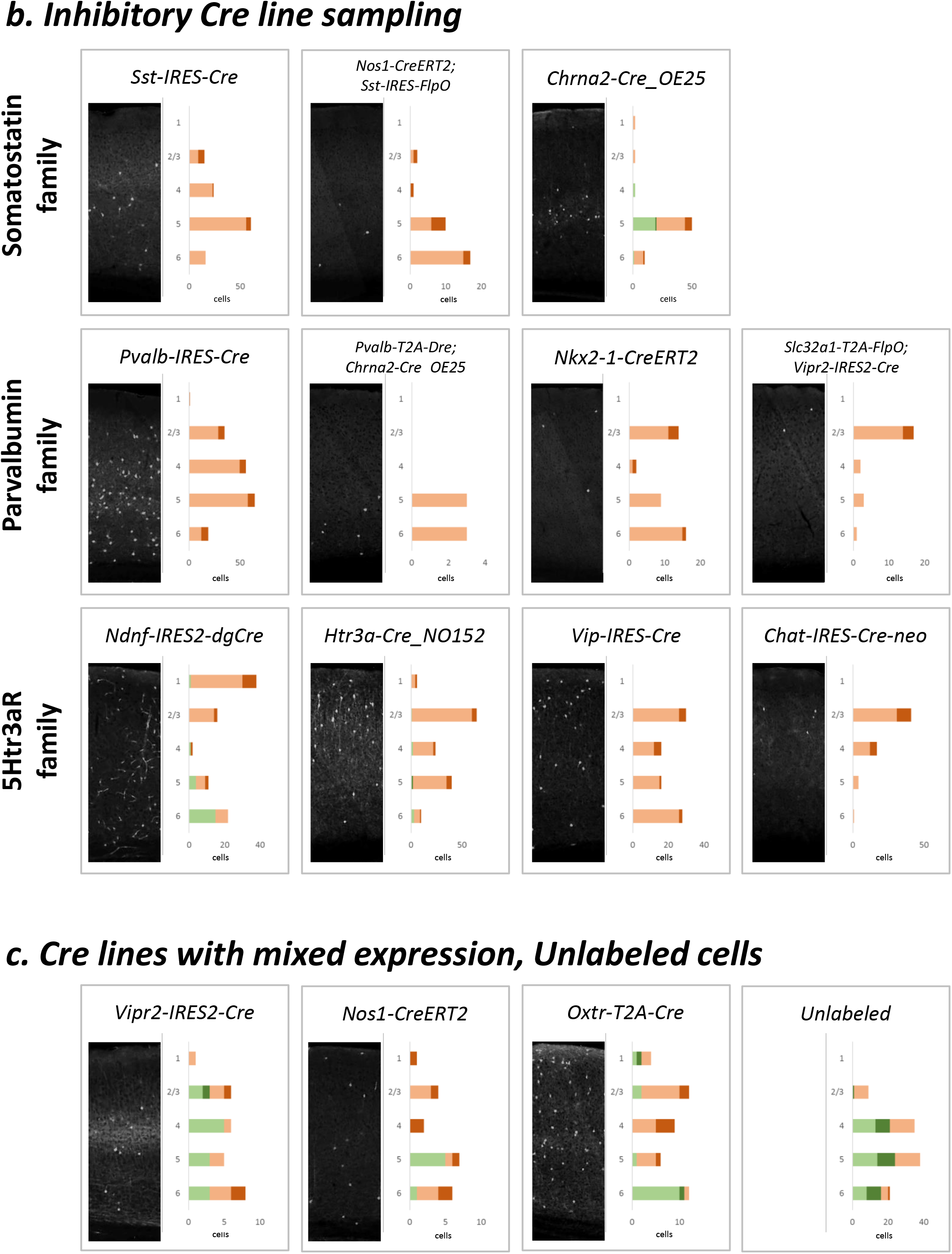
Sampling results per transgenic line. A summary of the layer distribution of cells recorded from each transgenic line. For each transgenic line: *Left*, 2-photon composite image of coronal slice of primary visual cortex showing distribution of fluorescent neurons. Images were obtained and processed as described in Oh et al., Nature 2014. *Column 2*, Stacked histogram of spiny (green) and aspiny (brown) cells sampled. Darker bars indicate those cells that were also morphologically reconstructed.

**Supplementary Figure 3:**
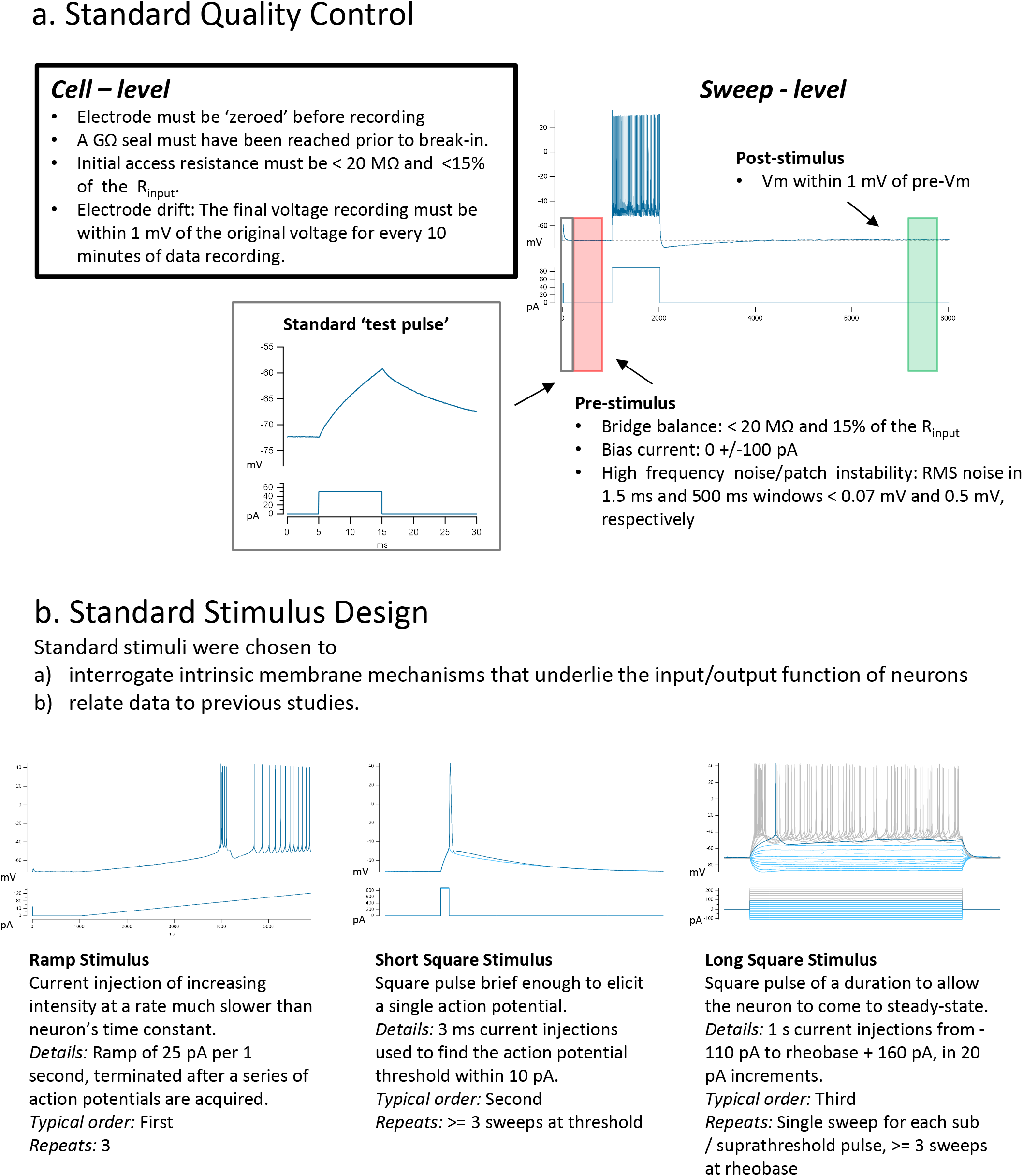
Electrophysiology quality control and stimuli. (**a**) Each cell was subject to a number of gates to insure stable quality recordings. Sweeps were manually inspected for artifacts and for correct bridge balance settings using a short standard ‘test pulse’ preceding the stimulus. In addition, poor quality sweeps were automatically rejected from analysis using a series of criteria before and after the stimulus. (**b**) The electrophysiology properties of each cell were probed using three standard stimuli: long and short square steps, as well as a ramp current injection.

**Supplementary Figure 4:**
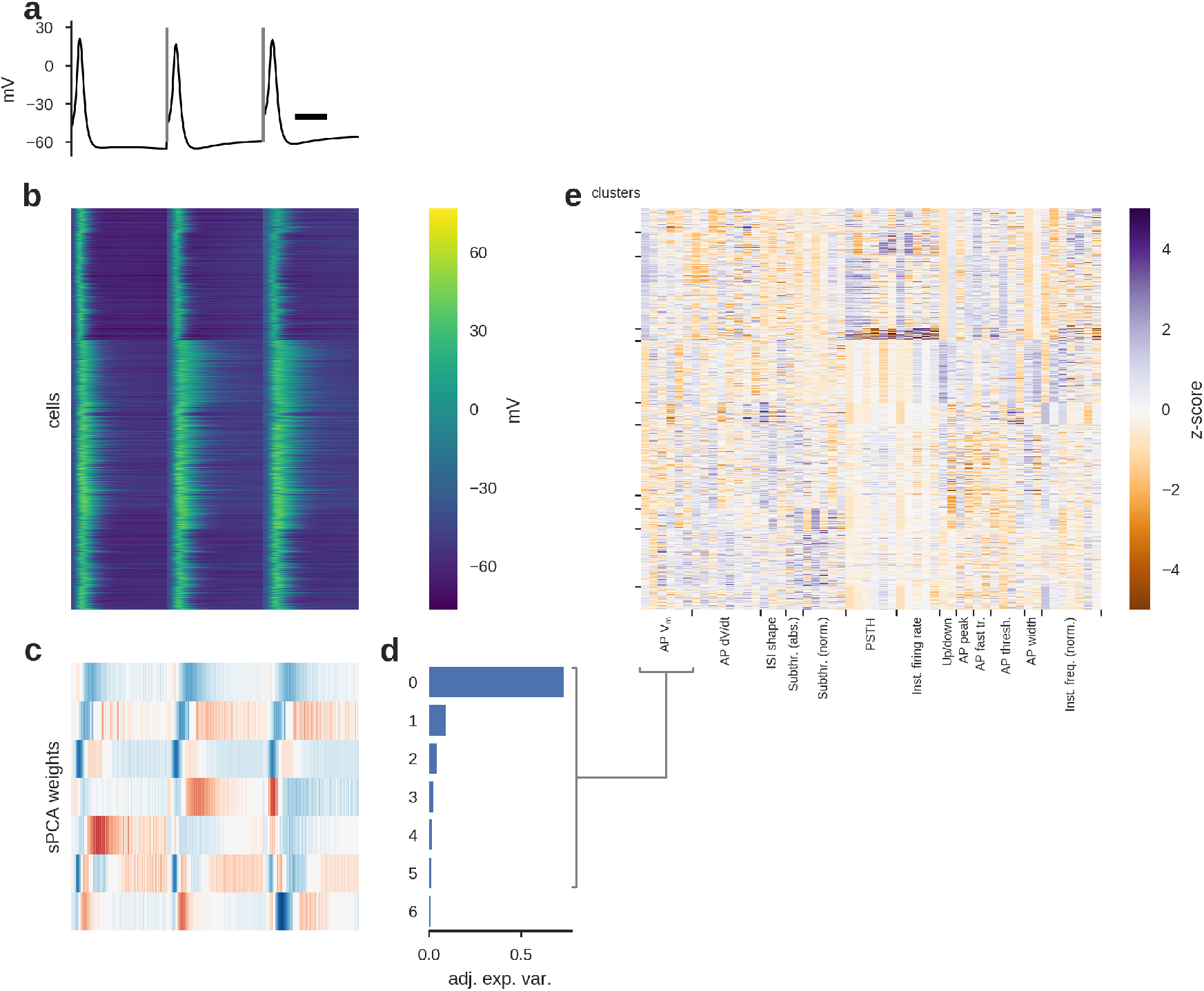
Electrophysiology clustering methodology. (**a**) Example action potential waveforms of an example cell evoked by a short (3 ms) current pulse, a long square (one second) current step, and a slow current ramp (25 pA/s). (**b**) Heat map of all action potential waveforms from inhibitory cells (n=966). (**c**) Sparse principal component weights of the data in b. Time scale is the same in (**a-c**). (**d**) Adjusted explained variances of sparse principal components shown in (**c**). (**e**) Sparse principal component values collected from each data type, indicated by labels at the bottom. For example, the seven sparse principal components obtained from the action potential waveforms populate the first seven columns of the matrix in (**d**). Component values were transformed into a z-score. Rows are sorted into clusters indicated by left tick marks (Methods).

**Supplementary Figure 5:**
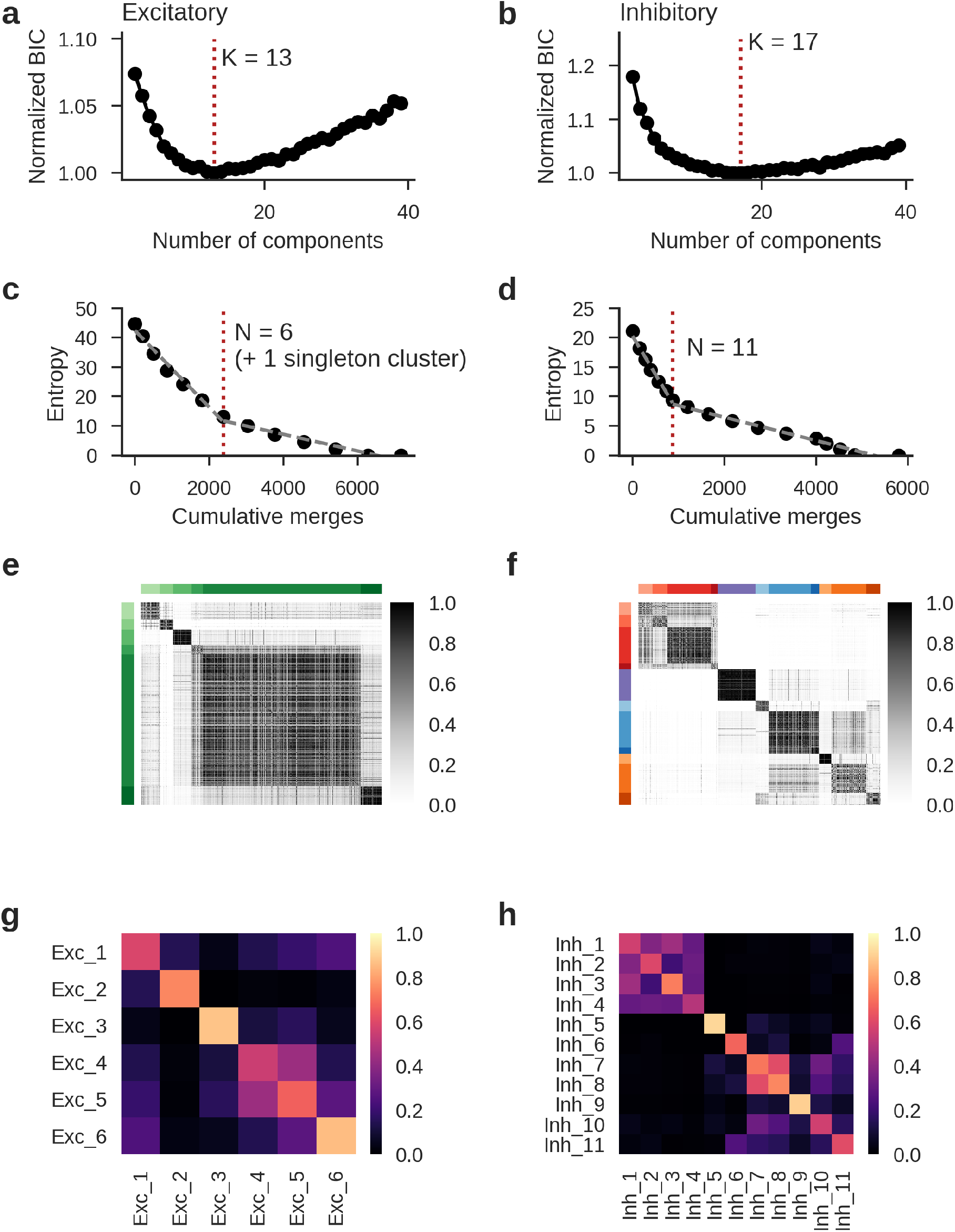
Merging GMM components and co-clustering analysis. (**a-b**) Bayesian information criteria (BIC) values (normalized to the minimum) for Gaussian mixture models fit using the excitatory (**a**) and inhibitory (**b**) neuron data with different numbers of components. The model with the lowest BIC (K=13 components in (**a**), K=17 components in (**b**)) was selected as the best representation of the data. (**c-d**) Entropy as Gaussian mixture model components were merged, plotted against the cumulative number of samples merged. Merging was stopped when the rate of entropy decrease slowed (excitatory: N=6 clusters with n > 1 cell in (**c**); inhibitory: N=11 clusters in (**d**)), determined by a two-part piecewise linear fit (Methods). (**e-f**) Pairwise co-clustering results for excitatory (**e**) and inhibitory (**f**) cells. 100 random subsamples containing 80% of the data were generated and clustered by GMM fit and merging. Heatmap shows the fraction of times a given pair of cells were in the same cluster. Cells are ordered by clusters determined from the full data set, indicated by row and column colors. (**g-h**) Average co-clustering fractions between final clusters for excitatory (**g**) and inhibitory (**h**) cells. All within-cluster fractions were observed to be at least 0.1 than the maximum cross-cluster fractions for each cluster.

**Supplementary Figure 6:**
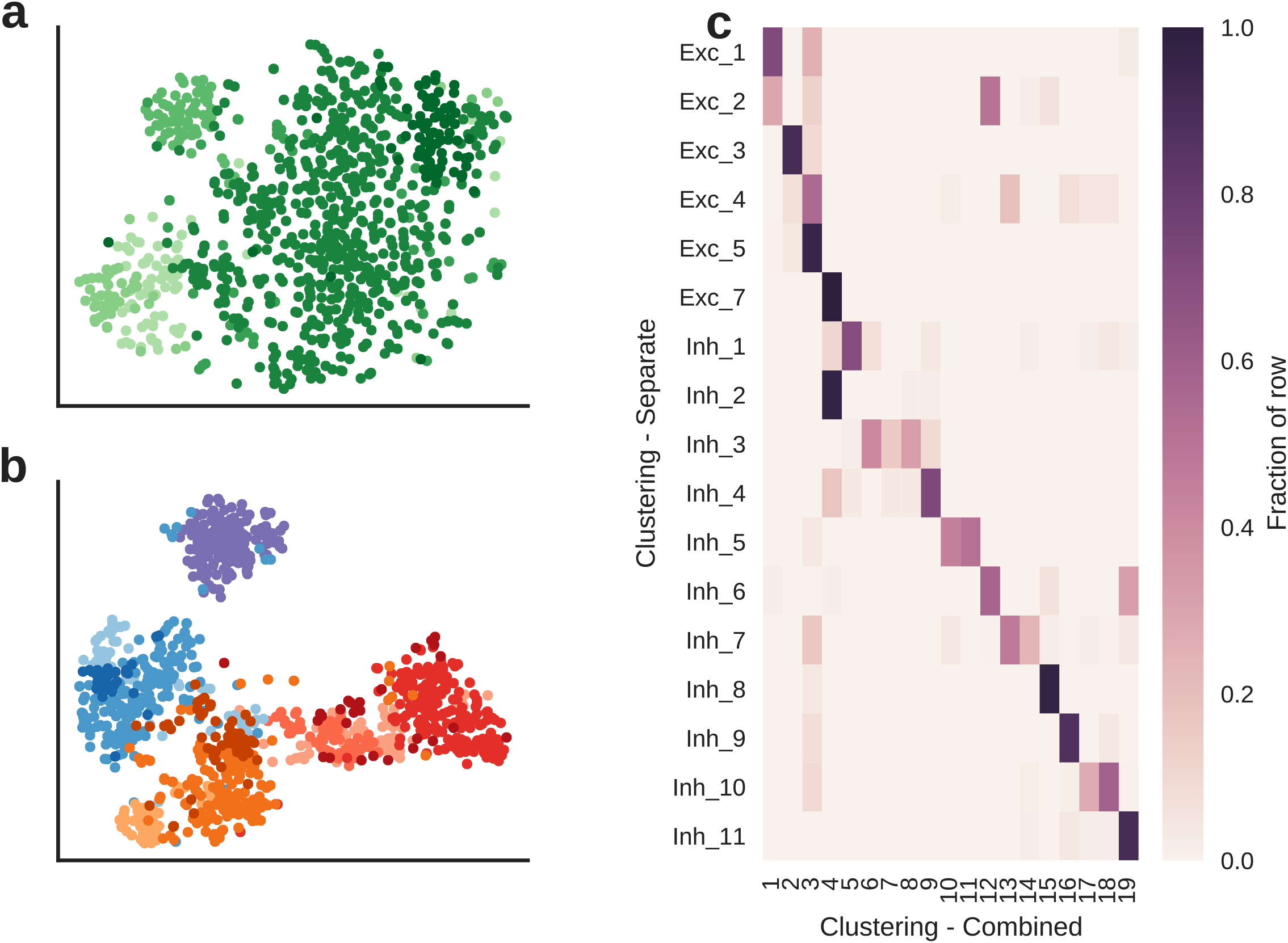
Comparison of separate and combined electrophysiology clustering analyses. (**a**) t-SNE projection of electrophysiological data from only excitatory (spiny) neurons. Colors indicate excitatory electrophysiological clusters. (**b**) t-SNE projection of electrophysiological data from only inhibitory (aspiny) neurons. Colors indicate inhibitory electrophysiological clusters. (**c**) Comparison of electrophysiological clusters obtained by separate analyses of excitatory and inhibitory neurons (rows) and a combined analysis of all cells together (columns). Note that nearly all the excitatory neurons fall into the combined clusters 1–3, while inhibitory neurons generally fall into similar clusters in both the separate and combined cases.

**Supplementary Figure 7:**
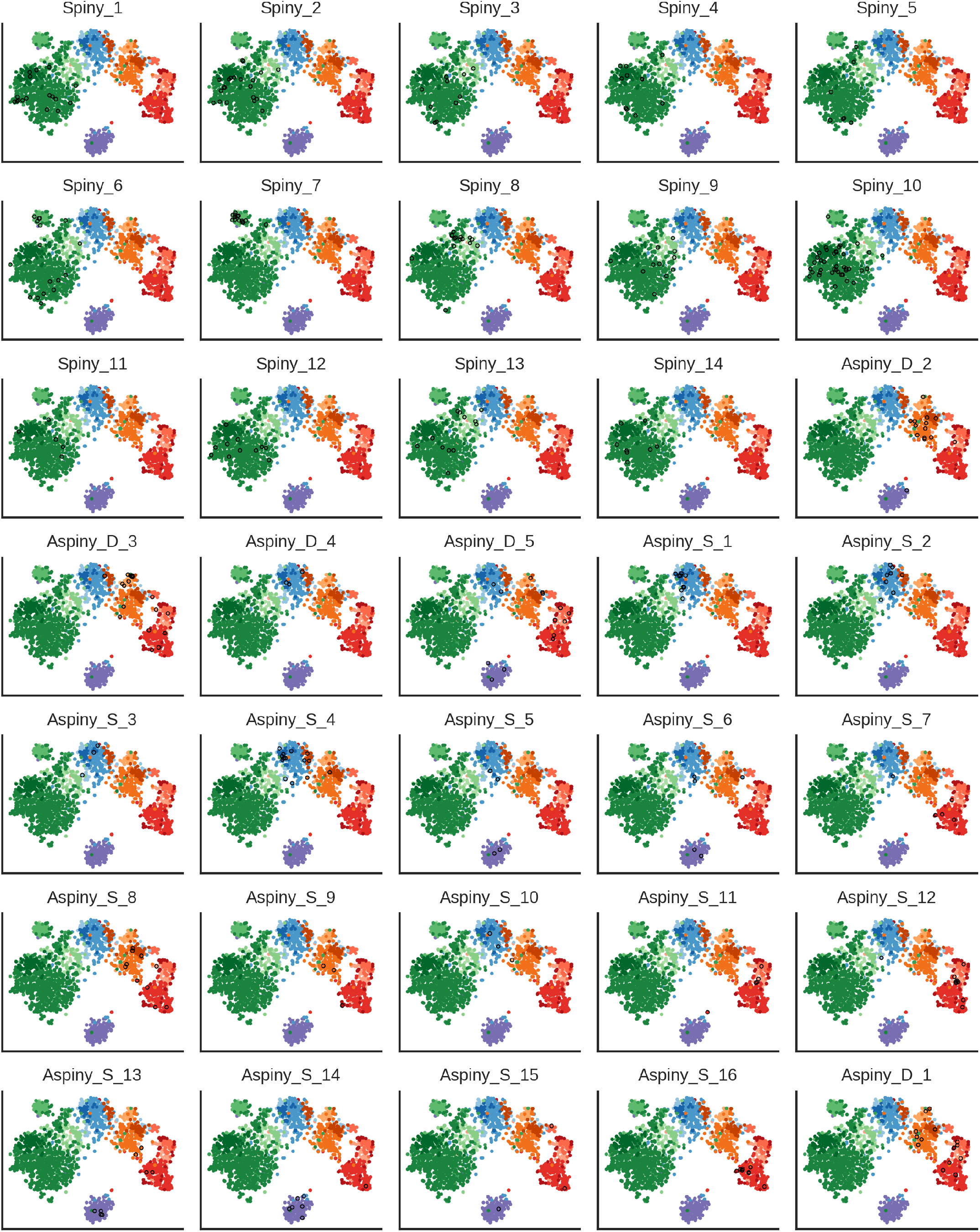
Transgenic lines on the electrophysiological projection. Electrophysiology-based t-SNE plots with cells from different transgenic lines highlighted. Colors indicate electrophysiological cluster labels (see Fig. 2). Cells that were fluorescent-reporter positive with a given transgenic driver are indicated with black circles.

**Supplementary Figure 8:**
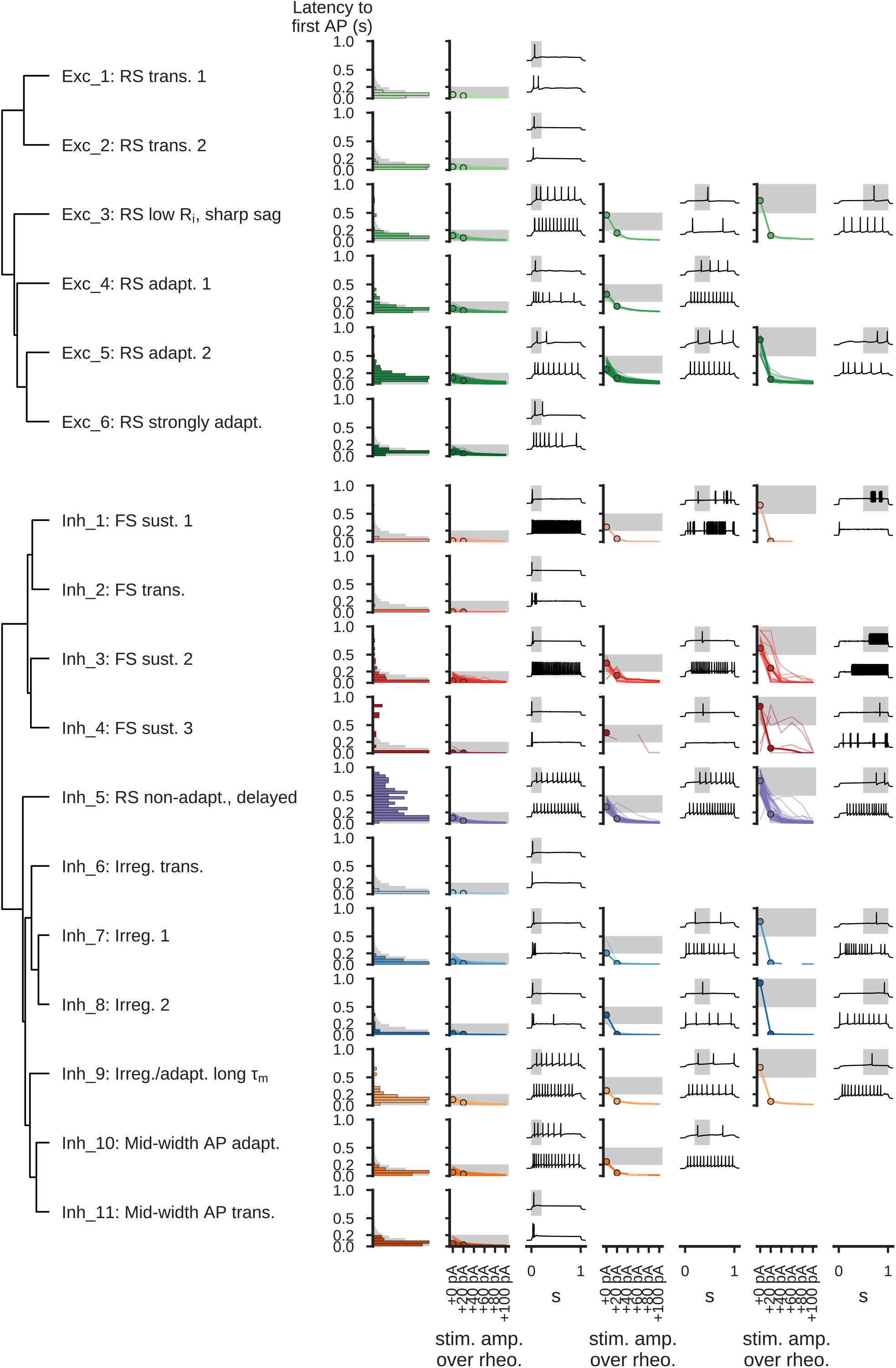
Latency to first action potential. Each row presents information from a different electrophysiological cluster. Dendrogram on left based on distances between cluster centroids (see Fig. 2). Histogram shows the maximum latency to the first spike observed per cell across six long square current steps (from rheobase to rheobase + 100 pA). Gray histogram is all cells, colored histograms are cells of that cluster. Histograms are normalized to their maximum values. On the right, cells in each cluster have been divided into groups based on their maximum latency: 0 s to 0.2 s, 0.2 s to 0.5 s, and 0.5 to 1 s (indicated by shaded regions on line plots and upper example traces). Line plots show how the latency per sweep changes as the stimulus amplitude is varied. Example traces show a representative cell for each cluster/category combination; upper trace is the one with the longest latency from a cell, lower trace is the next longest from the same cell. Selected examples are indicated on the line plots by thicker lines and circles.

**Supplementary Figure 9:**
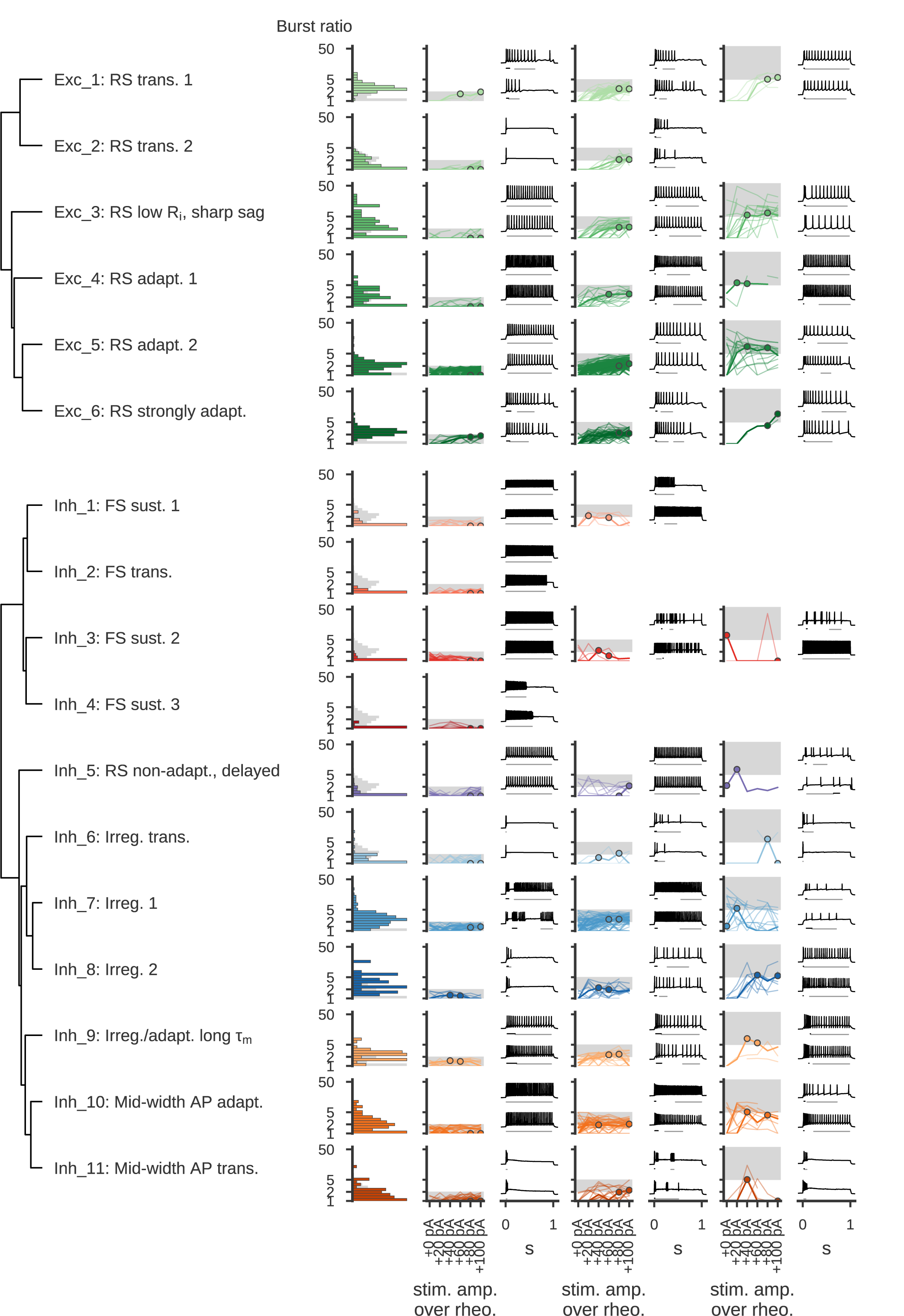
Bursting firing patterns. Each row presents information from a different electrophysiological cluster. Dendrogram on left based on distances between cluster centroids (see Fig. 2). Histogram shows the maximum burst ratio observed per cell across six long square current steps (from rheobase to rheobase + 100 pA). The burst ratio is defined as the firing rate of the fastest segment divided by the firing rate of the segment(s) with the most action potentials (Methods); the median across segments was used for the latter in the case of ties. Gray histogram is all cells, colored histograms are cells of that cluster. Histograms are normalized to their maximum values. On the right, cells in each cluster have been divided into groups based on their maximum burst ratio: 1 to 2, 2 to 5, and 5 or more (indicated by shaded regions on line plots). Line plots show how the maximum burst ratio per sweep changes as the stimulus amplitude is varied. Example traces show a representative cell for each cluster/category combination; upper trace is the one with the highest burst ratio, lower trace is the next highest from the same cell. Lines underneath the traces indicate the segment with highest firing rate (black) and the segment(s) with the most action potentials (gray). Selected examples are indicated on the line plots by thicker lines and circles.

**Supplementary Figure 10:**
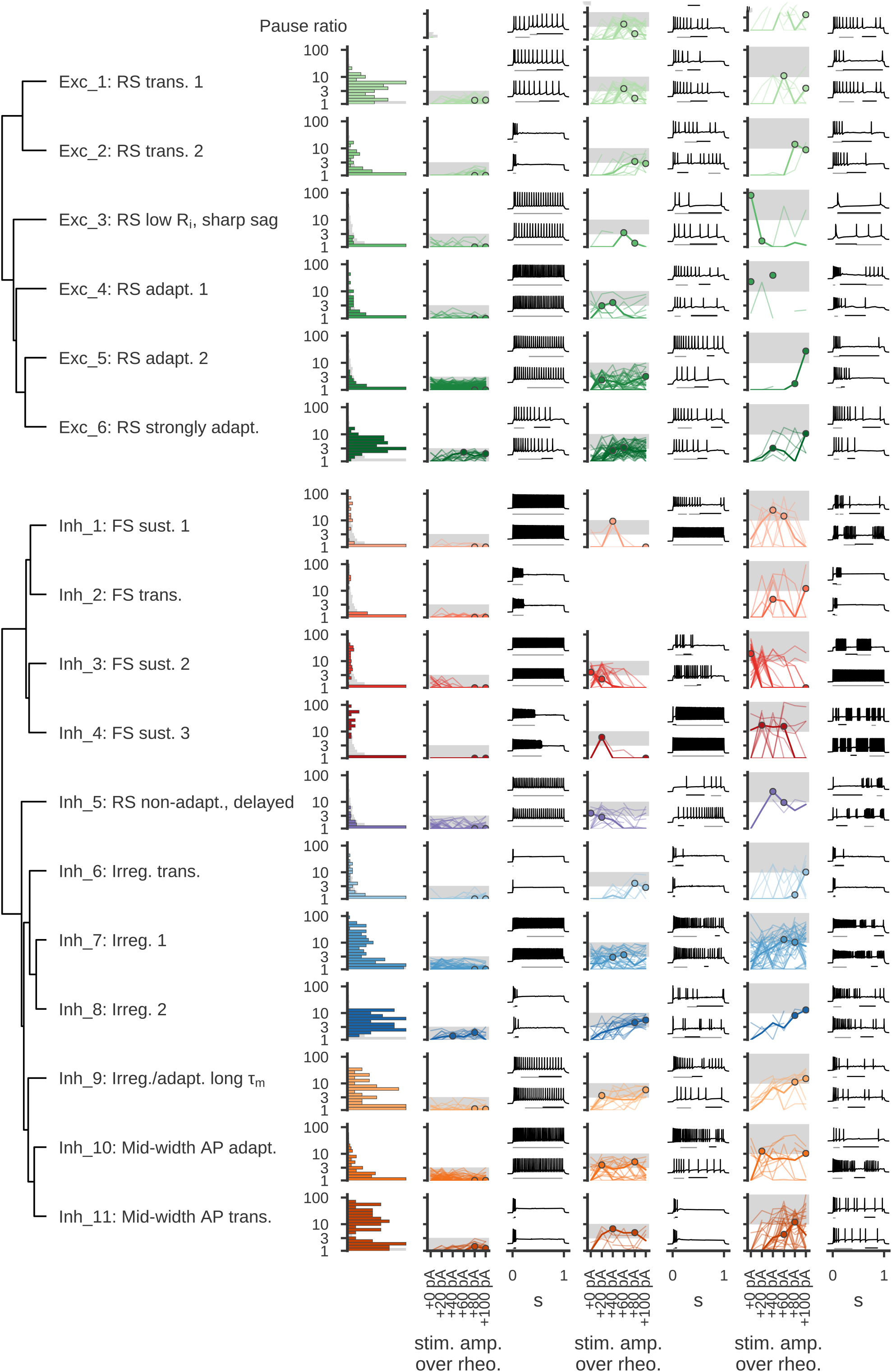
Pausing firing patterns. Each row presents information from a different electrophysiological cluster. Dendrogram on left based on distances between cluster centroids (see Fig. 2). Histogram shows the maximum pause ratio observed per cell across six long square current steps (from rheobase to rheobase + 100 pA). The pause ratio is defined as the average interspike interval duration of the segment with the slowest firing divided by the average interspike interval of the segment(s) with the most action potentials (Methods); the median across segments was used for the latter in the case of ties. Gray histogram is all cells, colored histograms are cells of that cluster. Histograms are normalized to their maximum values. On the right, cells in each cluster have been divided into groups based on their maximum pause ratio: 1 to 3, 3 to 10, and 10 or more (indicated by shaded regions on line plots). Line plots show how the maximum burst ratio per sweep changes as the stimulus amplitude is varied. Example traces show a representative cell for each cluster/category combination; upper trace is the one with the highest burst ratio, lower trace is the next highest from the same cell. Lines underneath the traces indicate the segment with highest firing rate (black) and the segment(s) with the most action potentials (gray). Selected examples are indicated on the line plots by thicker lines and circles.

**Supplementary Figure 11:**
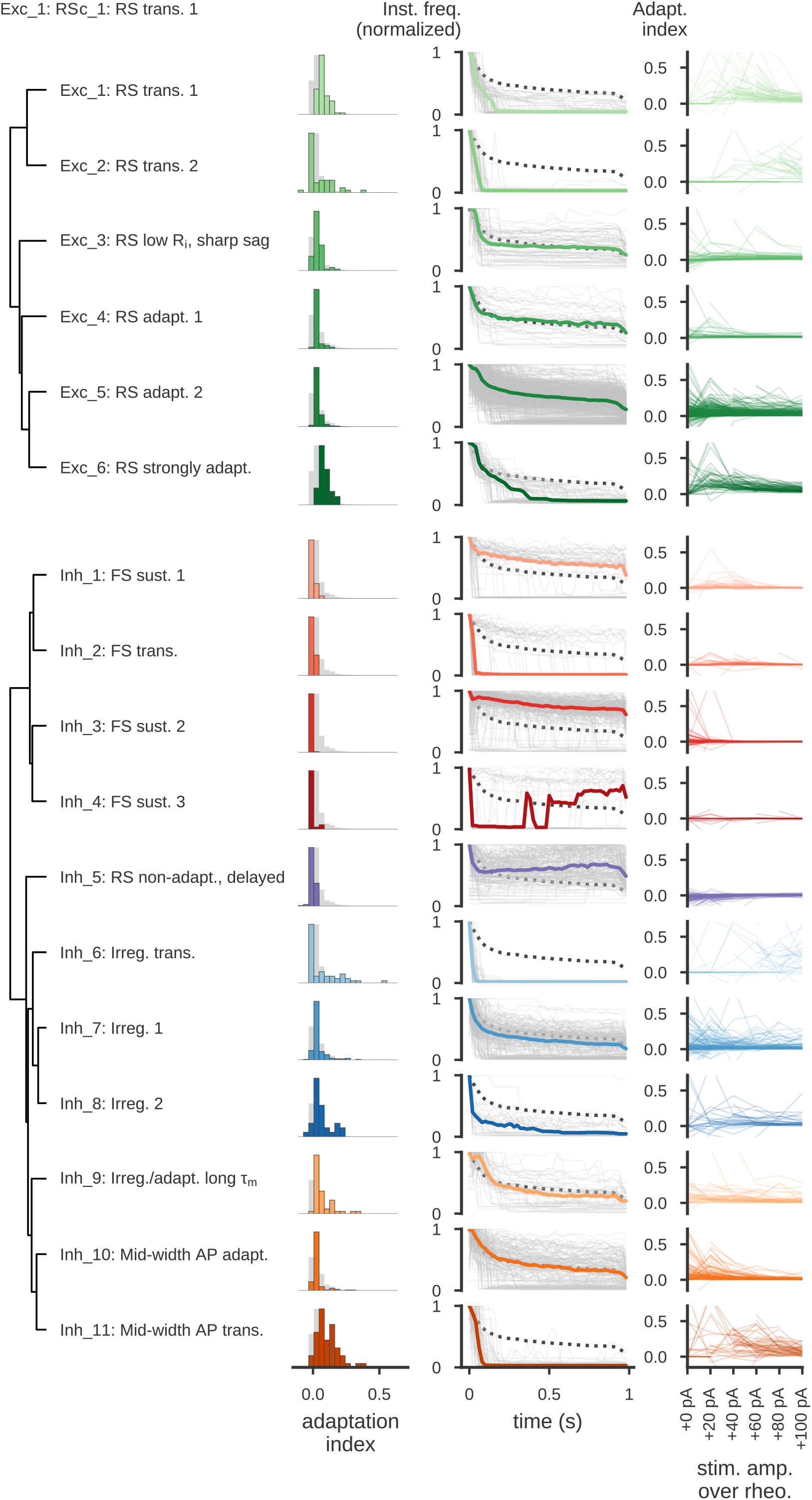
Firing frequency adaptation. Each row presents information from a different electrophysiological cluster. Dendrogram on left based on distances between cluster centroids (see Fig. 2). Histogram shows the median adaptation index observed per cell across six long square current steps (from rheobase to rheobase + 100 pA). Center plots show the firing rates (calculated in 20 ms bins) from the sweep at the median adaptation index, normalized to the highest firing rate of the sweep. All cells from the cluster are shown as gray lines, the cluster medians are shown as the thick colored lines, and the grand median across all cells is shown as a dotted line. Note that the non-monotonic median of Inh_4 is due to many cells in that cluster exhibiting pauses in firing (where the instantaneous firing rate falls to zero) toward the start of the stimulus period. Right plots show how the adaptation index varied across six long square current steps (from rheobase to rheobase + 100 pA).

**Supplementary Figure 12:**
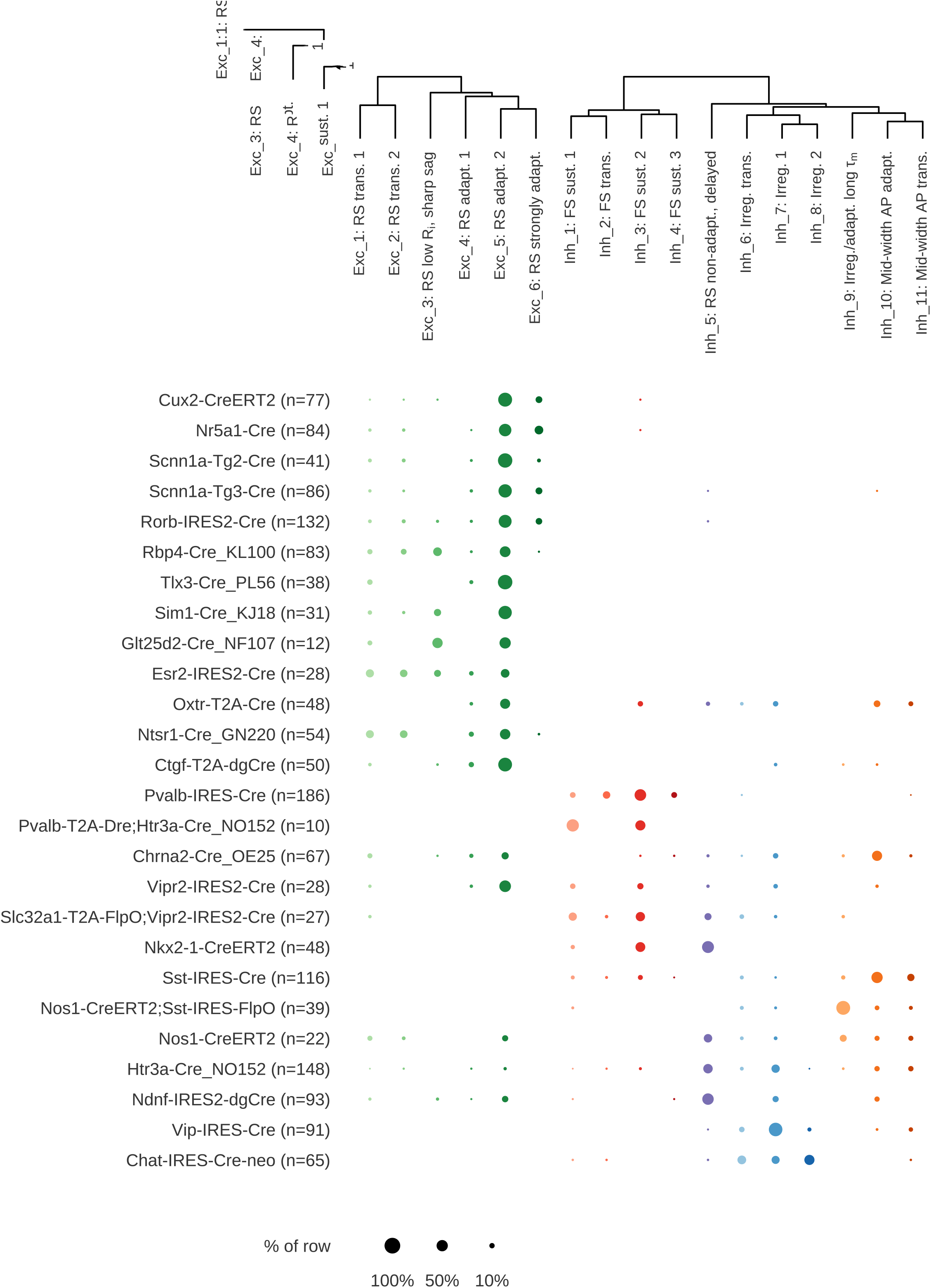
Transgenic lines and electrophysiological types. Fraction of cells from each transgenic line examined (rows) that fall into each e-type (columns). Dot size indicates the fraction of the row falling into a given column, and color indicates e-type.

**Supplementary Figure 13:**
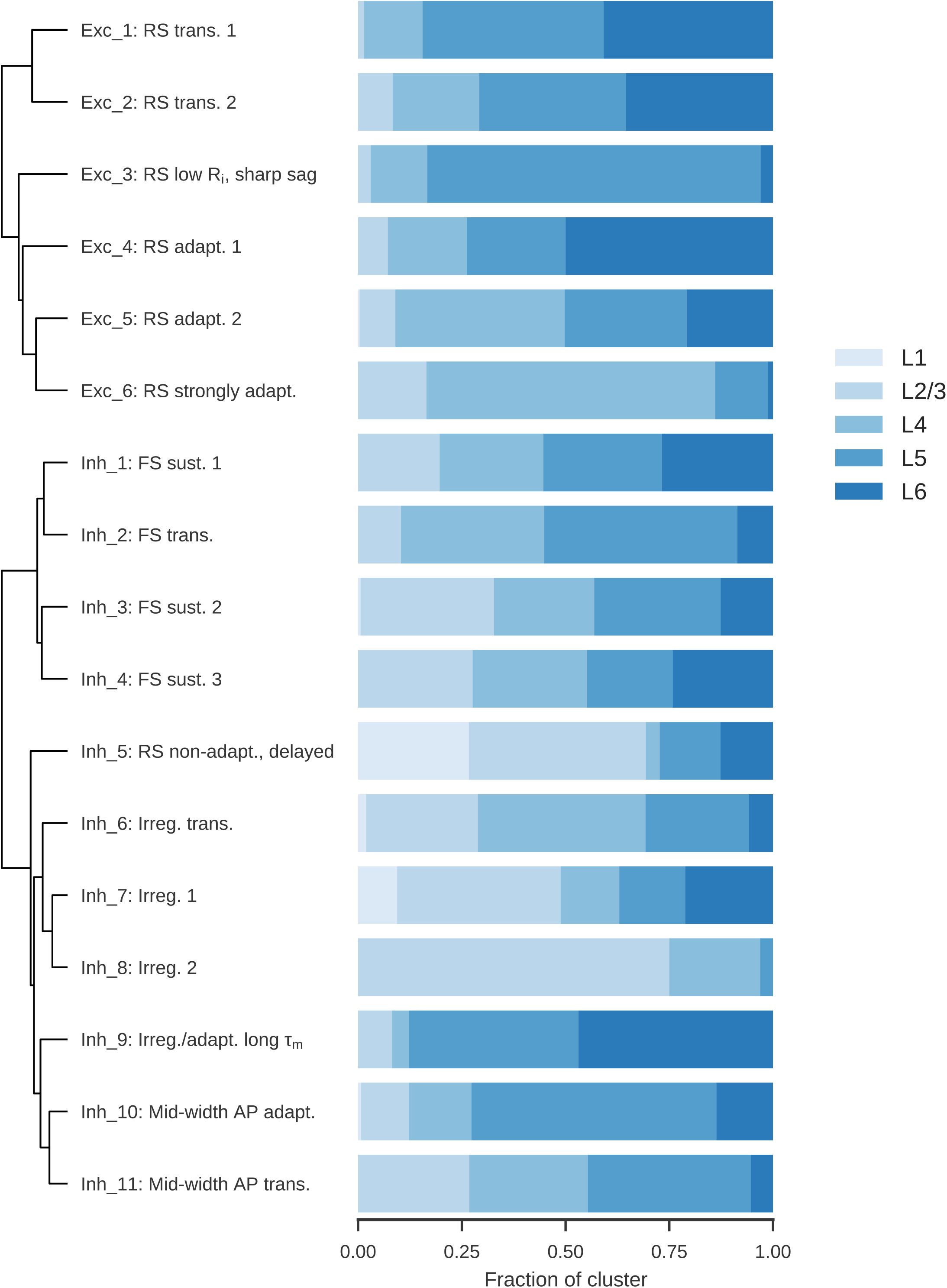
Electrophysiological types and cortical layers. Distribution of cells from each electrophysiological cluster across the cortical layers.

**Supplementary Figure 14:**
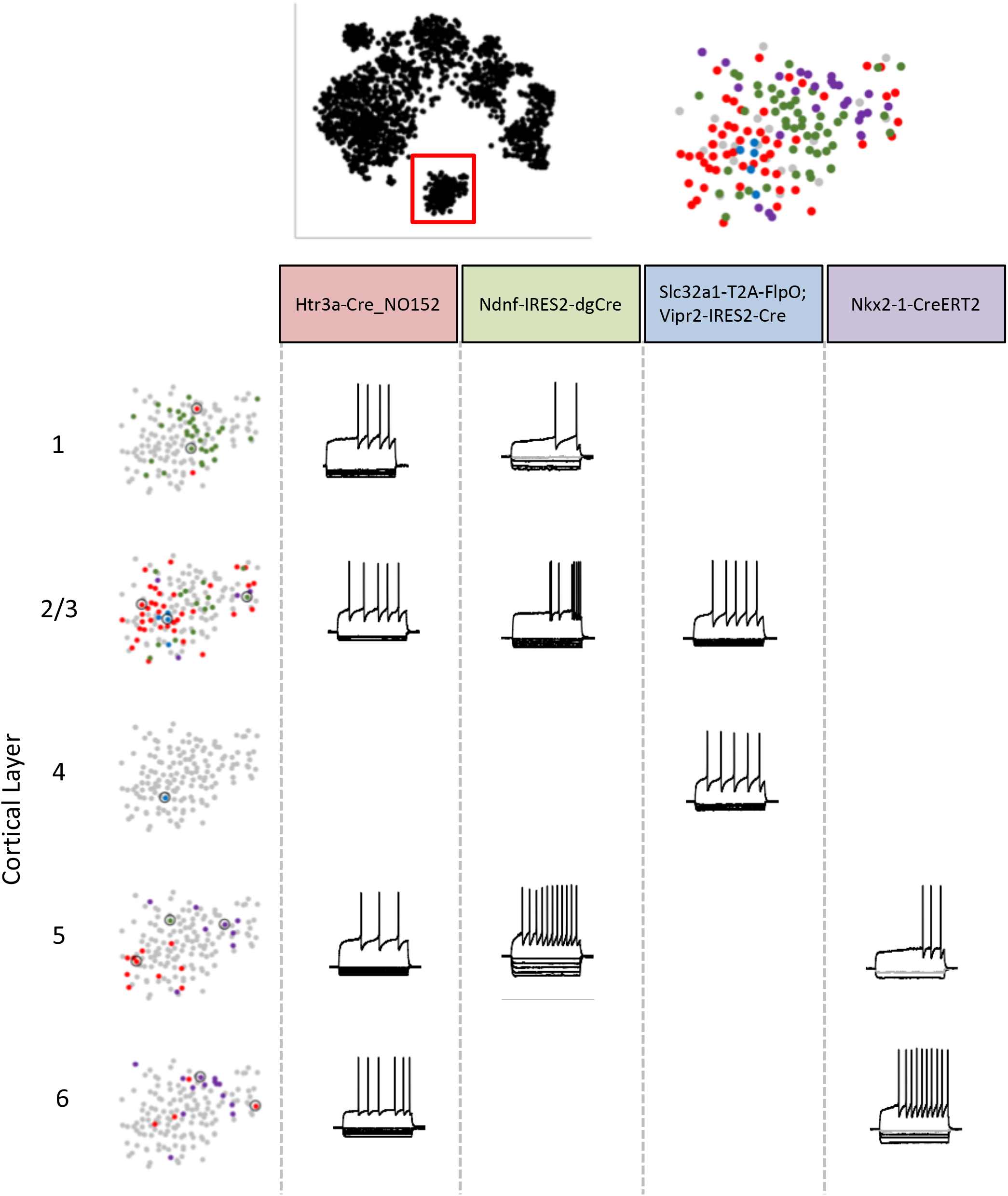
Inh_5 electrophysiology cluster t-SNE distribution by Cre line and layer. Focused analysis of the region of the electrophysiology t-SNE feature-space where e-type Inh_5 shows patterns based on Cre line and layer distribution. Cells within this cluster are divided between a superficial group, dominated by Ndnf+ cells in layer 1 and Htr3a+ cells in layer 2/3. Deeper cells within this cluster are either Htr3a+, with longer action potentials and more regular firing, or a mixture of Htr3a+ and Nkx2.1+ neurons with faster action potentials and more irregular firing patterns. Note that these different cells are all found within the large southern inhibitory island of the t-SNE plot (magnified on the top right hand).

**Supplementary Figure 15:**
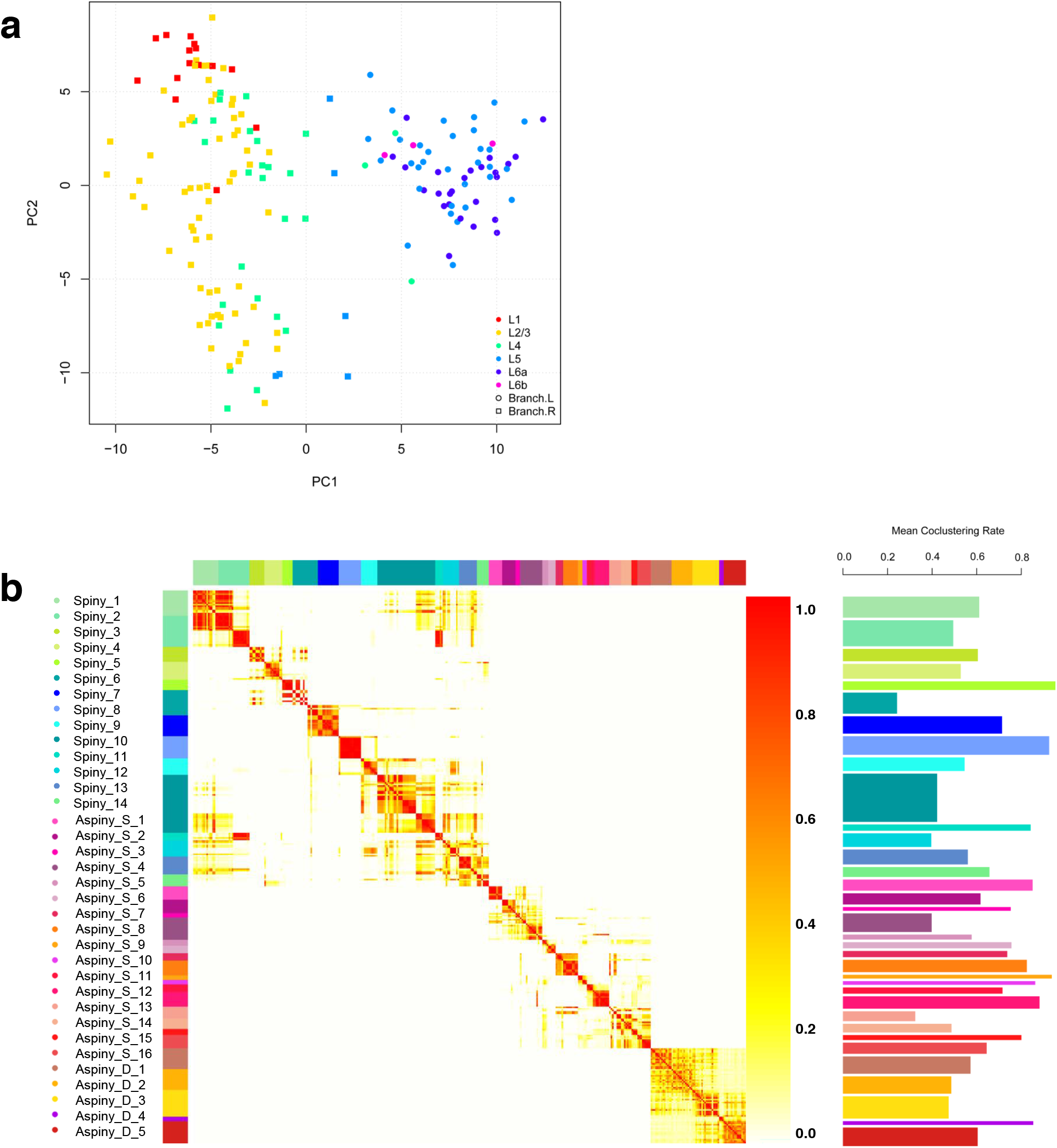
(**a**) Partitioning of Aspiny neurons into superficial and deep populations. Hierarchical clustering of aspiny neurons with all aspiny neurons and features shows a clear division between two groups, which correspond to superficial and deep layers. The Left/Right partition of the clustering tree is shown in PCA domain. (**b**) Co-clustering Analysis. Co-clustergram (top) and mean co-clustering rate (bottom). 100 runs of clustering with 90% subsampling in 10fold cross validation manner shows robustness of each cluster.

**Supplementary Figure 16:**
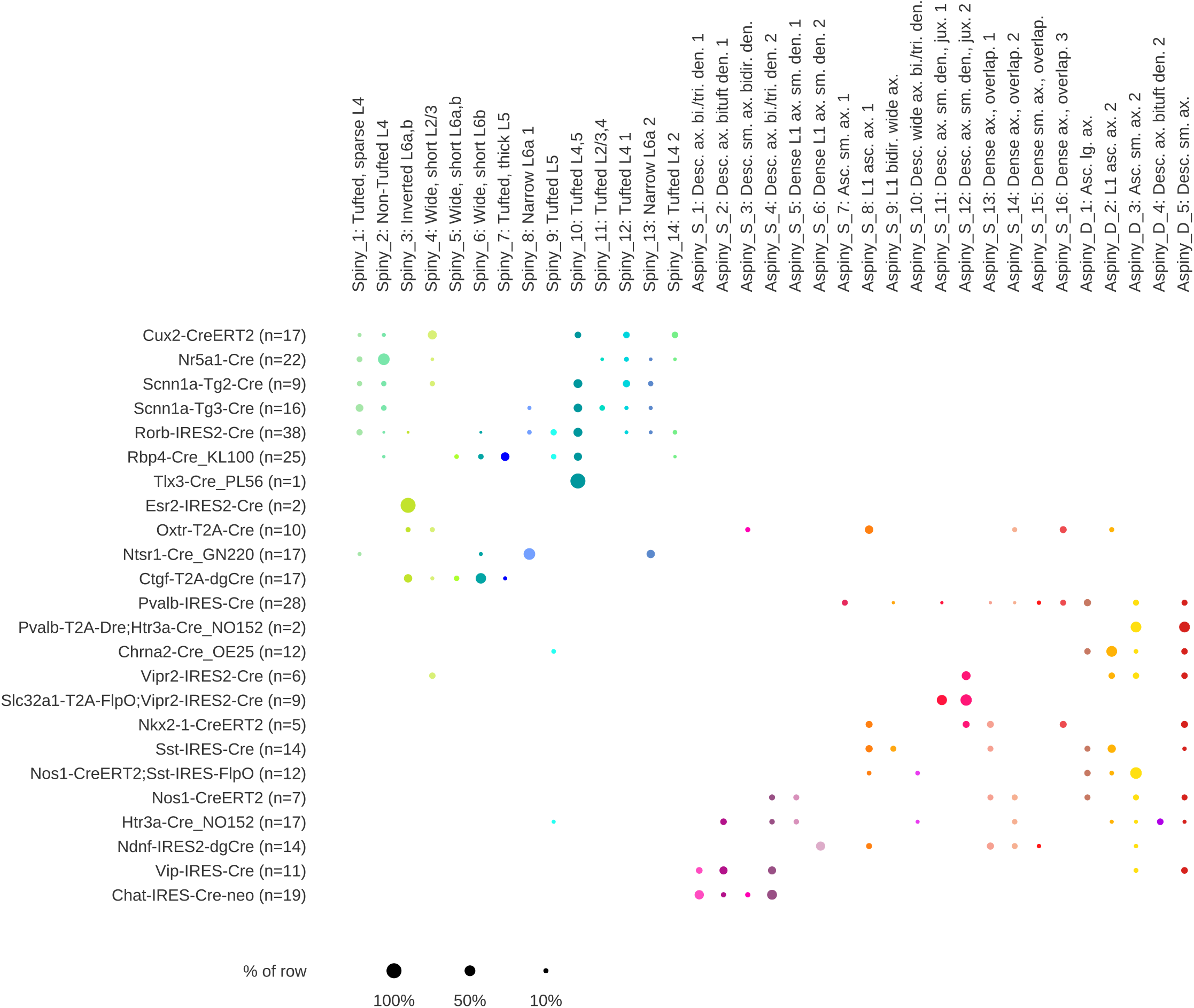
Transgenic lines and morphological types. Fraction of cells from each transgenic line examined (rows) that fall into each m-type (columns). Dot size indicates the fraction of the row falling into a given column, and color indicates m-type.

**Supplementary Figure 17:**
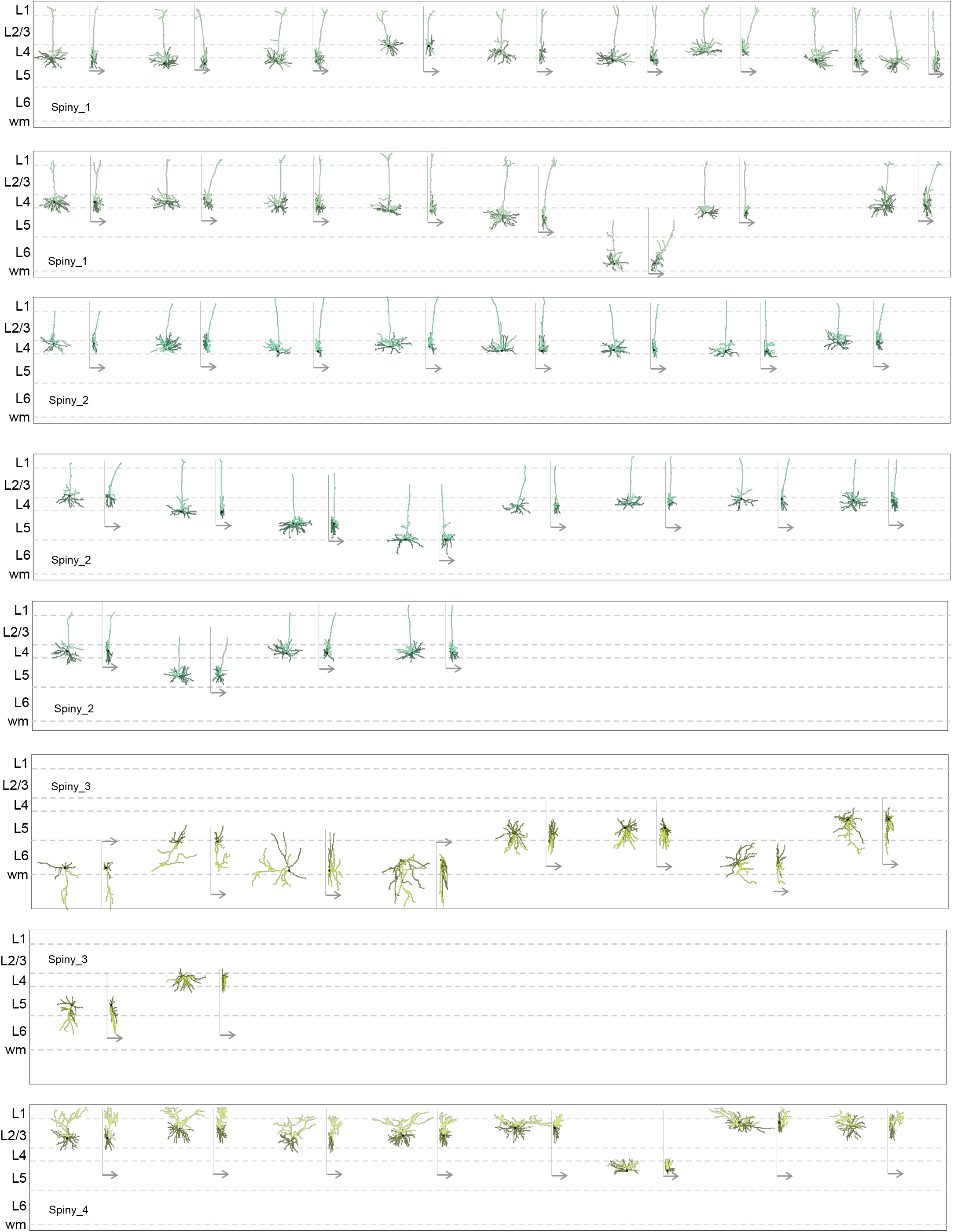
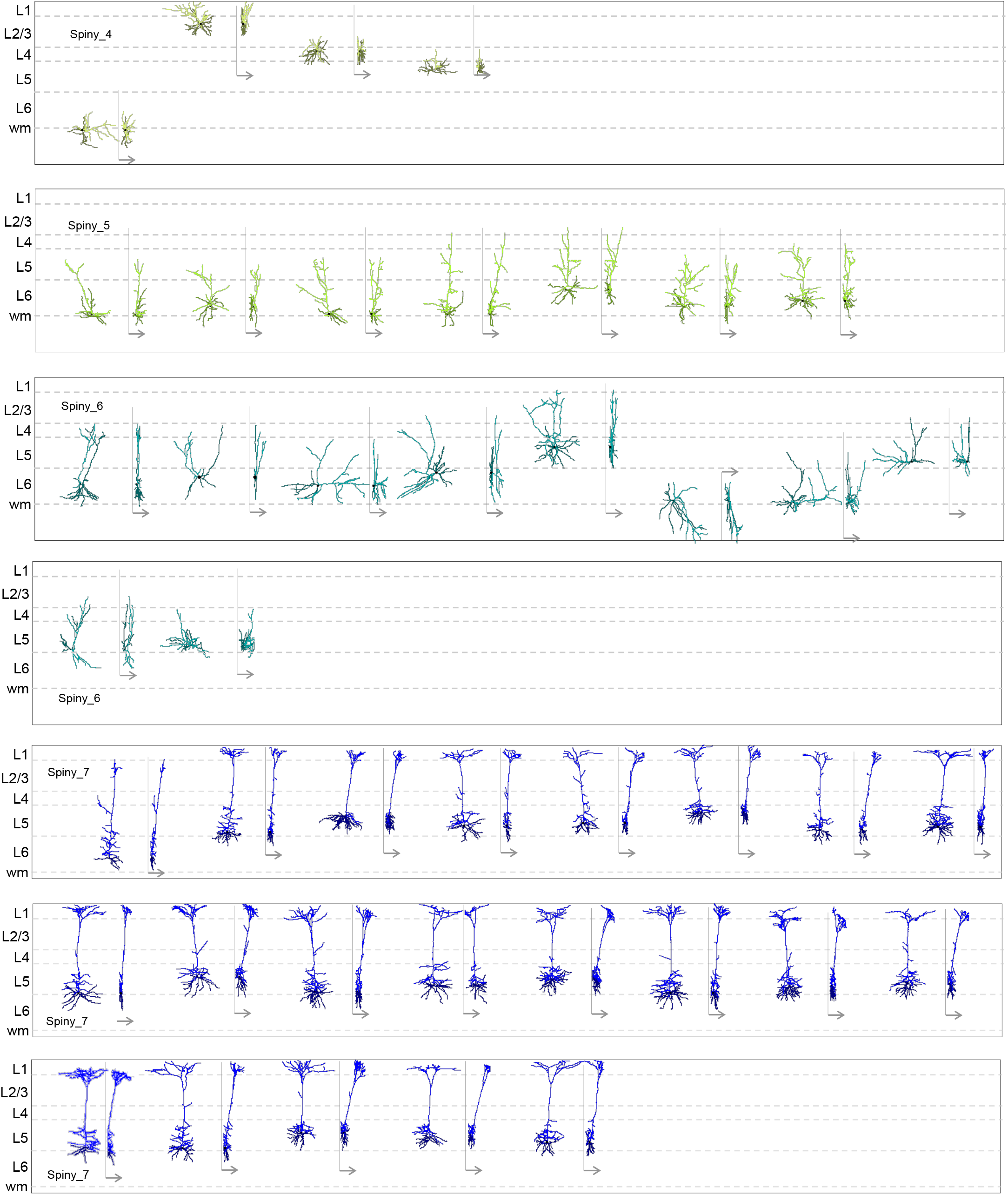
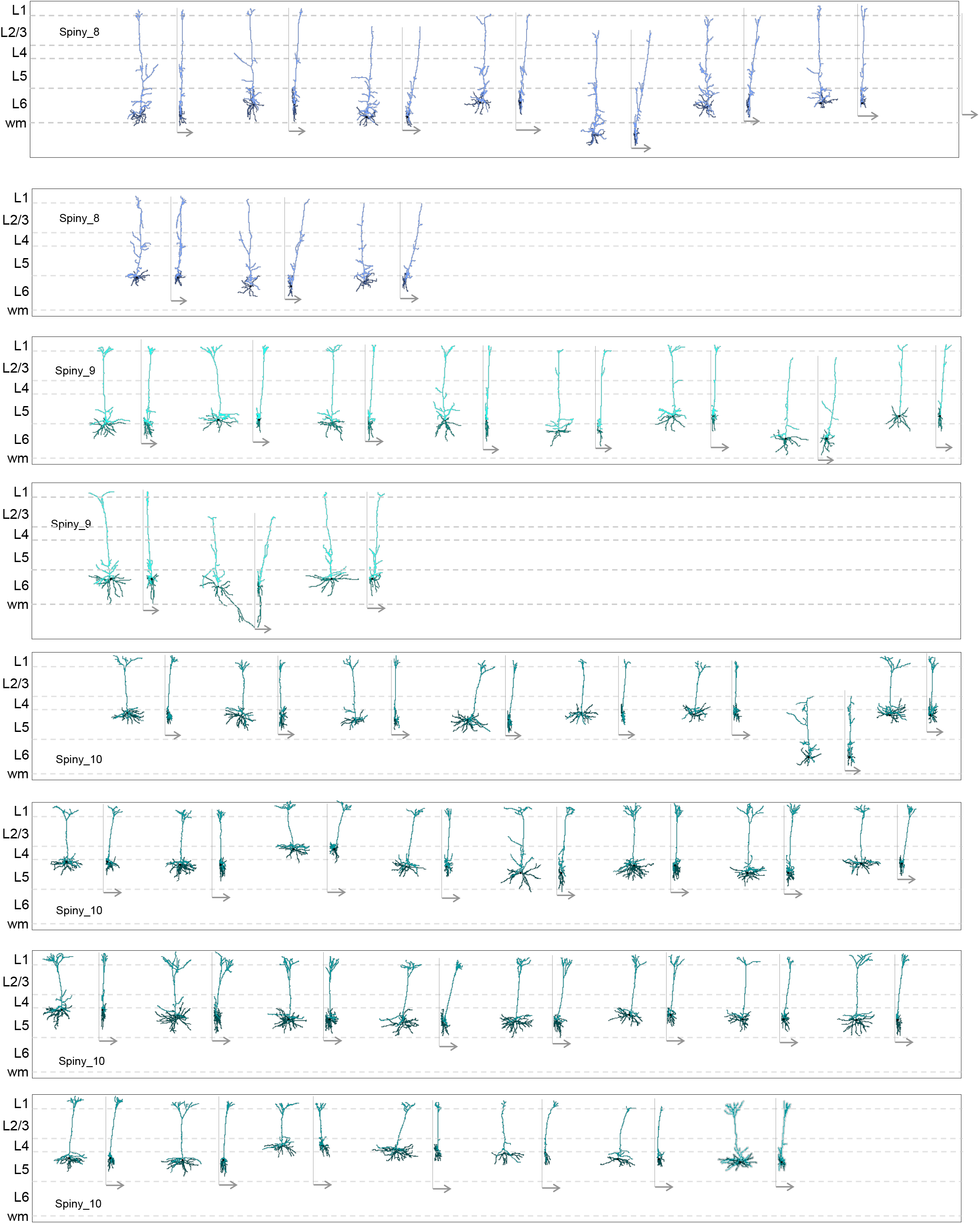
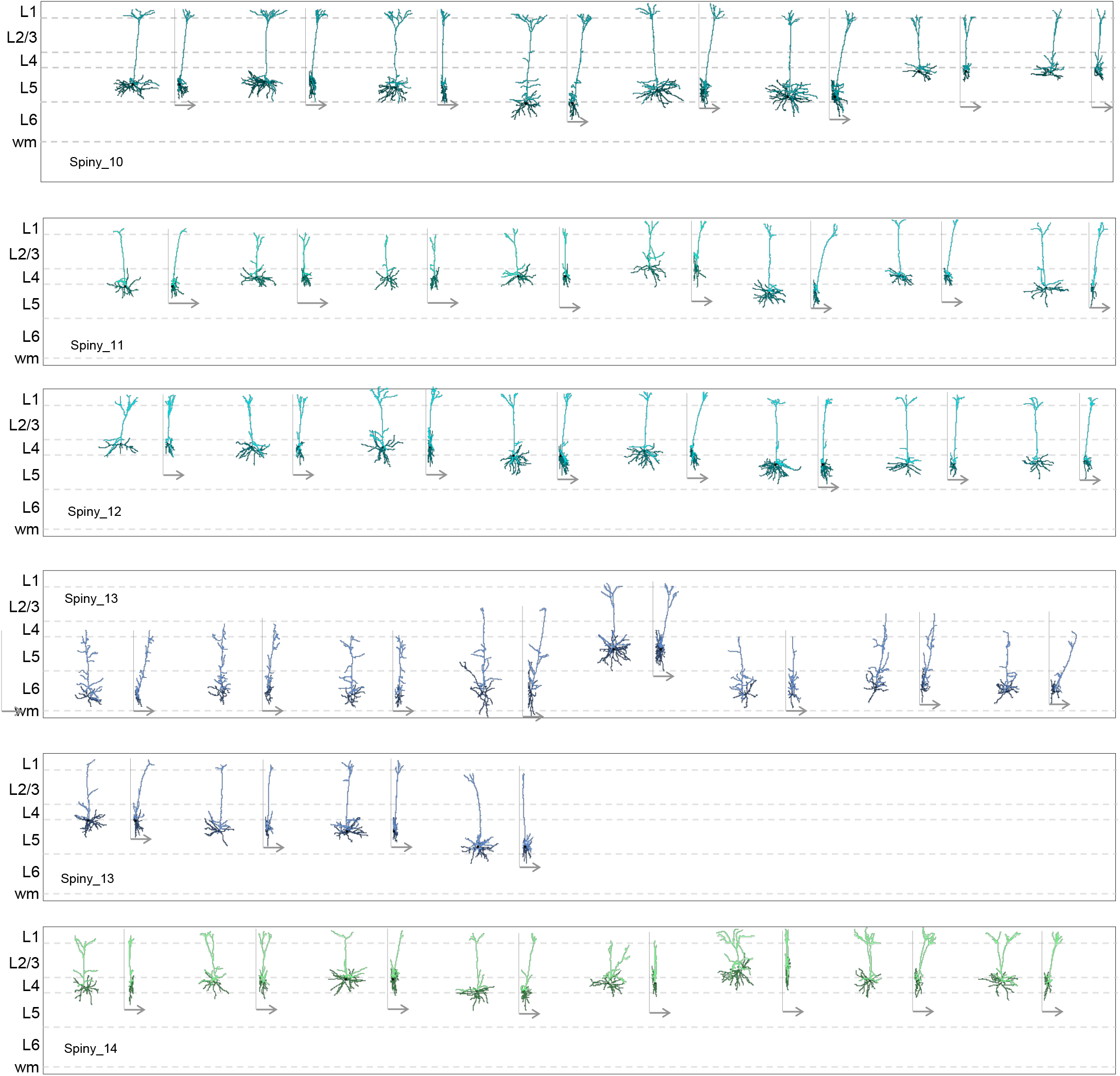
Spiny neuron morphology. All 3D reconstructions used in our quantitative analysis are displayed in their approximate laminar location within the cortical thickness. Two views of each reconstruction are shown. For each cell, the XY dimension view is on the left and the YZ dimension view is on the right and has an arrow indicating the Z dimension (in this case, Z is into the depth of the coronal slice). Reconstructions are color-coded by cluster. Apical dendrites appear in the lighter hue and basal dendrites in the darker hue. We reconstructed neurons with intact, apical dendrites and healthy, relatively intact basal dendrites. Neurons were sampled from cortical layers 2/3–6b and using a range of layer-selective Cre lines in mouse primary visual cortex. A wide diversity of morphologies can be observed. (N = 199).

**Supplementary Figure 18:**
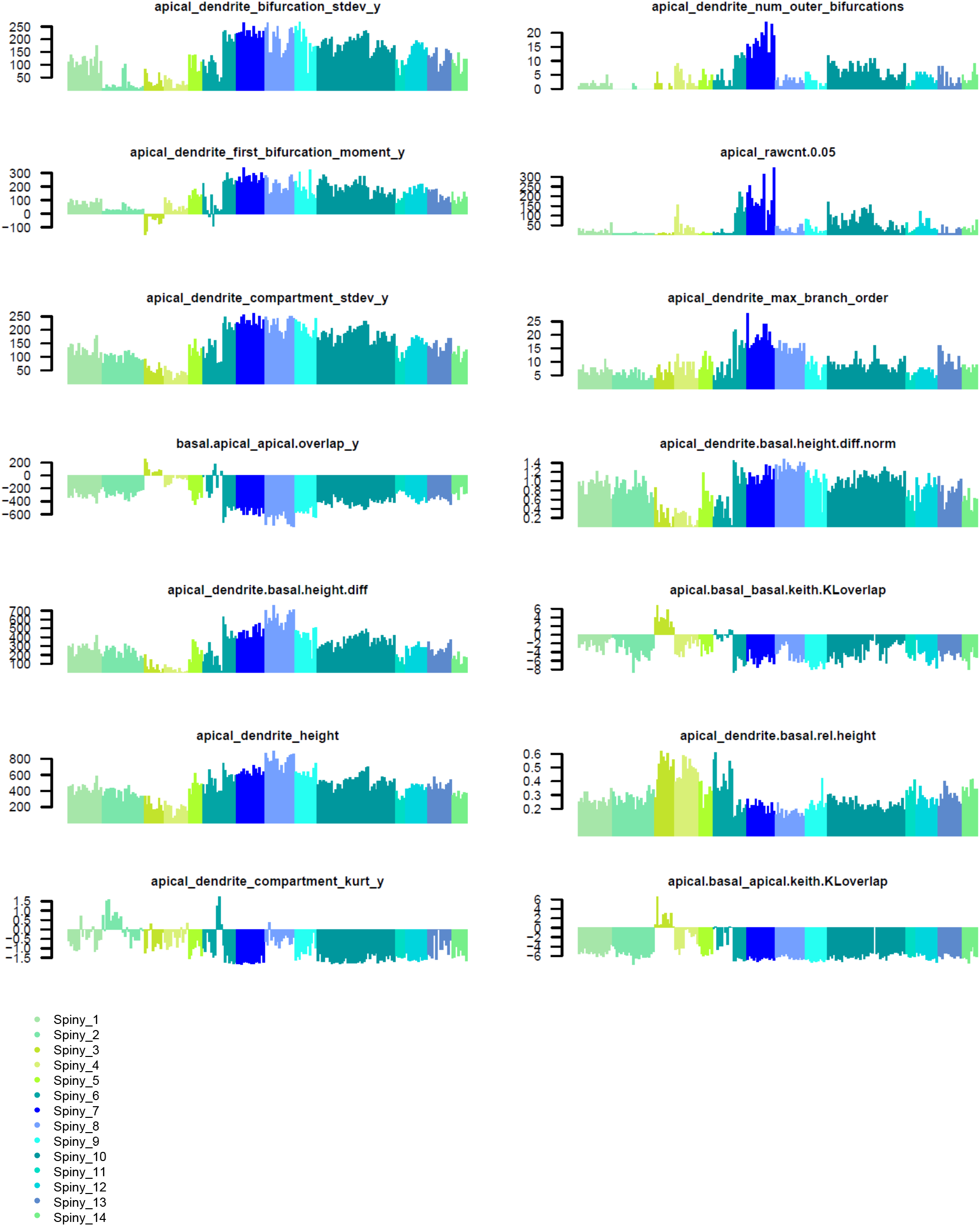
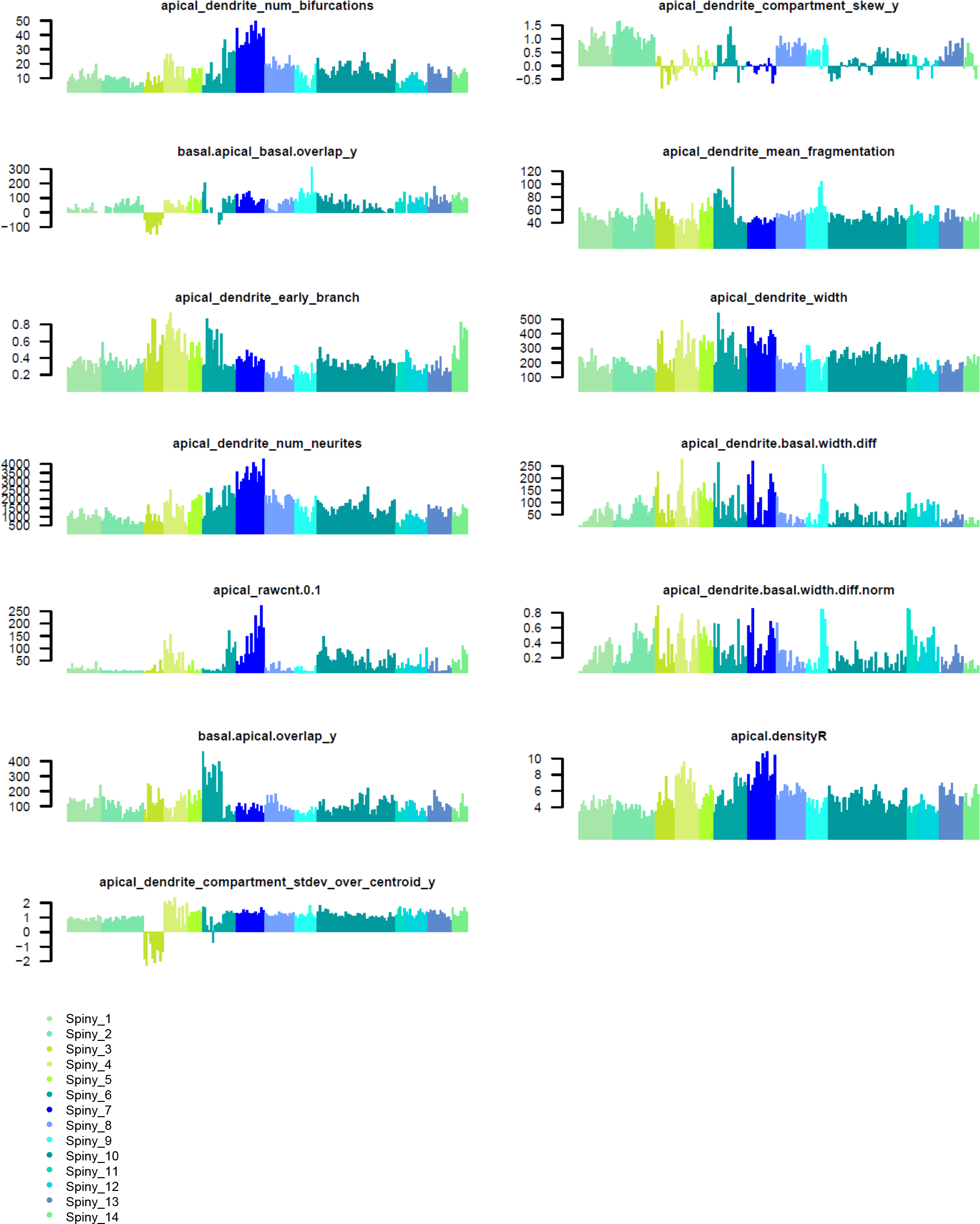
Spiny neuron morphology features by cluster. Based on 3D reconstructions of the apical and basal dendrites, we extracted numerous morphological features from each neuron. Population histograms of each of 27 representative features are shown. Many of the features vary substantially across m-types (N = 199).

**Supplementary Figure 19:**
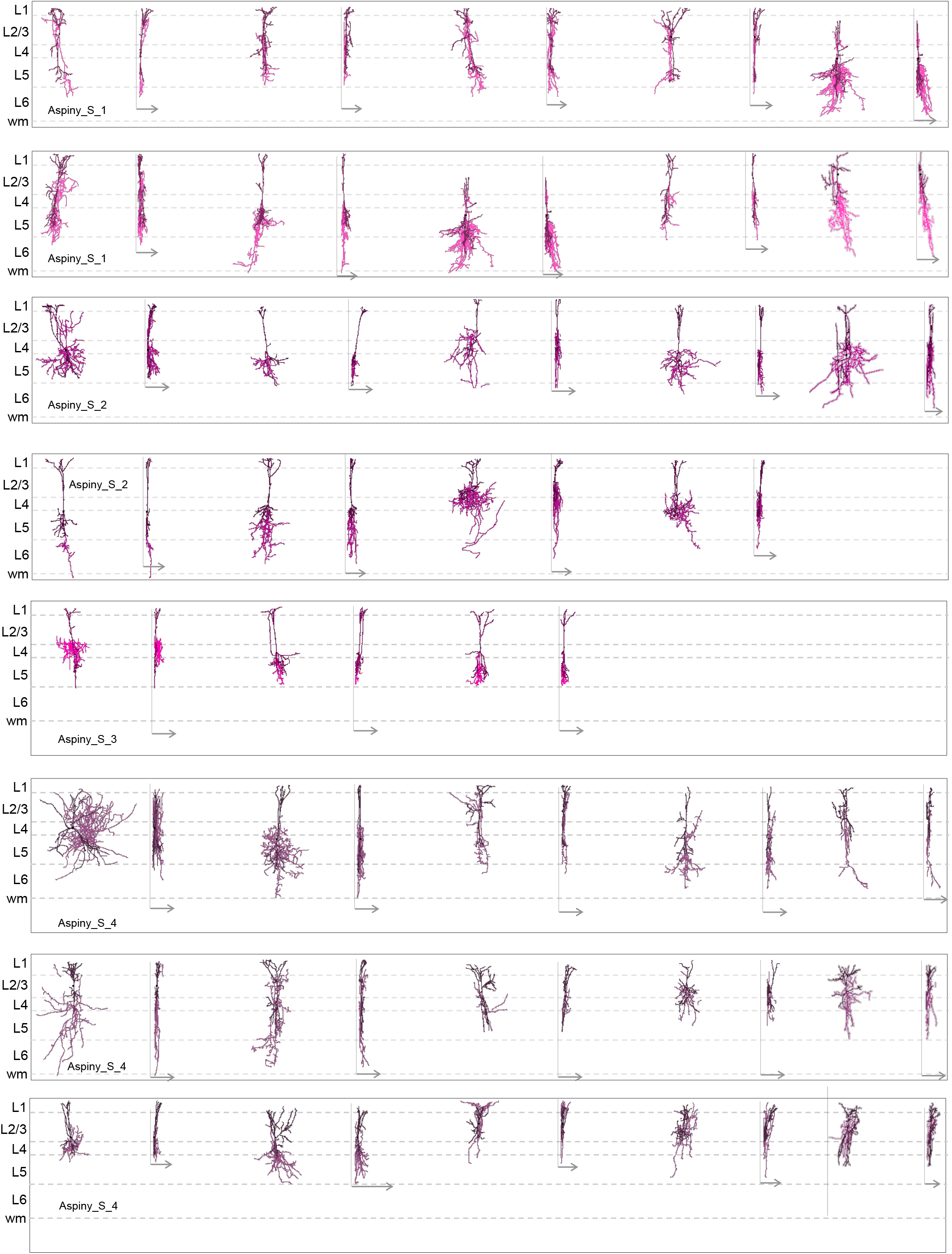
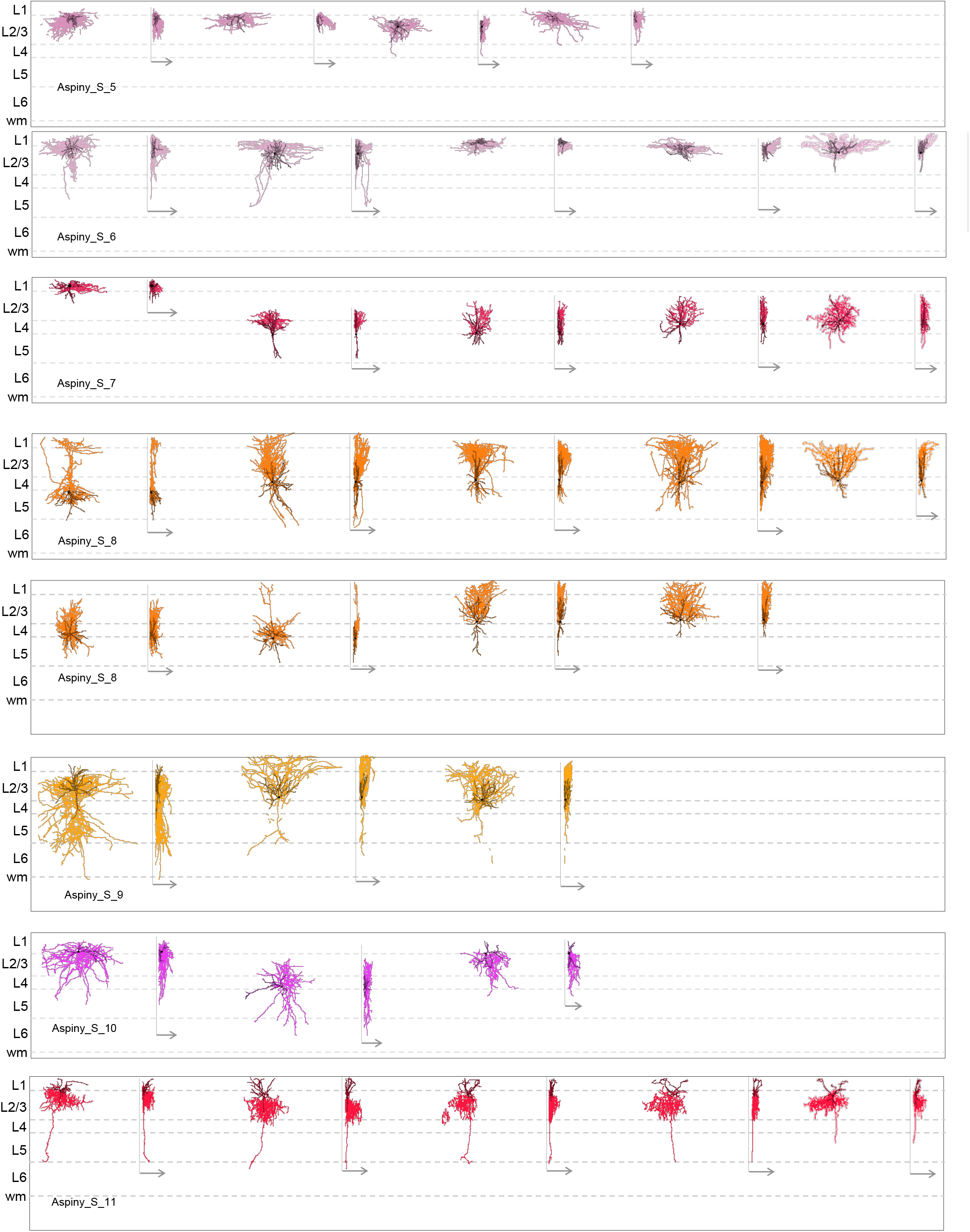
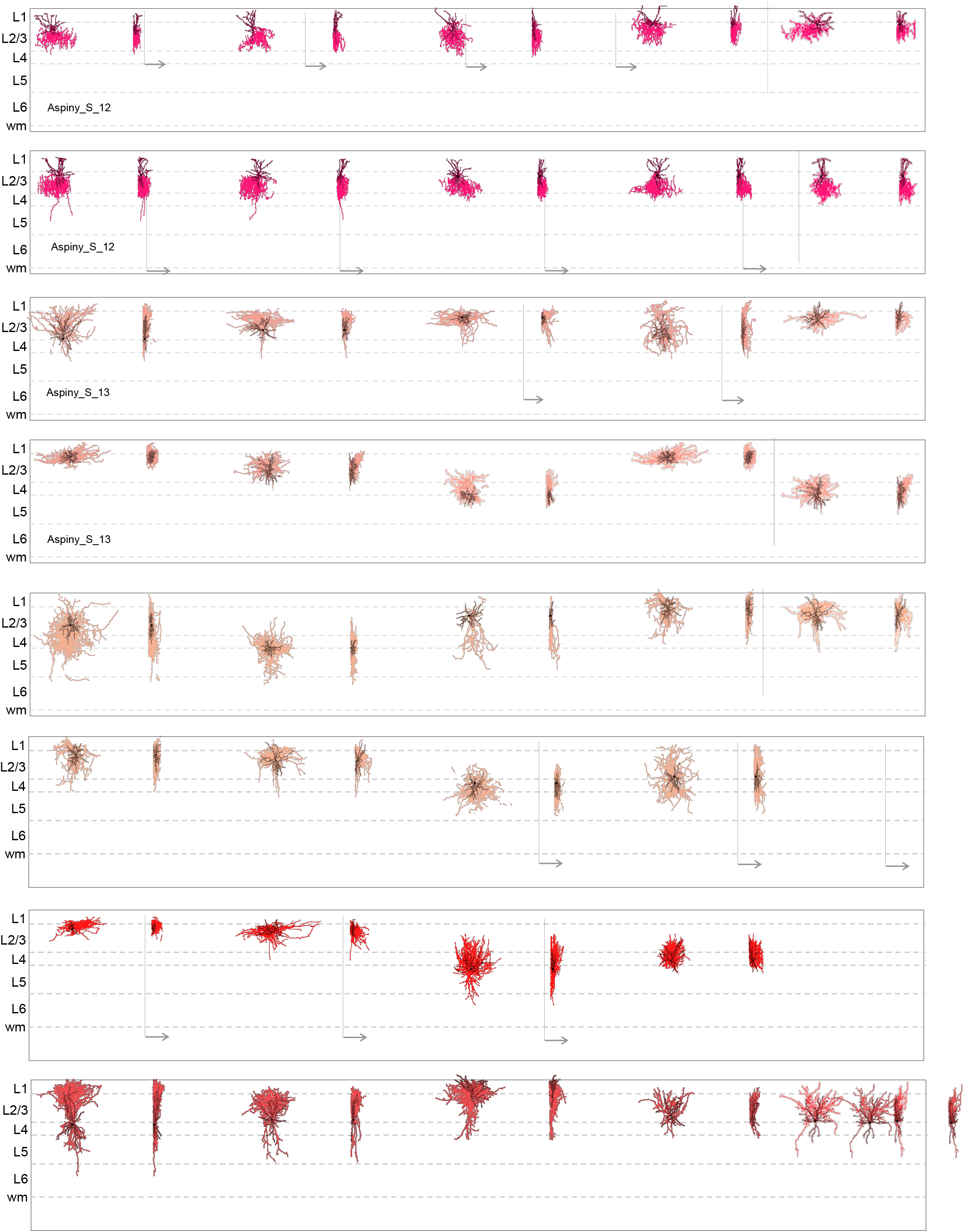
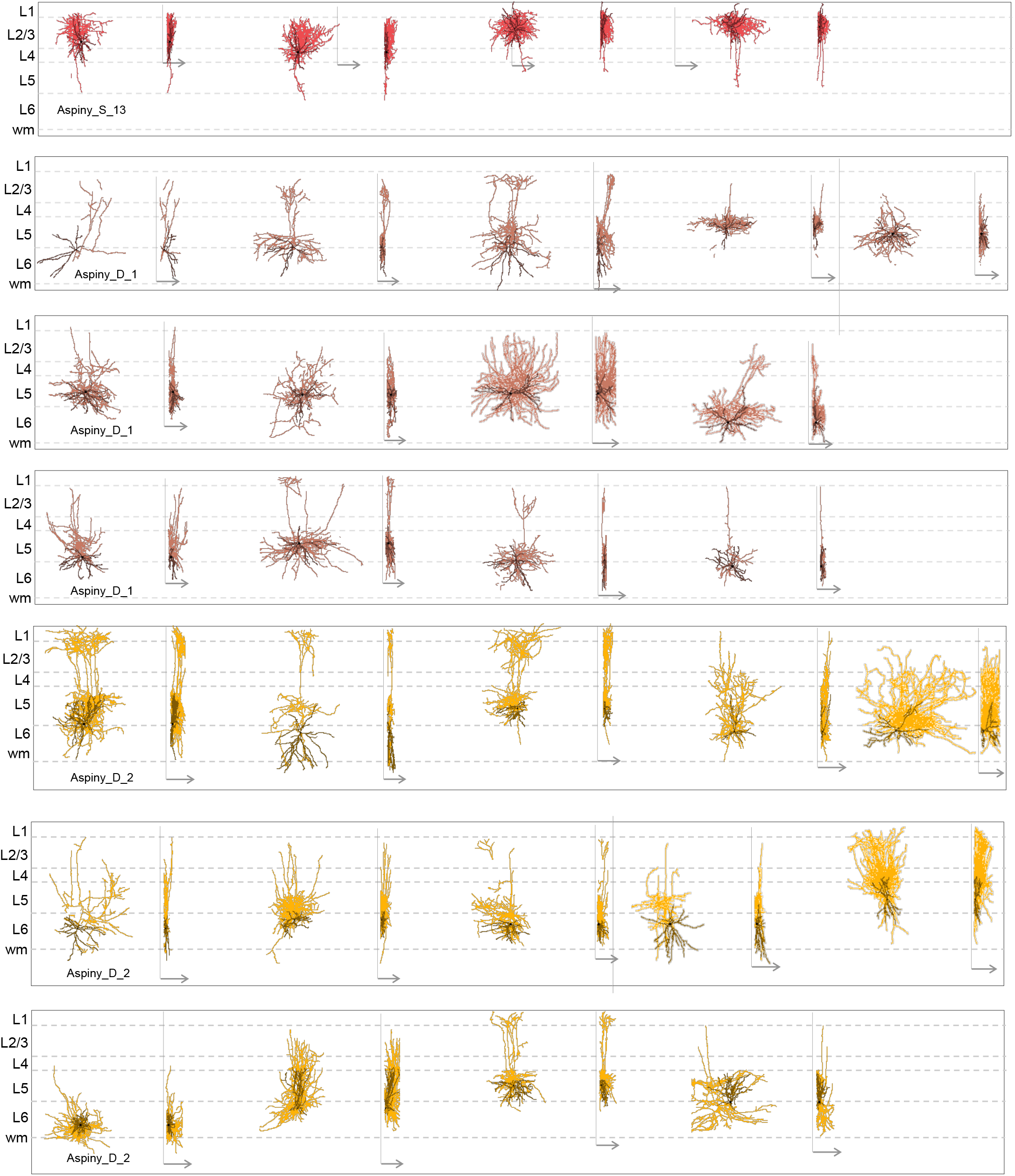
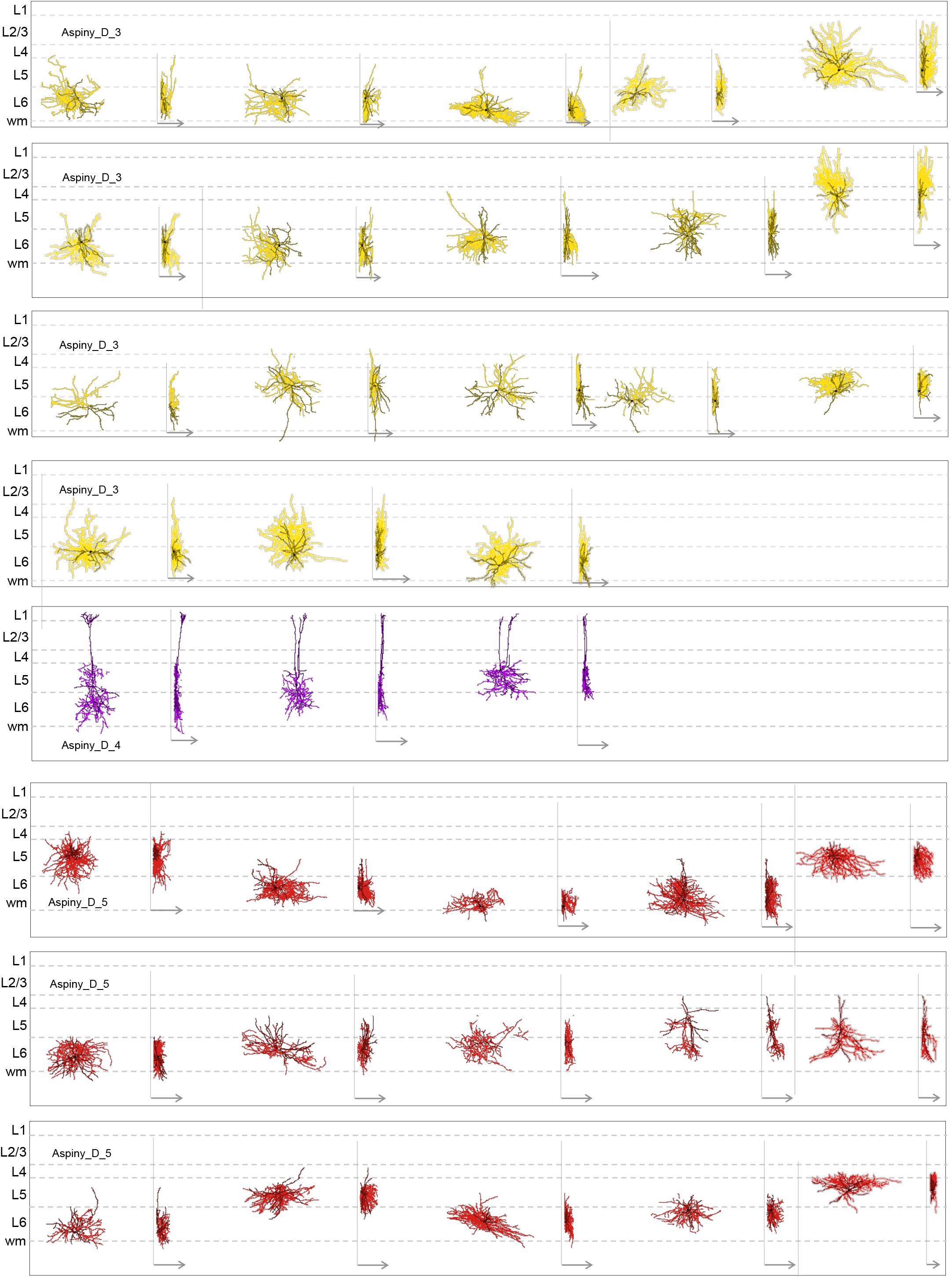
Aspiny and sparsely spiny neuron morphology. All 3D reconstructions that went into our quantitative analysis are displayed in their approximate laminar location within the cortical thickness. Two views of each reconstruction are shown. For each cell, the XY dimension view is on the left and the YZ dimension view is on the right and has an arrow indicating the Z dimension (in this case, Z is into the depth of the coronal slice). Dendrites are displayed in the darker hue and axon in the lighter hue. We reconstructed neurons with healthy, relatively intact dendrites and extensive local axon. Neurons were sampled from all cortical layers and across the major genetically and/or morphologically defined classes in mouse primary visual cortex. A wide diversity of morphologies can be observed. (N = 173).

**Supplementary Figure 20:**
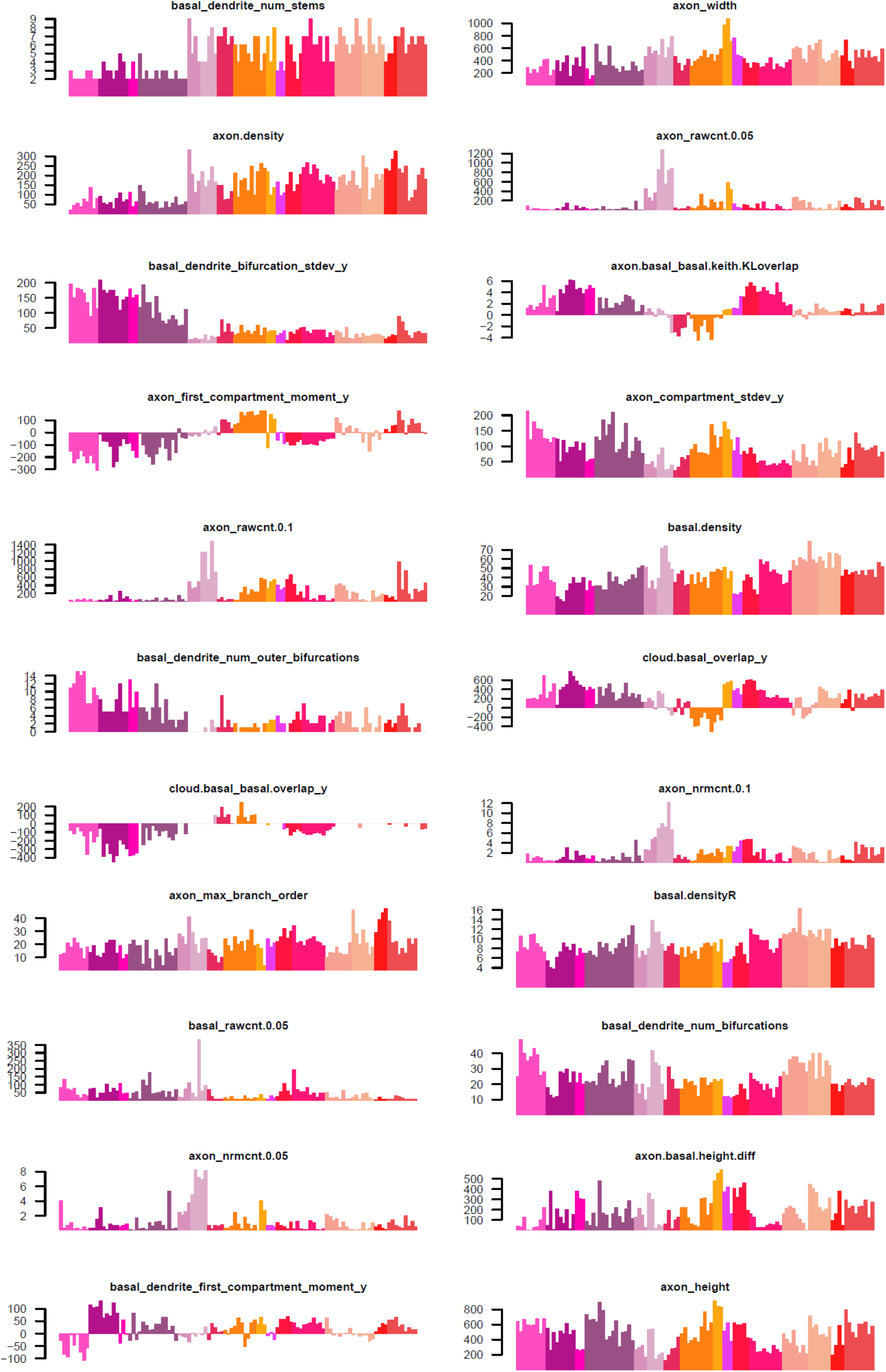
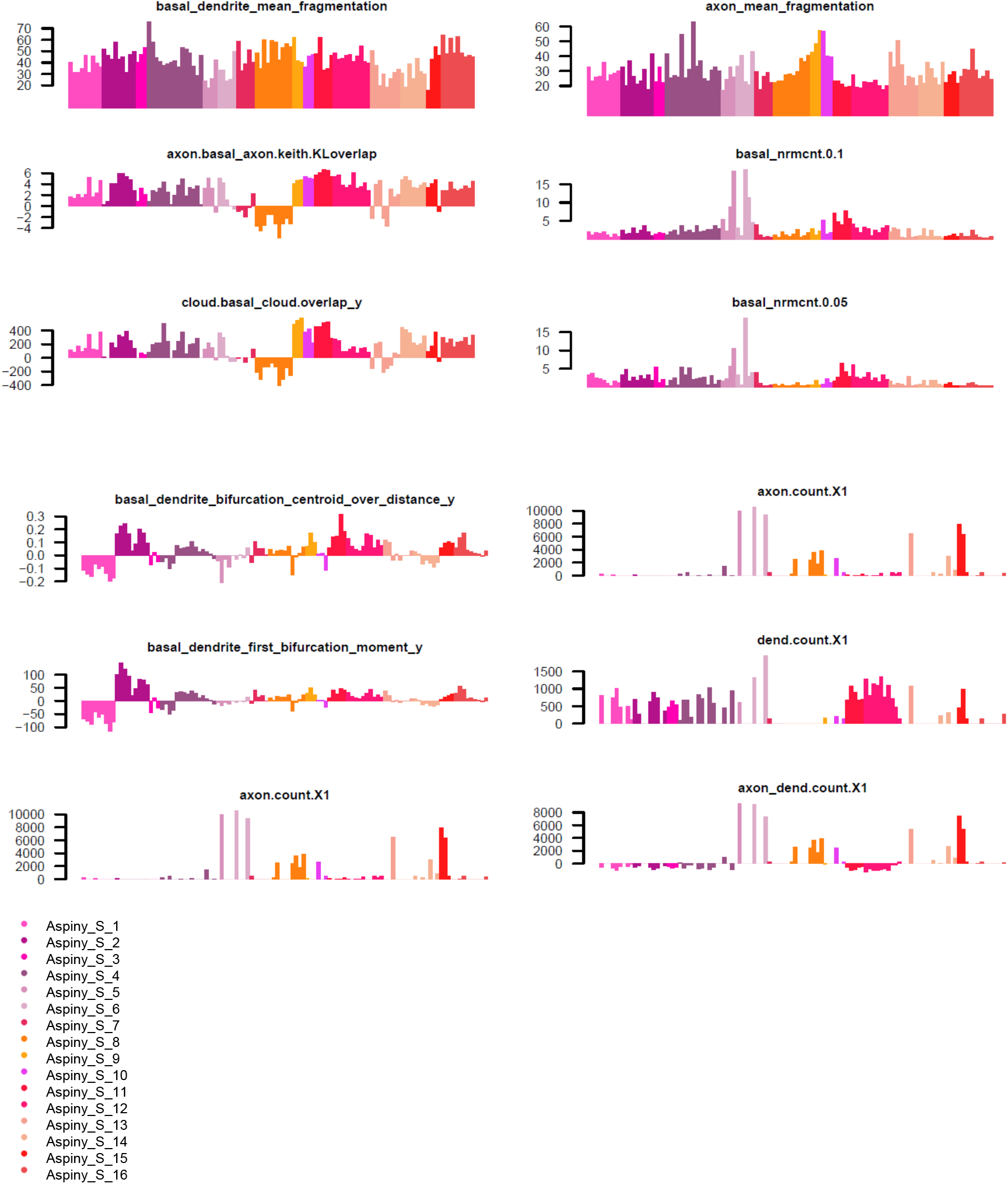
Aspiny and sparsely spiny superficial neuron morphology features by cluster. Based on 3D reconstructions of the basal dendrites and local axon, we extracted numerous morphological features from each neuron. Population histograms of 34 representative features are shown. Many of the features vary substantially across m-types (N = 109).

**Supplementary Figure 21:**
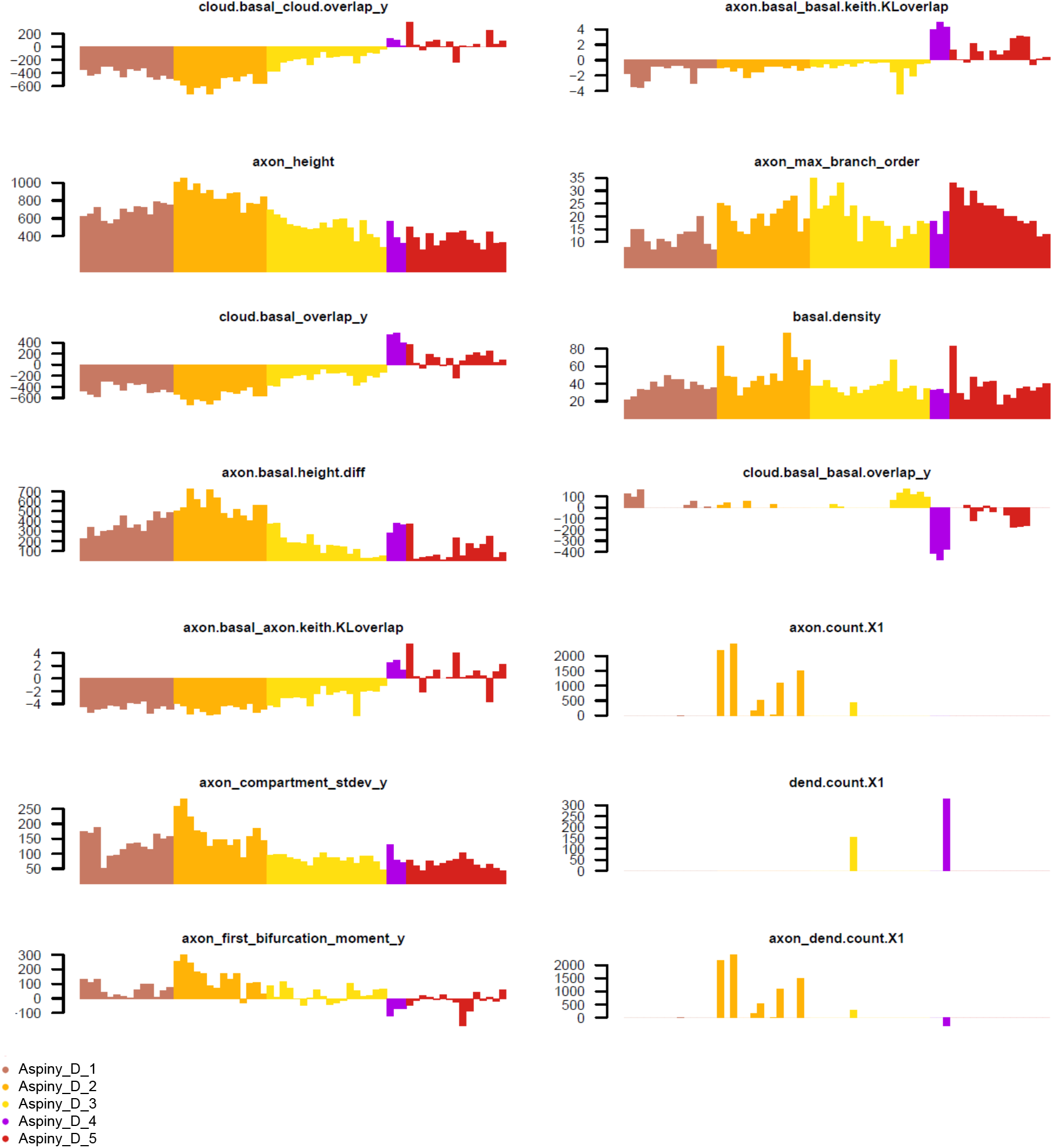
Aspiny and sparsely spiny deep neuron morphology features by cluster. Based on 3D reconstructions of the basal dendrites and local axon, we extracted numerous morphological features from each neuron. Population histograms of 14 representative features are shown. Many of the features vary substantially across m-types (N = 64).

**Supplementary Figure 22:**
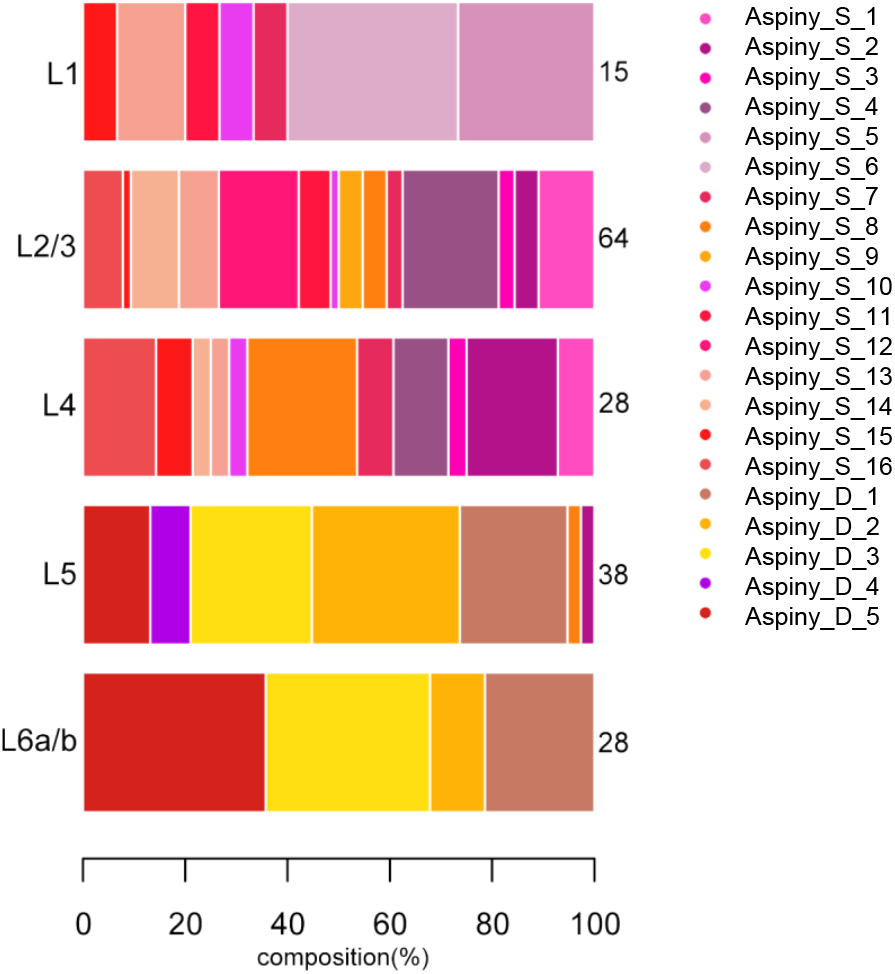
Representation of Aspiny superficial and deep neuron morphological clusters per layer. Relative distribution of the aspiny m-types across cortical layers 1–6.

**Supplementary Figure 23:**
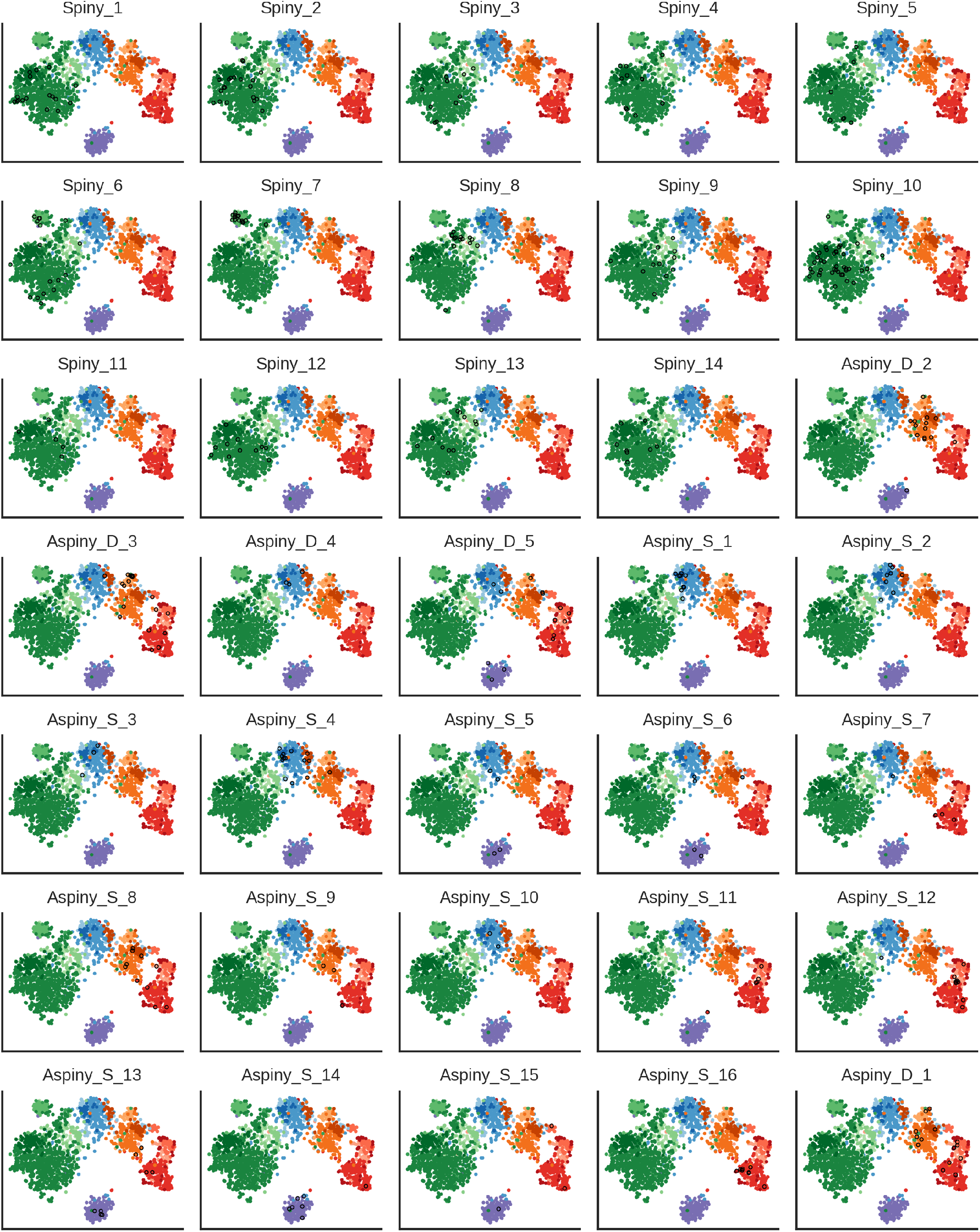
Morphological types on the electrophysiological projection. Electrophysiology-based t-SNE plots with cells from different m-types highlighted. Colors indicate e-type labels (see Fig. 2). Cells with the indicated m-type are indicated with black circles.

**Supplementary Table 1.** Electrophysiological data sets.

|  | NAME | DESCRIPTION | TYPE | SPARSE<br>COMPONENTS<br>(EXC. / INH. / ALL) |
| --- | --- | --- | --- | --- |
| 1. | AP $V_m$ | Vm of first AP from short pulse, long step, and ramp; includes 3 ms after AP threshold | Waveform | 7 / 6 / 5 |
| 2. | AP $dV/dt$ | Time derivative of (1) | Waveform | 8 / 8 / 8 |
| 3. | ISI shape | Average of ISI voltage trajectories, aligned to the threshold of the initial AP and normalized in duration | Waveform | 3 / 3 / 3 |
| 4. | Subthr. (abs.) | Concatenated responses to hyperpolarizing current steps (from -10 pA to -90 pA) | Waveform; steps from -90 pA to -10 pA | 2 / 2 / 2 |
| 5. | Subthr. (norm.) | Response to largest amplitude hyperpolarizing current step, aligned to baseline membrane potential and normalized by maximum voltage deflection | Waveform | 4 / 5 / 5 |
| 6. | PSTH | AP counts in 50 ms bins, divided by bin width | Binned (50 ms); steps from rheobase to rheobase + 100 pA | 6 / 6 / 5 |
| 7. | Inst. firing rate | Instantaneous firing rate across long steps | Binned (20 ms); steps from rheobase to rheobase + 100 pA | 6 / 5 / 5 |
| 8. | Up/down | Upstroke/downstroke ratio across long steps | Binned (20 ms); steps from rheobase to rheobase + 100 pA | 2 / 2 / 2 |
| 9. | AP peak | AP peak across long steps | Binned (20 ms); steps from rheobase to rheobase + 100 pA | 2 / 2 / 2 |
| 10. | AP fast tr. | AP fast trough across long steps | Binned (20 ms); steps from rheobase to rheobase + 100 pA | 2 / 2 / 2 |
| 11. | AP thresh. | AP threshold across long steps | Binned (20 ms); steps from rheobase to rheobase + 100 pA | 3 / 4 / 4 |
| 12. | AP width | Upstroke/downstroke ratio across long steps | Binned (20 ms); steps from rheobase to rheobase + 100 pA | 2 / 2 / 2 |
| 13. | Inst. freq. (norm.) | Instantaneous firing rate across long steps, normalized to maximum rate for each step | Binned (20 ms); steps from rheobase to rheobase + 100 pA | 6 / 7 / 7 |

**Supplementary Table 2:**
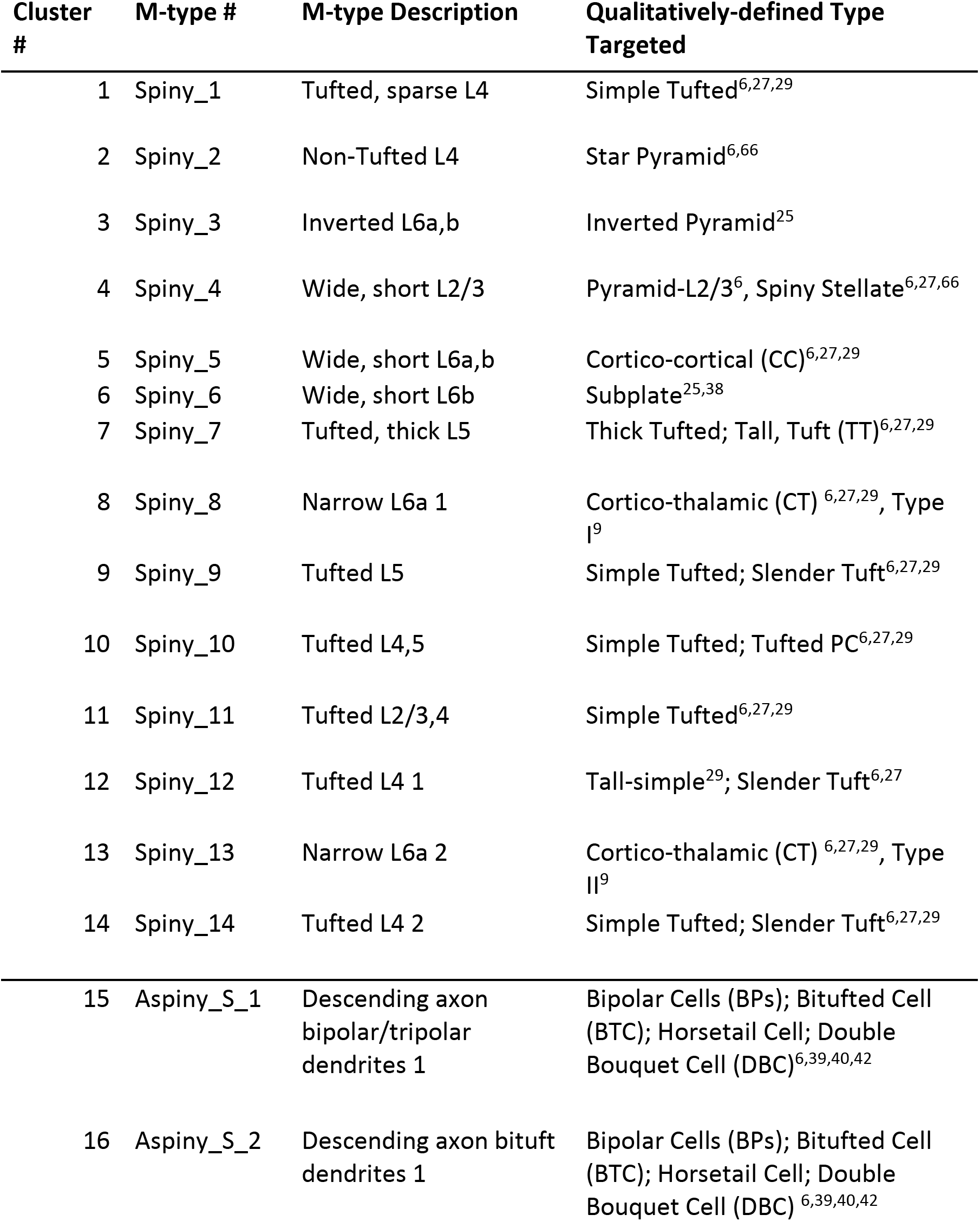
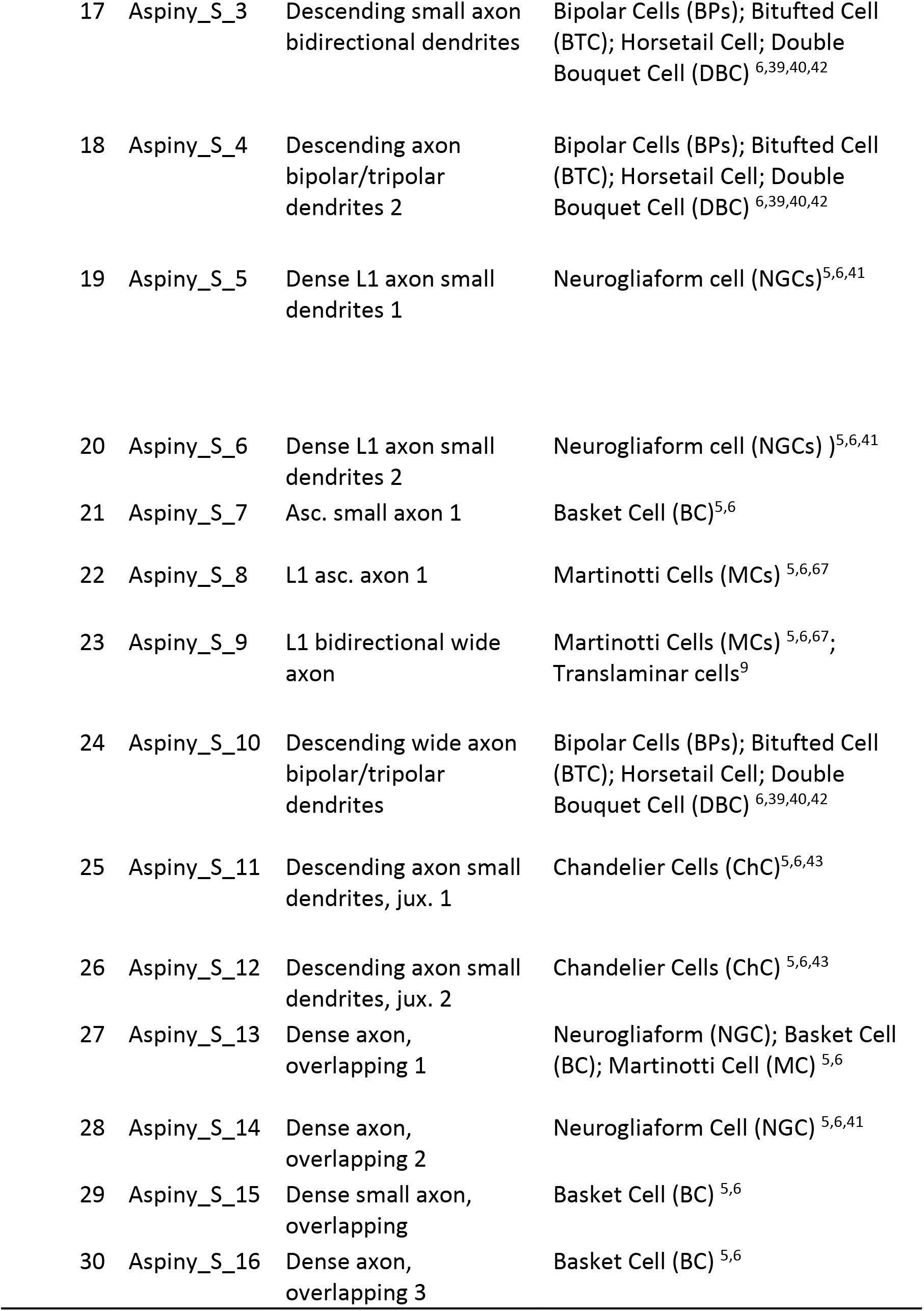
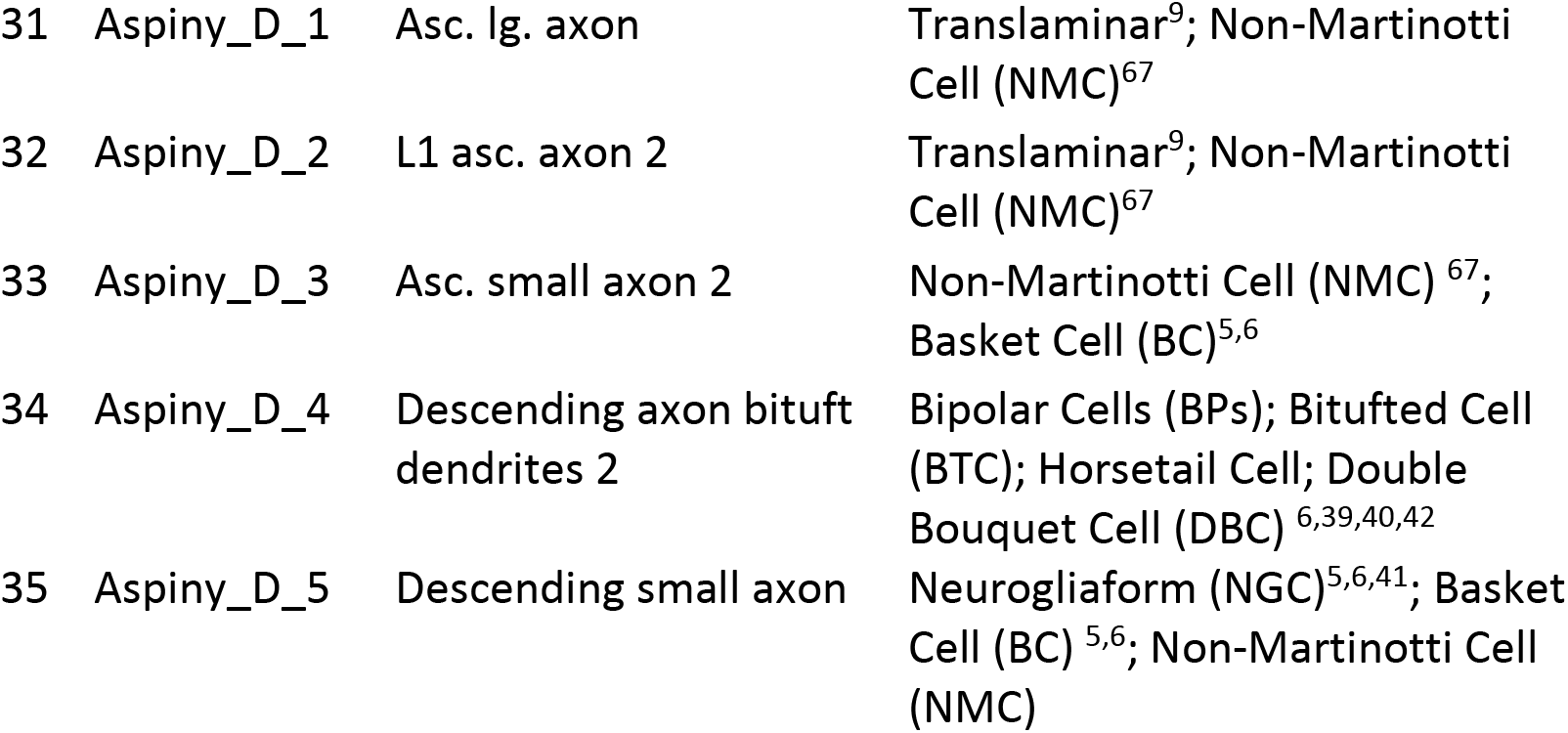
Comparison between Allen morphological types and existing literature.

| **Cluster #** | **M-type #** | **M-type Description** | **Qualitatively-defined Type Targeted** |
| --- | --- | --- | --- |
| 1 | Spiny_1 | Tufted, sparse L4 | Simple Tufted<sup>6,27,29</sup> |
| 2 | Spiny_2 | Non-Tufted L4 | Star Pyramid<sup>6,66</sup> |
| 3 | Spiny_3 | Inverted L6a,b | Inverted Pyramid<sup>25</sup> |
| 4 | Spiny_4 | Wide, short L2/3 | Pyramid-L2/3<sup>6</sup>, Spiny Stellate<sup>6,27,66</sup> |
| 5 | Spiny_5 | Wide, short L6a,b | Cortico-cortical (CC)<sup>6,27,29</sup> |
| 6 | Spiny_6 | Wide, short L6b | Subplate<sup>25,38</sup> |
| 7 | Spiny_7 | Tufted, thick L5 | Thick Tufted; Tall, Tuft (TT)<sup>6,27,29</sup> |
| 8 | Spiny_8 | Narrow L6a 1 | Cortico-thalamic (CT) <sup>6,27,29</sup>, Type I<sup>9</sup> |
| 9 | Spiny_9 | Tufted L5 | Simple Tufted; Slender Tuft<sup>6,27,29</sup> |
| 10 | Spiny_10 | Tufted L4,5 | Simple Tufted; Tufted PC<sup>6,27,29</sup> |
| 11 | Spiny_11 | Tufted L2/3,4 | Simple Tufted<sup>6,27,29</sup> |
| 12 | Spiny_12 | Tufted L4 1 | Tall-simple<sup>29</sup>; Slender Tuft<sup>6,27</sup> |
| 13 | Spiny_13 | Narrow L6a 2 | Cortico-thalamic (CT) <sup>6,27,29</sup>, Type II<sup>9</sup> |
| 14 | Spiny_14 | Tufted L4 2 | Simple Tufted; Slender Tuft<sup>6,27,29</sup> |
| 15 | Aspiny_S_1 | Descending axon bipolar/tripolar dendrites 1 | Bipolar Cells (BPs); Bitufted Cell (BTC); Horsetail Cell; Double Bouquet Cell (DBC)<sup>6,39,40,42</sup> |
| 16 | Aspiny_S_2 | Descending axon bituft dendrites 1 | Bipolar Cells (BPs); Bitufted Cell (BTC); Horsetail Cell; Double Bouquet Cell (DBC) <sup>6,39,40,42</sup> |
| 17 | Aspiny_S_3 | Descending small axon<br>bidirectional dendrites | Bipolar Cells (BPs); Bitufted Cell<br>(BTC); Horsetail Cell; Double<br>Bouquet Cell (DBC) <sup>6,39,40,42</sup> |
| 18 | Aspiny_S_4 | Descending axon<br>bipolar/tripolar<br>dendrites 2 | Bipolar Cells (BPs); Bitufted Cell<br>(BTC); Horsetail Cell; Double<br>Bouquet Cell (DBC) <sup>6,39,40,42</sup> |
| 19 | Aspiny_S_5 | Dense L1 axon small<br>dendrites 1 | Neurogliaform cell (NGCs) <sup>5,6,41</sup> |
| 20 | Aspiny_S_6 | Dense L1 axon small<br>dendrites 2 | Neurogliaform cell (NGCs) <sup>5,6,41</sup> |
| 21 | Aspiny_S_7 | Asc. small axon 1 | Basket Cell (BC) <sup>5,6</sup> |
| 22 | Aspiny_S_8 | L1 asc. axon 1 | Martinotti Cells (MCs) <sup>5,6,67</sup> |
| 23 | Aspiny_S_9 | L1 bidirectional wide<br>axon | Martinotti Cells (MCs) <sup>5,6,67</sup>;<br>Translaminar cells<sup>9</sup> |
| 24 | Aspiny_S_10 | Descending wide axon<br>bipolar/tripolar<br>dendrites | Bipolar Cells (BPs); Bitufted Cell<br>(BTC); Horsetail Cell; Double<br>Bouquet Cell (DBC) <sup>6,39,40,42</sup> |
| 25 | Aspiny_S_11 | Descending axon small<br>dendrites, jux. 1 | Chandelier Cells (ChC) <sup>5,6,43</sup> |
| 26 | Aspiny_S_12 | Descending axon small<br>dendrites, jux. 2 | Chandelier Cells (ChC) <sup>5,6,43</sup> |
| 27 | Aspiny_S_13 | Dense axon,<br>overlapping 1 | Neurogliaform (NGC); Basket Cell<br>(BC); Martinotti Cell (MC) <sup>5,6</sup> |
| 28 | Aspiny_S_14 | Dense axon,<br>overlapping 2 | Neurogliaform Cell (NGC) <sup>5,6,41</sup> |
| 29 | Aspiny_S_15 | Dense small axon,<br>overlapping | Basket Cell (BC) <sup>5,6</sup> |
| 30 | Aspiny_S_16 | Dense axon,<br>overlapping 3 | Basket Cell (BC) <sup>5,6</sup> |
| 31 | Aspiny_D_1 | Asc. lg. axon | Translaminar<sup>9</sup>; Non-Martinotti Cell (NMC)<sup>67</sup> |
| 32 | Aspiny_D_2 | L1 asc. axon 2 | Translaminar<sup>9</sup>; Non-Martinotti Cell (NMC)<sup>67</sup> |
| 33 | Aspiny_D_3 | Asc. small axon 2 | Non-Martinotti Cell (NMC)<sup>67</sup>; Basket Cell (BC)<sup>5,6</sup> |
| 34 | Aspiny_D_4 | Descending axon bituft dendrites 2 | Bipolar Cells (BPs); Bitufted Cell (BTC); Horsetail Cell; Double Bouquet Cell (DBC)<sup>6,39,40,42</sup> |
| 35 | Aspiny_D_5 | Descending small axon | Neurogliaform (NGC)<sup>5,6,41</sup>; Basket Cell (BC)<sup>5,6</sup>; Non-Martinotti Cell (NMC) |

**Supplementary Table 3:**
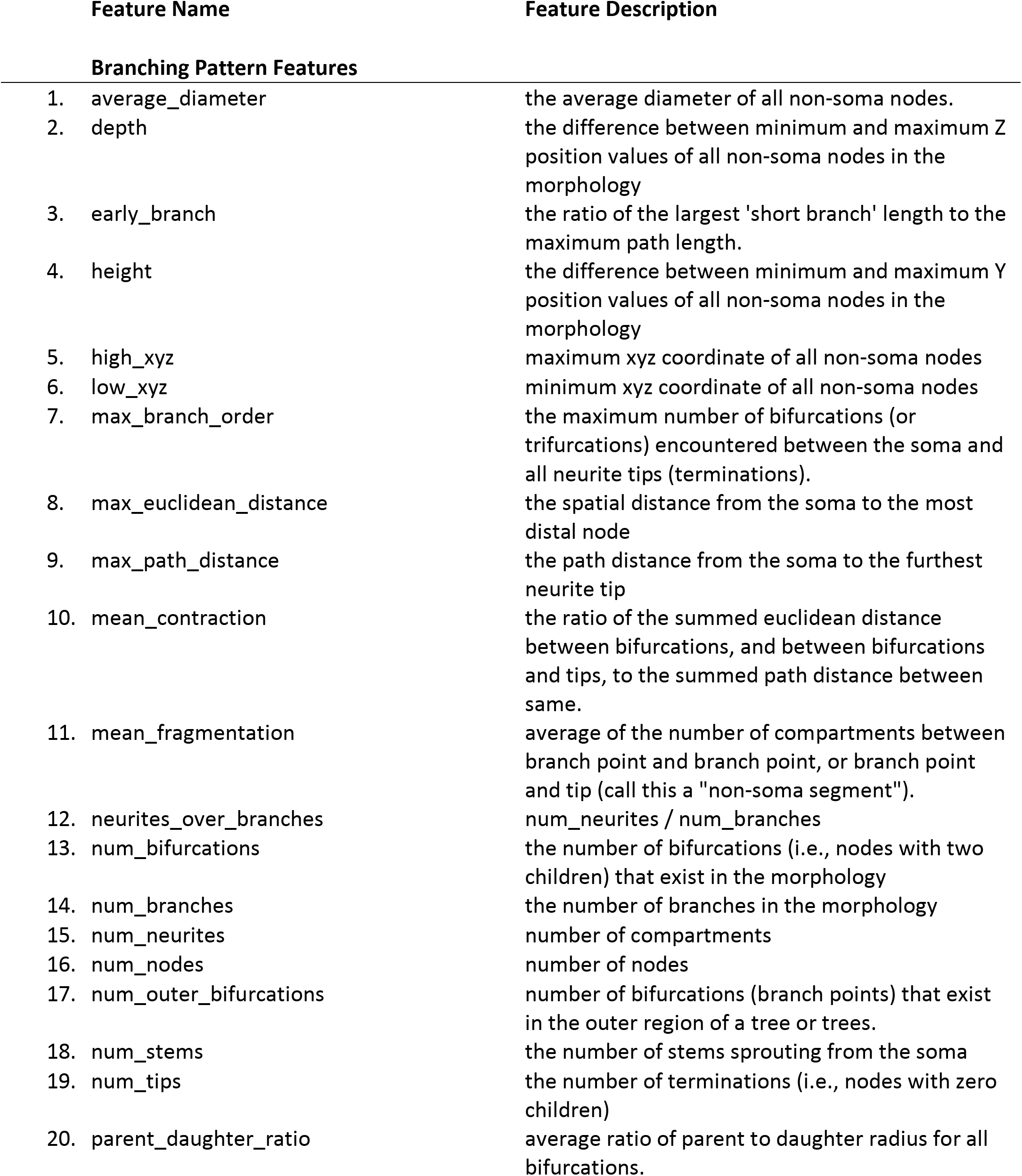
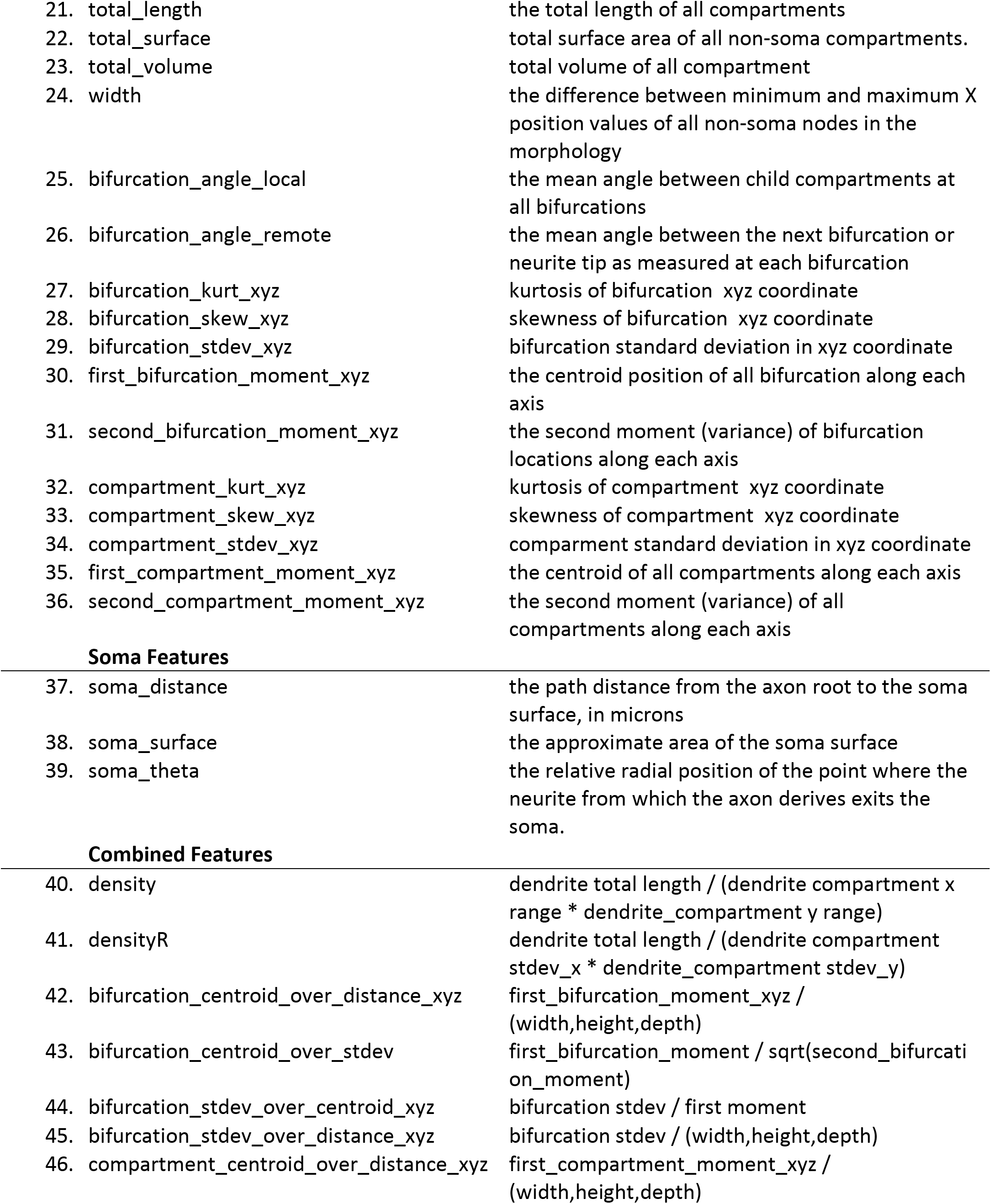
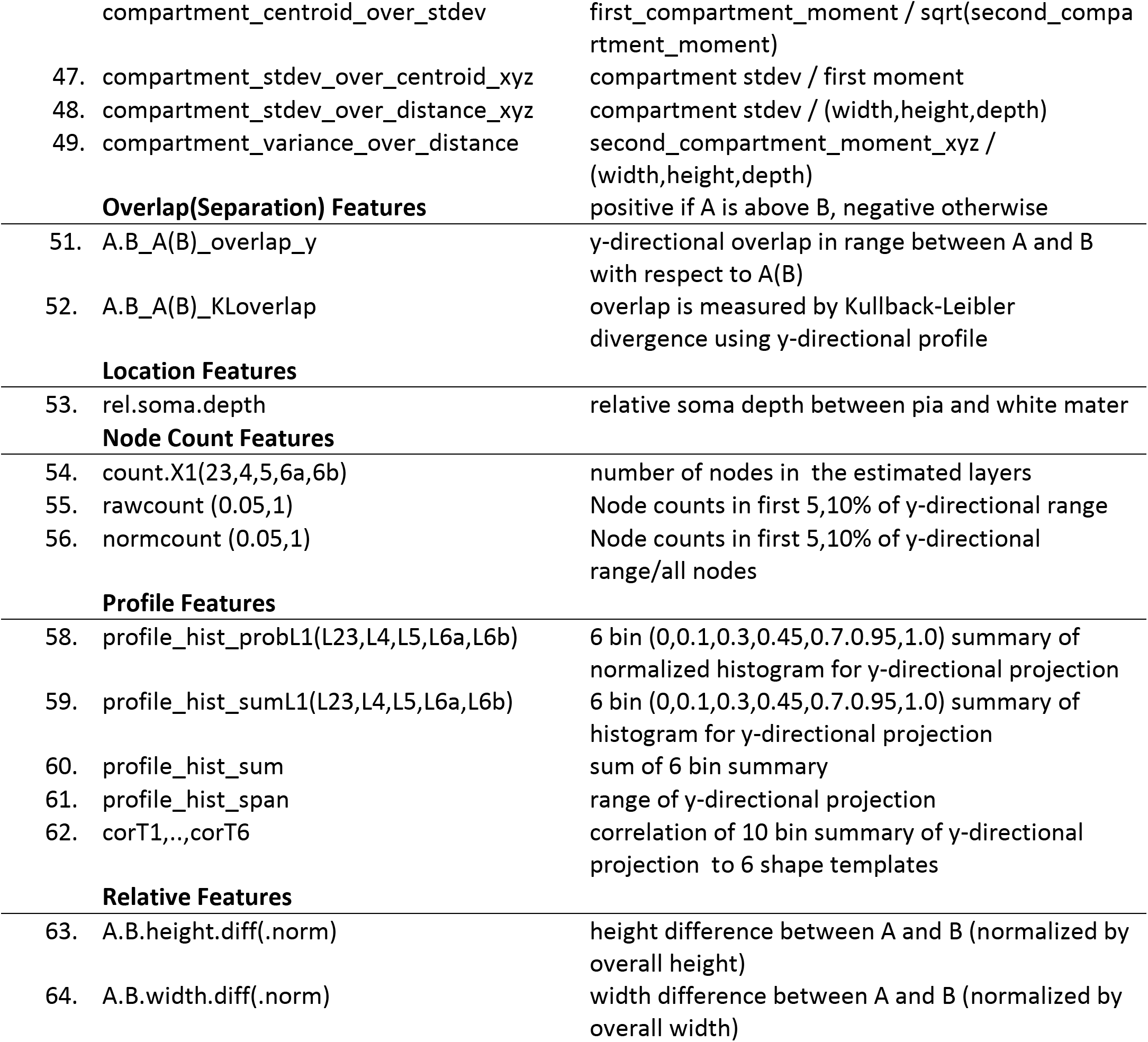
Description of morphological features.

**Supplementary Table 4.**
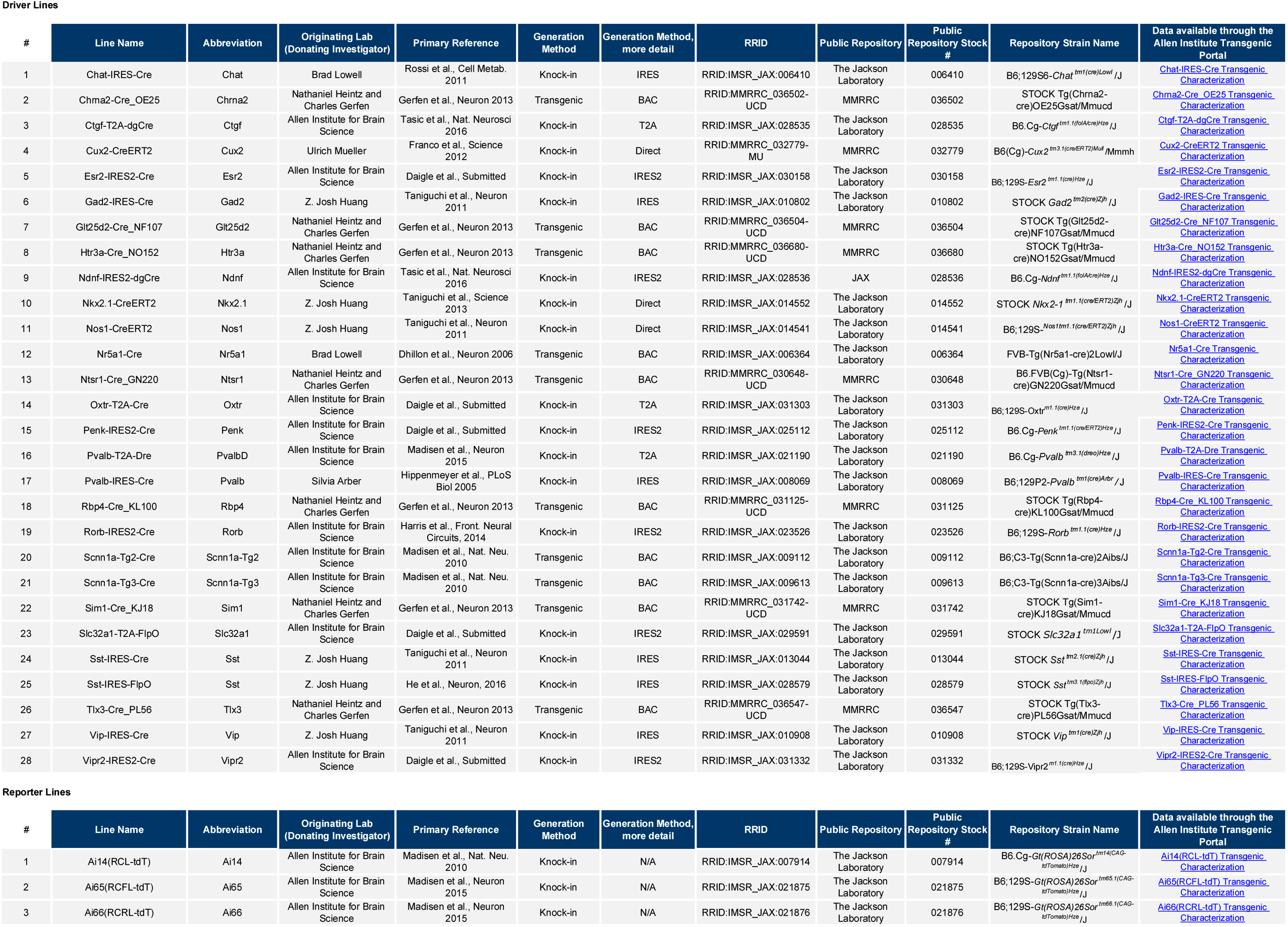

